# Direct monitoring of the thermodynamics and kinetics of DNA and RNA dinucleotide dehybridization from gaps and overhangs

**DOI:** 10.1101/2023.04.10.536266

**Authors:** Brennan Ashwood, Michael S. Jones, Aleksandar Radakovic, Smayan Khanna, Yumin Lee, Joseph R. Sachleben, Jack W. Szostak, Andrew L. Ferguson, Andrei Tokmakoff

## Abstract

Hybridization of short nucleic acid segments (<4 nucleotides) to single-strand templates occurs as a critical intermediate in processes such as non-enzymatic nucleic acid replication and toehold-mediated strand displacement. These templates often contain adjacent duplex segments that stabilize base pairing with single-strand gaps or overhangs, but the thermodynamics and kinetics of hybridization in such contexts are poorly understood due to experimental challenges of probing weak binding and rapid structural dynamics. Here we develop an approach to directly measure the thermodynamics and kinetics of DNA and RNA dinucleotide dehybridization using steady-state and temperature-jump infrared spectroscopy. Our results suggest that dinucleotide binding is stabilized through coaxial stacking interactions with the adjacent duplex segments as well as from potential non-canonical base pairing configurations and structural dynamics of gap and overhang templates revealed using molecular dynamics simulations. We measure timescales for dissociation ranging from 0.2 to 40 µs depending on the template and temperature. Dinucleotide hybridization and dehybridization involves a significant free energy barrier with characteristics resembling that of canonical oligonucleotides. Together, our work provides an initial step for predicting the stability and kinetics of hybridization between short nucleic acid segments and various templates.

## Introduction

DNA and RNA duplex hybridization has been investigated for more than sixty years with an aim of developing predictive models for how thermodynamic and kinetic properties will vary with molecular factors such as strand length, sequence, and chemical modifications.^1–3^ The development of numerous models, such as the quantitatively accurate nearest-neighbor (NN) models^4–5^ for thermodynamics or the kinetic-zipper model,^6–7^ shape the modern understanding of nucleic acid properties as well as the development of nucleic acid technology.^8–9^ However, most of these models were developed for oligonucleotides, and the hybridization properties of nucleic acid segments shorter than six nucleotides in length have largely been neglected. Hybridization of such short segments is often perceived as irrelevant due to their poor binding stability in aqueous solution, yet there are biological processes where segments as short as mononucleotides must hybridize. Many of these processes, such as the binding of 2′-deoxynucleotide triphosphates to single-strand DNA (ssDNA) during strand replication or binding between three nucleotide anticodons of tRNA and their complement mRNA, utilize additional protein-nucleic acid interactions to promote binding.^10–11^ However, processes such as non-enzymatic replication of nucleic acids and toehold-mediated strand displacement require hybridization of 1-4 base pair patches without the help of proteins.^8, 12–14^

A conserved feature among the examples of short oligonucleotide hybridization is that a short segment binds to a single-strand overhang adjacent to a duplex region or in a gap between two duplex segments. The adjacent duplexes offer coaxial stacking interactions that stabilize hybridization of the short oligonucleotide. The thermodynamic benefit of coaxial stacking has been measured for DNA and RNA oligonucleotides and is often found to be similar to stacking in B-DNA or A-RNA, respectively.^15–20^ The structural constraints of the sugar-phosphate backbone at a nick site are relaxed relative to a covalently-linked base pair step and may enable more stabilizing stacking configurations.^21–22^ Additionally, the single-strand segments in overhang and gap templates may have different physical properties from free single-strands such as reduced configurational flexibility, increased stacking, and different hydration that influence the stability and dynamics of binding with short oligonucleotides. A few studies have shown evidence for increased rigidity of gap and overhang single-stranded regions relative to free single-strands as well as bending of duplexes containing 1 or 2 nucleotide gaps,^23–29^ and these properties are likely highly sensitive to nucleobase sequence and cation concentrations.^30–32^

The binding stability of mononucleotides with overhang and gap templates has recently been studied. NMR-monitored titrations of mononucleotides with overhangs revealed dissociation constants (*K*_d_) ranging from ∼10 mM for G:C base pairing to ∼200 mM for A:T base pairing.^33–34^ Isothermal titration calorimetry (ITC) and FRET-monitored titrations have shown that the *K*_d_ for binding onto a gap decreases nearly 3 orders of magnitude to values of 0.2-0.6 mM,^35–36^ which corresponds to a ∼12 kJ mol^-1^ increase in the dissociation free energy 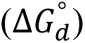 and is consistent with forming an additional nearest-neighbor step.^4, 37^ These studies give a sense for the stability of short oligonucleotide hybridization to overhangs and gaps and indicate that it depends on sequence and whether DNA or RNA is used, yet, with the exception of ITC, the thermodynamic information is limited to dissociation constants (*K*_d_ or 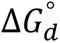) at the measurement temperature. A detailed understanding of hybridization onto overhangs and gaps requires knowledge of enthalpic and entropic contributions to stability and the role of relevant environmental variables such as temperature and counterion concentration.^38–40^ More importantly, no studies have directly monitored the time-dependence of hybridization and dehybridization for short oligonucleotides onto templates, and these timescales determine whether reactions such as non-enzymatic extension or toehold-mediated strand displacement are possible.

Here we report the temperature-dependent thermodynamics and kinetics of short oligonucleotide dehybridization from overhangs and gaps using steady-state and temperature-jump infrared (T-jump IR) spectroscopy. IR spectroscopy is sensitive to base pairing and stacking interactions and distinctly resolves changes in A:T and G:C base pairing without the need for synthetic labels.^41–42^ Characterization of temperature-dependent binding stability enables accurate determination of enthalpic and entropic contributions and is particularly relevant in non-enzymatic replication where temperature variations may play a key role.^38–39^ We focus on the binding of DNA and RNA dinucleotides and demonstrate our approach with a model system of an adenine-adenine (AA) dinucleotide binding next to pure G:C duplex regions. This model system was chosen to maximize spectral contrast between binding of the dinucleotide and the structural changes and dynamics within the template itself, but our approach is applicable for any nucleobase sequence. We find quantitative agreement between thermodynamic results obtained from IR spectroscopy and those from complementary temperature-dependent ^1^H NMR and ITC measurements of binding. From T-jump IR measurements, we extract a time constant ranging from 200 ns to 40 µs for dissociation of AA from the template that depends on the temperature and template, while the time constant for association is a few microseconds and shows a minor dependence on temperature. We also study the binding of 3-mer (AAA) and 4-mer (AAAA) adenine DNA oligonucleotides to assess the length-dependence of association in this short oligonucleotide regime. Our experimental results as well as all-atom molecular dynamics (MD) simulations suggest that differences in the structure and dynamics of the bound complex and template compared to free oligonucleotides lead to significantly different binding thermodynamics and kinetics for overhangs and gaps.

## Materials and Methods

### Synthesis and purification of RNA oligonucleotides

RNA oligonucleotides were synthesized in house on a K&A H-6 solid-phase synthesizer. Phosphoramidites and reagents for the K&A synthesizer were purchased from Glen Research and Chemgenes. 1.5 mL of a 1:1 mixture of 28 % aqueous ammonium hydroxide and 40 % aqueous methylamine was used to cleave the RNA from the solid support for 15 minutes at room temperature, and was followed by deprotection in the same solution for 15 minutes at 65 °C. After cooling, the solutions were evaporated for 1 hour in a SpeedVac vacuum concentrator and lyophilized to dryness. The dried 2′-O-tertbutyldimethylsilyl (2′-O-TBDMS) protected RNA was dissolved in 115 µL DMSO, 60 µL triethylamine, and 75 µL triethylamine trihydrofluoride and deprotected for 2.5 hours at 65 °C. After cooling, the RNA was purified by two different methods depending on its length:

Short (≤ 6 nucleotides) oligonucleotides were purified by a modified Glen-Pak protocol (Glen Research). After the RNA was quenched, bound to the resin, and washed to remove failed sequences, it was eluted with 50 % acetonitrile/water v/v. Following evaporation to dryness, the RNA was dissolved in 500 µL 2 % trifluoroacetic acid to deprotect the 5′-O-dimethoxytitryl (5′-O-DMT) for 5 minutes at room temperature. The deprotection reaction was quenched with 500 µL 1 M triethylammonium bicarbonate pH 9 and desalted using Sep-Pak C18 cartridges (Waters).

Longer (> 7 nucleotides) oligonucleotides were precipitated with 25 µL 5 M ammonium acetate and 1 mL 2-propanol at −78 °C for 20 minutes, washed with 1 mL 80 % ethanol in water v/v, air-dried, dissolved in 5 mM EDTA, 99 % formamide in water v/v, and purified by preparative 20 % polyacrylamide gel electrophoresis (19:1 with 7 M urea). After crushing and soaking the desired gel band in 5 mM sodium acetate, 2 mM EDTA for 16 hours at room temperature, the extracted RNA was desalted using Sep-Pak C18 cartridges (Waters).

Purified RNA was characterized by high-resolution mass spectrometry on an Agilent 6540 Q-TOF mass spectrometer.

### Preparation of DNA oligonucleotides

DNA oligonucleotides were purchased from Integrated DNA Technologies (IDT) at desalt grade purity. Short (≤ 6 nucleotides) oligonucleotides were further desalted using Sep-Pak C18 cartridges (Waters), evaporated for 1 hour in a SpeedVac vacuum concentrator, and lyophilized to dryness. Longer (> 7 nucleotides) oligonucleotides were dialyzed in ∼1.5 L ultrapure water for 24 h using Slide-A-Lyzer cassettes (2 kDa cutoff, Thermo Scientific), where the water was replaced every ∼6 hrs, and then lyophilized to dryness. For all measurements, oligonucleotides were prepared in deuterated pH* 6.8 400 mM sodium phosphate buffer (SPB, [Na^+^] = 600 mM). Oligonucleotide concentration was verified with UV absorbance using a NanoDrop UV/Vis spectrometer (Thermo Scientific).

### FTIR spectroscopy

FTIR spectra were measured with a Bruker Tensor 27 FTIR spectrometer at 2 cm^-1^ resolution. 30 µL of sample was placed between two 1 mm thick CaF_2_ windows with a pathlength defined by a 50 µm Teflon spacer. The windows were enclosed in brass jacket that is temperature-controlled using a recirculating chiller (Ministat 125, Huber). The temperature at the sample was measured with a thermocouple in thermal contact to one CaF_2_ window for chiller set points from −3 to 105 °C, which was used to calibrate the reported temperatures in this work.

Prior to titration experiments performed at 1 °C, the gap or overhang template solution was incubated at 85 °C for 2 min and cooled to room temperature under ambient conditions. Samples were prepared with an AA-to-template molar ratio ranging from 0 to 4 with a constant template concentration of 1 mM. Samples were incubated at 1 °C for 10 min prior to each measurement. For FTIR temperature series, all oligonucleotides were prepared at a 1 mM concentration and annealed prior to measurements as for titration measurements. Temperature series were performed in ∼2.6 °C steps and the sample was equilibrated for 3 min at each temperature prior to acquiring spectra. A discrete wavelet transform using the Mallat algorithm and symlet family was applied to the 1490 – 1750 cm^-1^ region of the FTIR spectra to separate and subtract the D_2_O background absorption from each spectrum.^43^

### Two-dimensional IR Spectroscopy

Two-dimensional IR (2D IR) measurements were performed using a previously described setup with a pump-probe beam geometry.^44^ Spectra were collected using parallel pulse polarization (ZZZZ) at a fixed waiting time (t_2_) of 150 fs. The pump pulse pair delay (t_1_) was scanned from −160 to 1900 fs in 16 fs steps. Samples were prepared identically as for FTIR measurements. Sample temperature was controlled using a recirculating chiller (Ministat 125, Huber). Temperature-dependent drift of the sample out of the pump and probe focus and overlap region was minimized by programmatically maximizing the pump-probe signal at each temperature using a motorized translation stage prior to measuring the 2D IR data.

### Temperature-jump IR Spectroscopy

The T-jump IR spectrometer used in this work has been described previously.^45–46^ Optical heating increases the sample temperature within 7 ns and the ensuing changes in base pairing and stacking of DNA and RNA are monitored with heterodyned dispersed vibrational echo spectroscopy (HDVE)^47^ from nanosecond to millisecond time delays. The HDVE spectrum is collected with parallel pulse polarization (ZZZZ) and at a fixed waiting time (t_2_) of 150 fs and reports on the nucleobase ring and carbonyl vibrational bands similarly to an IR pump-probe spectrum. The T-jump laser power was attenuated with a polarizer to set a T-jump magnitude (*ΔT* = *T*_*f*_ − *T*_*i*_) of ∼13 °C for all measurements. The initial temperatures (*T*_*i*_) were set using a recirculating chiller connected to a brass sample jacket and ΔT was determined from the change in mid-IR transmission following the T-jump pulse.

Given the complex form of the T-jump relaxation kinetics spanning multiple time scales, we chose a model-free analysis method to extract experimental relaxation rates. The time-domain t-HDVE response at different IR frequencies was converted into a rate spectrum using an inverse Laplace transform that employs a maximum entropy regularization method (MEM-iLT) which is described elsewhere.^48–49^ We calculate the observed relaxation rate, *k*_*obs*_, for a given process by calculating the amplitude-weighted mean value of the rate for a corresponding peak in the rate spectrum.

### Isothermal Titration Calorimetry

Isothermal titration calorimetry (ITC) measurements were performed using a MicroCal iTC200 (Malvern Panalytical). Template oligonucleotides were placed in the cell with a concentration that varied from 60 μM to 3 mM depending on the sequence and the complement short oligonucleotide was placed in the syringe with a concentration of 100-300 μM. Sample conditions for each measurement are listed in Table S2. All samples were prepared in deuterated pH 6.8 400 mM SPB and degassed under vacuum for >20 min at the experimental temperature prior to each measurement. ITC measurements started with an initial 0.4 μL injection followed by 19 injections of 2 μL aliquots of the titrant oligonucleotide solution into the cell. The syringe needle constantly stirred the cell solution with a spin rate of 1000 rpm, and the injection duration and time interval between injections were adjusted to avoid signal saturation in high concentration samples. Injection settings for each sample are listed in Table S2.

### NMR Spectroscopy

NMR spectra were acquired on a Bruker AVANCE III 500 MHz spectrometer equipped with a Bruker TXI probe. Temperature series were performed in 2.5 °C steps and the sample was equilibrated for 5 min and auto gradient and lock shimmed with TopShim at each temperature. prior to acquiring spectra. Samples were prepared in fully deuterated 400 mM SPB for all measurements and contained ∼1 mM 3-(Trimethylsilyl)propionic-2,2,3,3-d_4_ acid (Sigma-Aldrich) as a frequency reference.

Total correlation spectroscopy (TOCSY) measurements were measured with DIPSI II isotropic mixing and a mixing time of 60 ms. 2048 and 256 complex points were acquired in t2 and t1, respectively, over sweep widths of 24 ppm.

Diffusion-ordered spectroscopy (DOSY) was performed on a Bruker AVANCE IIIHD 600 MHz spectrometer. Measurements were performed using 2D stimulated echo pulse sequence with bipolar gradients and WATERGATE solvent suppression (Bruker TopSpin, stebpgp1s19). Spectra were acquired from 0 to 95% of the maximum gradient field strength in 5 % intervals where the maximum field strength was 5.35 T/cm. The magnetic field gradient was calibrated with a 3D printed phantom with known spacing between the water samples.

### Molecular dynamics simulations

DNA and RNA duplex topologies were constructed using AMBERTools22^50^ in canonical B and A forms, respectively. AA-gap complexes were generated from the canonical topologies by removing phosphate atoms after the 6^th^ (5′-**G**A-3′) and 8^th^ position (5′-**A**G-3′) nucleotides and adding terminal 5′ and 3′ hydrogens using the GROMACS molecular modeling suite.^51^ Gap templates were generated by entirely removing the A nucleotides from the canonical topologies. MD simulations were performed with the force fields that are currently most accurate for modelling canonical DNA and RNA duplex structure: The bsc1-AMBER force field^52^ with TIP3P^53^ water model for DNA systems, and the DES-AMBER RNA force field^54^ with TIP4P water model^55^ for RNA systems. To enable more direct comparisons between DNA and RNA simulation results, we performed additional simulations of DNA using the DES-AMBER force field for DNA^56^ with TIP4P water model and simulations of RNA using the bsc1-AMBER force field with TIP3P water model (Section S5). Cubic simulation boxes with periodic boundaries were constructed to ensure a minimum of 1 nm of spacing between any nucleic acid atoms and the boundaries. For 14-mer DNA this corresponded to a box with 7.16 nm side lengths with 11770 water molecules, and for 14-mer RNA a box with 6.77 nm side lengths with 9845 water molecules. NaCl ion pairs were added to the simulation box to maintain an ionic concentration of 600 mM, and additional Na^+^ ions were added to balance the negatively charged phosphate backbone.

Energy minimization was performed on each topology using the steepest descent algorithm until the maximum force was below 1000 kJ mol^-1^ nm^-1^. Equilibration was performed for 100 ps in the NVT ensemble and then 100 ps in the NPT ensemble. Production runs were performed for up to 4 µs in the NVT ensemble. Simulations were propagated with a 2 fs time step using a leap-frog integrator, and frames were saved every 200 ps. Simulation temperature was set to 310 K and maintained by a velocity-rescaling thermostat.^57^ In the NPT equilibration runs, pressure was set to 1 bar and maintained using a Parinello-Rahmen barostat. Particle Mesh Ewald^58^ was used to calculate long-range electrostatic interactions, employing a 1.0 nm real space cutoff and a 0.16 Fourier grid spacing that was optimized during runtime, and constraints were placed on hydrogen bonds using the LINCS algorithm.^59^

Cambridge conventional helical parameters^60^ and backbone torsions were calculated from MD trajectories using a Biobb BioExcel Building Blocks workflow^61–62^, which leverages the Curves+ program and its associated Canal tool^63–64^. Intra-molecular residue distances and solvent distributions were calculated using the MDTraj Python library. Additional statistical analyses and visualizations were generated using the PyEMMA and Seaborn Python libraries.^65–66^

## Results

### Low-temperature hybridization of dinucleotides to overhangs and gaps

Figure 1a illustrates the model sequences that we designed to study the hybridization of a 2′-deoxyadenosine dinucleotide onto G:C rich DNA templates incorporating a single-stranded thymine overhang or thymine gap. Overhang templates (TTo) are composed of a 6-mer primer strand bound to the 3′ end of a 14-mer template strand. Gap templates also have a 6-mer helper strand bound to the 5′-end of the template. Additionally, we studied the binding of adenosine dinucleotide onto a single-stranded uracil RNA gap template (UUg). The abbreviation AA will be used to refer to both the 2′-deoxyadenosine dinucleotide for DNA and adenosine dinucleotide for RNA samples throughout. The primer and helper segments consist of six guanine:cytosine (G:C) base pairs in order to maximize their binding stability while also keeping their size relatively small to minimize IR absorption from the template itself. The hybridized product between the template and AA is termed the AA-overhang (TTo:AA) or AA-gap complex (TTg:AA, UUg:AA). Although formation of a fully formed gap or overhang complex can potentially involve numerous binding equilibria, the choice of a weakly bound AA to a far more stable GC-rich template means that the dissociation of the complex is well described through two equilibrium constants: *K*_d_ for the dissociation of AA from the template and *K*_d,Temp_ for dissociation of the template.

**Figure 1.**
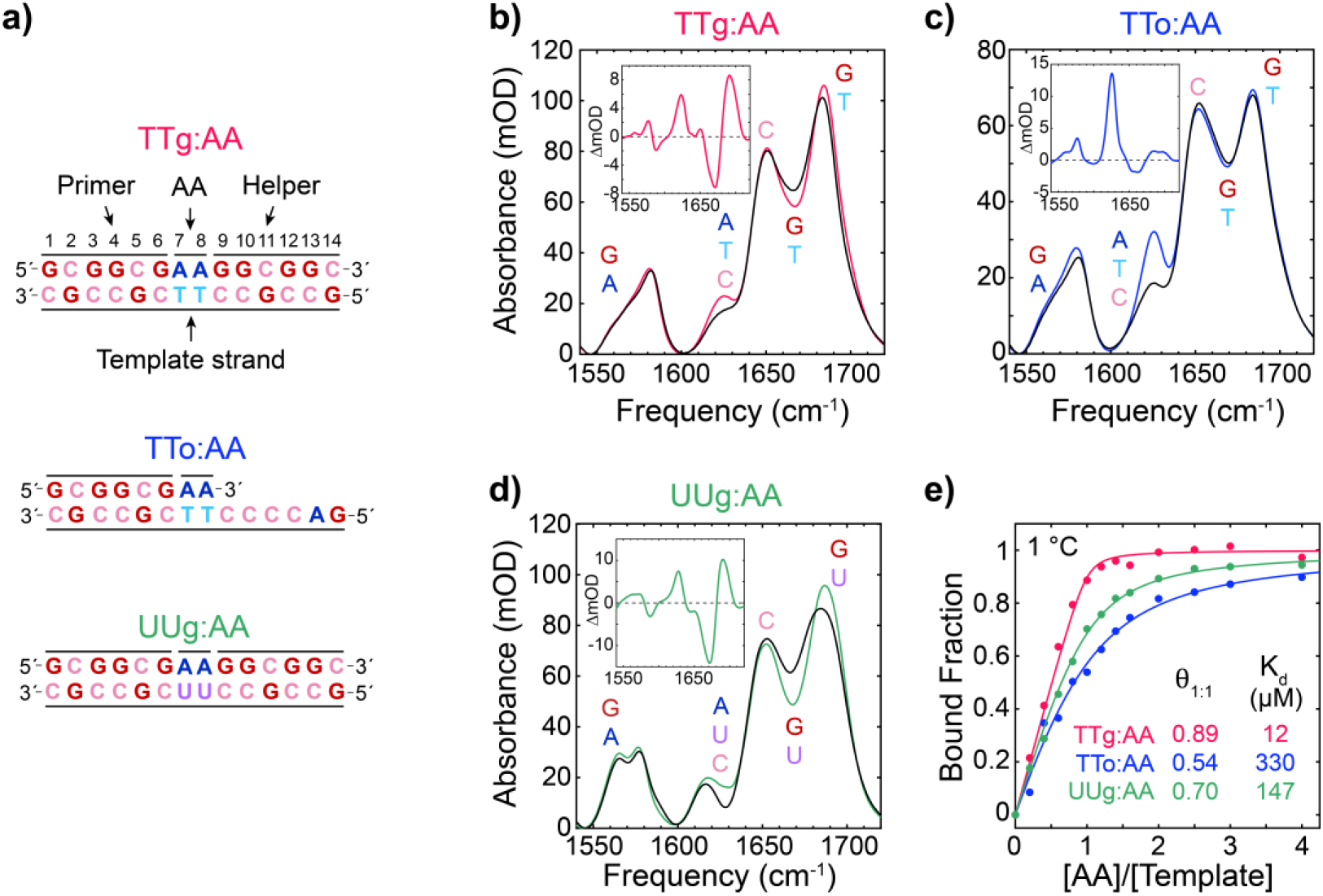
Low temperature hybridization of AA dinucleotide to gap and overhang. **(a)** Model sequences studied in this work. Gap templates are comprised of a 14-mer template strand bound to 6-mer primer and 6-mer helper strands whereas the overhang templates are only bound to the primer strand. **(b, c, d)** FTIR spectra at 1 °C of 1 mM gap and overhang templates in pH* 6.8 400 mM sodium phosphate buffer (SPB) without AA (black lines) and after adding 1 mM AA (color lines). Difference spectra upon addition of AA are shown as insets. Labels indicate the nucleobases contributing to each frequency region of the FTIR spectra. **(e)** Titrations of AA against 1 mM TTg, TTo, and UUg templates at 1 °C monitored with FTIR. The fraction of template bound to AA is determined from fitting the second singular value decomposition (SVD) component of FTIR data between 1650 – 1720 cm^-1^ to a two-state binding model (solid lines, eq. S2). The fraction of AA bound to template at a 1:1 molar AA:template ratio (*θ*_1:1_) and the dissociation constant (*K*_d_) are listed for each complex. Additional spectra and fitting details are provided in Section S1.1.

We first test for evidence that AA binds to gaps and overhangs at low temperature (1 °C) using FTIR spectroscopy. Figure 1 compares the FTIR spectra in the 1540 – 1720 cm^-1^ region of isolated TTg, TTo, and UUg templates with samples containing an equimolar mixture of template and AA. This spectral region contains in-plane carbonyl and ring stretching vibrations of the nucleobases that are sensitive to base pairing and stacking interactions and have previously been studied in detail.^41–42, 67–68^ The templates are fully intact at 1 °C such that spectral changes from adding AA report on binding of AA to the template. Each mixed sample exhibits an increase in absorbance near 1575 and 1625 cm^-1^ resulting from the A ring vibrations of AA. The 1650 – 1700 cm^-1^ region primarily contains carbonyl vibrations. The shifting of intensity from 1665 to 1685 cm^-1^ indicates increased stacking and base pairing of guanines adjacent to the gap, and the overlapping gain in intensity of the thymine carbonyl band near 1695 cm^-1^ indicates the formation of A:T base pairing.^41^ The intensity of the 1625 cm^-1^ band directly reports on the stacking and base pairing interactions experienced by AA, and its suppression in TTg:AA and UUg:AA relative to TTo:AA indicates greater binding of AA in the gap at 1°C. Further, FTIR-monitored titrations of AA with each template indicate an increase in fraction of TTg bound to AA at equimolar conditions (*θ*_1:1_) progressing from TTo:AA to UUg:AA to TTg:AA (Figs. 1e & S1). *K*_d_ decreases by over an order of magnitude in switching the template from an overhang to a gap as previously observed for binding of guanosine monophosphate (GMP) to RNA.^35^

### AA binding equilibrium monitored with temperature-dependent IR and NMR spectroscopy

To accurately measure temperature-dependent AA binding stability, IR and NMR spectroscopic probes are employed to monitor changes in base pairing as a function of temperature that are then modelled to extract melting curves for AA dissociation. FTIR temperature series were performed to track AA binding stability from 1 to 96 °C (Fig. 2a,b). Both AA-gap complexes show two melting transitions that correspond to dissociation of AA from the template followed by dissociation of the primer and helper from the template strand. The AA dissociation transition in TTg:AA and UUg:AA is observed between 1 and 60 °C as an increase in intensity of the adenine ring mode at 1625 cm^-1^ and the change of carbonyl bands at 1685-1695 and 1665 cm^-1^. The sharper dissociation transition of the primer and helper – characterized exclusively by changes in guanine ring and carbonyl bands – occurs at higher temperature as a single step melting transition since the binding stability of the primer and helper to the template strand are nearly identical. The midpoint for primer and helper dissociation is shifted 13 °C higher in UUg:AA due to the more stabilizing nature of GC base pair steps in A-RNA relative to B-DNA.^4-5^ AA unbinding in TTo:AA is already halfway complete at 1 °C as indicated in Fig. 1e and is completed by ∼40 °C (Fig. S2).

**Figure 2.**
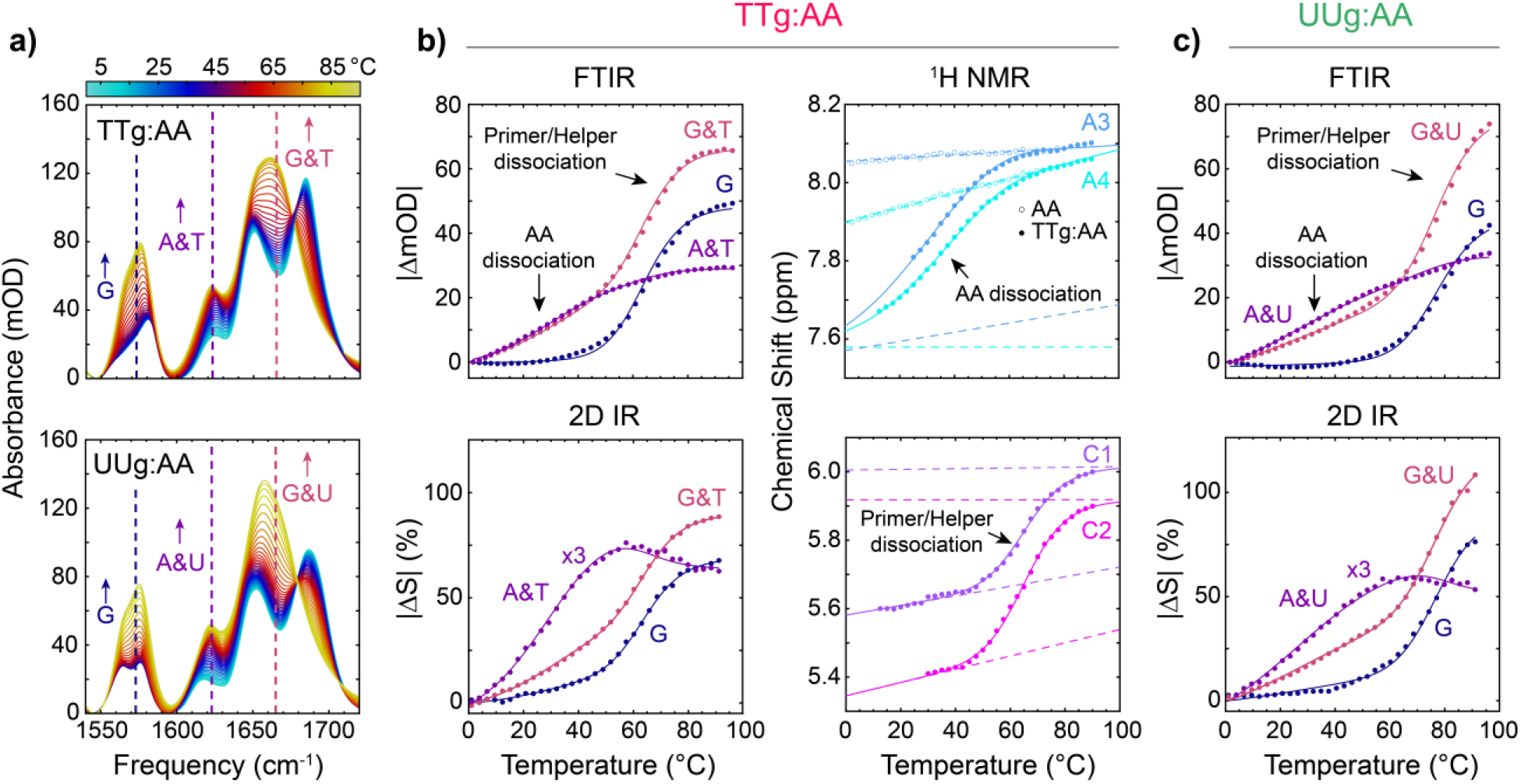
Temperature-induced dissociation of AA. **(a)** FTIR spectra of TTg:AA and UUg:AA from 1 to 96 °C in ∼2.6 °C steps. Solutions contain 1 mM each of AA, primer, helper, and template strands. **(b)** Thermal AA and template dissociation of the TTg:AA complex monitored with FTIR, 2D IR, and ^1^H NMR. FTIR traces are plotted as the absolute value change in absorption relative to 1 °C for select frequencies marked in **(a)** that primarily report on intensity changes in G (1573 cm^-1^, blue), A & T (1623 cm^-1^, purple), and G & T (1665 cm^-1^, pink). 2D IR traces at select frequencies similar to the FTIR data are plotted as the absolute value percent change in signal relative to 1 °C (|ΔS|). From ^1^H NMR, the AA dissociation transition is monitored using the chemical shift of two aromatic protons from adenine (A3 and A4), and H5 aromatic protons of cytosine (C1 and C2). The temperature-dependent chemical shifts of A3 and A4 in free AA are shown as open circles. The corresponding ^1^H NMR spectra are shown in Fig. S9. **(c)** Thermal AA and template dissociation transitions of the UUg:AA complex monitored with (top) FTIR and (bottom) 2D IR. Solid lines for each dataset correspond to fits to a three-state sequential model. FTIR and 2D IR data are globally fit together. The four ^1^H NMR peaks are globally fit to the same three-state model and include upper and lower baselines (dashed lines). Temperature-dependent FTIR and 2D IR data for TTo:AA and TTg, TTo, and UUg are shown in Sections S1.2 and S1.3.

The sigmoidal character of the AA thermal dissociation transition is superimposed on other slowly varying temperature-dependent changes, primarily a result of solvation. We account for additional independent spectral changes by measuring FTIR temperature series of TTg, TTo, and UUg and fully complementary duplexes (TTgd, UUgd) for reference, as described in Section S1.2. In contrast, we find that these linear background changes are only minor contributions in temperature series monitored by two-dimensional IR (2D IR) spectroscopy (Section S1.3). The 2D IR temperature series show profiles for AA dissociation and primer and helper dissociation transitions that correlate closely the transitions observed in FTIR data but with reduced linear baseline contribution to the sigmoidal transitions. We also obtain similar AA dissociation and primer and helper dissociation transitions from ^1^H NMR of TTg:AA (Figs. 2b and Section S1.4), lending confidence to the profiles of the IR-monitored dissociation transitions.

### Temperature-jump IR spectroscopy of AA dissociation kinetics

To directly monitor the time-dependence of AA dehybridization from gaps and overhangs, we employed T-jump IR spectroscopy^45–46^ using T-jumps of ΔT ≈13 °C across the center of the AA melting transition (Figs. 3 and S11). The absorption change at 1605 cm^-1^ by the adenine ring mode and at 1545 cm^-1^ by the guanine ring mode provide selective reporters for changes in A:T and G:C base pairing, whereas the 1672 cm^-1^ band tracks both A:T and G:C base pair disruption through the thymine and guanine carbonyl vibrations.^49, 69^ For all complexes, time traces at these three frequencies exhibit what appears to be a single kinetic component from 500 ns to 10 µs prior to decay of the signal due to thermal re-equilibration of the sample from 1 to 10 ms (Fig. 3b). The large amplitude of the A:T response at 1605 cm^-1^ and its microsecond timescale support assignment of the process to AA unbinding.

**Figure 3.**
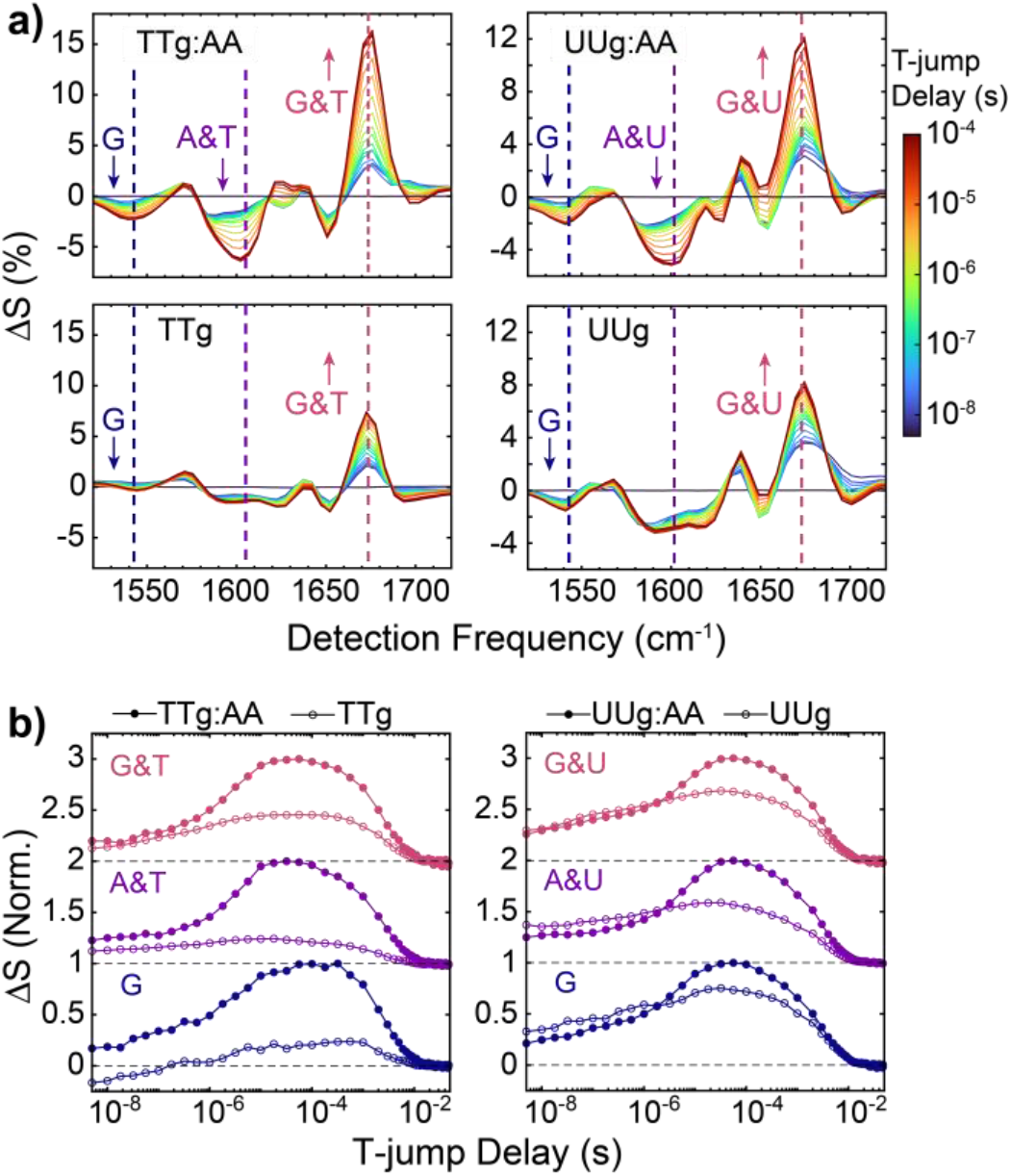
Time-dependence of AA dissociation from DNA and RNA gaps directly observed with T-jump IR spectroscopy. **(a)** Heterodyned dispersed vibrational echo difference spectra (t-HDVE) at T-jump delays ranging from 5 ns to 180 µs for TTg:AA, TTg, UUg:AA, and UUg following a T-jump from 19 to 33 °C. Spectra are plotted as the change relative to the maximum signal value of the initial temperature spectrum (Δ*S*(*t*) = *S*(*t*)/max(*S*(*T*_*i*_))). **(b)** Normalized Δ*S*(*t*) time traces at 1545 cm^-1^ (blue, guanine ring mode), 1605 cm^-1^ (purple, adenine ring mode), and 1672 cm^-1^ (red, guanine and thymine carbonyl modes) for TTg:AA and UUg:AA (filled circles) as well as TTg and UUg (open circles). The probe frequencies are marked as vertical dashed lines in the t-HDVE spectra. Traces at different frequencies are shifted vertically with respect to one another and dashed lines indicate respective baselines. AA dissociation is observed as a single kinetic component from 1 to 10 µs, and additional base pairing dynamics from the template itself occur on an overlapping timescale.

Without AA, the gap and overhang templates also exhibit a response over a similar time window but with lower amplitude, particularly at the adenine ring mode, relative to the AA-template complexes. The dominant G:C character of the response suggests that it may arise from base pair disruption adjacent to the gap or at the primer and helper termini; however, the same response is also observed in the fully formed duplexes (Figs. S12-S13), which indicates it arises from fraying of the primer and helper G:C termini. This observation is consistent with previous NMR studies that found G:C base pairs adjacent to a gap to be more stable than those at the duplex termini,^28^ and T-jump experiments indicating G:C fraying in DNA oligonucleotides on microsecond timescales.^70^ G:C fraying occurs faster than AA dissociation yet with enough temporal overlap that only a single kinetic component is resolved at each frequency. AA dissociation dominates the observed kinetic component at most frequencies, but the signal change at 1672 cm^-1^ contains enough amplitude from G:C fraying that the extracted observed rate is faster than at 1605 cm^-1^ for TTg:AA at low temperature (Figs. S14-S15).

### Thermodynamic analysis of AA dissociation

The results demonstrate steady-state and time-resolved thermal dissociation of AA, which we self-consistently model to extract quantitative thermodynamic and kinetic information. In general, complete dissociation of the AA-template complex may proceed through multiple pathways differentiated by the order in which AA, primer, or helper segments dissociate. Exact treatment of these coupled equilibria leads to a fourth or higher-order dependence of each binding fraction on the concentration of a given species, which we find does not have a stable solution over the full temperature range of each transition. However, the much greater binding stability of the primer and helper allows us to neglect species with AA bound to the template strand without the primer and helper. We also model primer and helper dissociation from the template strand as a unimolecular equilibrium between the template and “dissociated” template that is equivalent to treating binding of the primer and helper as folding of hairpins.^16^ This results in a three-state sequential model described by equilibrium constants for AA dissociation (*K*_d_) and dissociation of the primer and helper (*K*_d,Temp_). Using TTg:AA as an example:

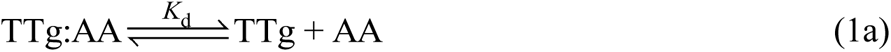

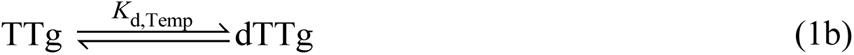

where dTTg is the “dissociated” template. The assumption in eq. 1b sharpens the primer and helper melting profile relative to its true bimolecular form as illustrated in Fig. S23, but this is not a serious concern because this only affects our analysis in the temperature range where the transitions overlap.

To extract melting curves for AA dissociation, FTIR and 2D IR temperature series for a given sequence are globally fit to eqs. 1a-b using the spectral decomposition described in Section S3 and constraints from the titration at 1°C. The resulting melting curves are shown in Fig. 4a and the corresponding spectral components are shown in Figs. S20-S21. Overall, the global fit describes both data sets well (Fig. 2). The first spectral component corresponds to AA dissociation and contains signatures of A:T base pair breaking whereas the second component contains only G:C base pair breaking features. The AA dissociation curves resemble the raw temperature-dependent change in absorbance and start from a bound fraction less than 1 at 1 °C. We also find quantitative agreement between the TTg:AA dissociation curves determined from FTIR and 2D IR and from global fitting of temperature-dependent ^1^H NMR chemical shifts (Fig. S9), lending support to the accuracy of the IR dissociation curves.

**Figure 4.**
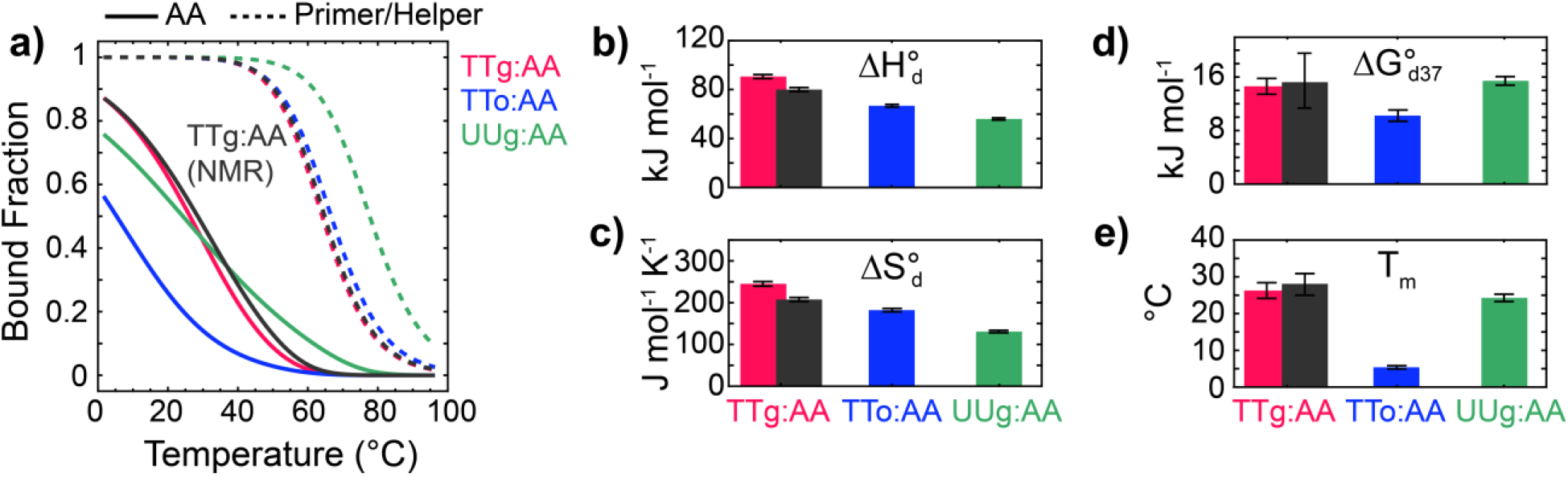
Thermodynamics of AA dissociation from gap and overhang templates extracted with three-state global fitting of FTIR and 2D IR temperature series. **(a)** AA (solid) and primer and helper (dashed) dissociation curves for TTg:AA, TTo:AA, and UUg:AA extracted from global fitting of FTIR and 2D IR temperature series to a three-state sequential model (eqs. 1a-b, Section S3). Dark gray lines correspond to dissociation components of TTg:AA obtained from ^1^H NMR temperature series (Fig. S9). AA dissociation **(b)** enthalpy 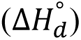), **(c)** entropy 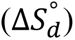, **(d)** free energy at 37 °C 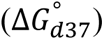, and **(e)** melting temperature (*T*_*m*_) determined for each sequence. Dark gray bars indicate values from ^1^H NMR. Error bars are derived from 95% confidence intervals in fit parameters 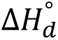 and 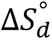.

**Table 1.** Thermodynamics for dissociation of AA from templates determined with global fitting to three-state sequential model (eqs. 1a-b). Error bars are derived from 95% confidence intervals in fit parameters 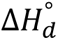 and 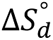.

| Sequence | Method | $T_m^c$<br>(°C) | $\Delta H_d^o$<br>(kJ mol<sup>-1</sup>) | $\Delta S_d^o$<br>(J mol<sup>-1</sup> K<sup>-1</sup>) | $\Delta G_{d8}^o$<br>(kJ mol<sup>-1</sup>) |
| --- | --- | --- | --- | --- | --- |
| TTg:AA | IR<sup>a</sup> | 26 ± 2 | 91 ± 2 | 245 ± 6 | 21.7 ± 1.1 |
|  | NMR | 28 ± 3 | 79 ± 5 | 203 ± 20 | 21.5 ± 4.5 |
|  | ITC<sup>b</sup> | - | 83 ± 5 | - | 23.1 ± 1.3 |
| UUg:AA | IR<sup>a</sup> | 24 ± 1 | 56 ± 1 | 131 ± 3 | 19.2 ± 0.6 |
|  | ITC<sup>b</sup> | - | 51 ± 1 | - | 23.5 ± 0.4 |
| TTo:AA | IR<sup>a</sup> | 5 ± 1 | 67 ± 1 | 182 ± 4 | 16.1 ± 0.8 |

<sup>a</sup>IR refers to global fits of FTIR and 2D IR temperature series. <sup>b</sup>Isothermal titration calorimetry (ITC) was performed at 8 °C. <sup>c</sup> $T_m$ is defined as the temperature where half of the AA is bound to the template. <sup>d</sup>Free energy change for AA dissociation at 8 °C.

The shape of AA dissociation curves varies between DNA and RNA as well as the type of template. TTg:AA and UUg:AA exhibit similar transition midpoints (*T*_*m*_), defined as the temperature where half of the total AA concentration is bound to the template, but UUg:AA shows a broader transition with a dissociation curve slope at *T*_*m*_of 0.013 K^-1^ relative to 0.021 K^-1^ in TTg:AA. The broader transition width in UUg:AA stems from a 35 kJ mol^-1^ lower dissociation enthalpy 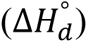 and 115 J mol^-1^ K^-1^ lower dissociation entropy 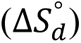 relative to TTg:AA. This difference in 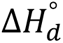 is also directly observed through ITC measurements (Section S4.2). It stems from the 5-methyl group of thymine that is known to enhance π-π stacking. Therefore substitution to uracil in RNA is expected to reduce base pair stacking.^71–72^ The AA dinucleotide is rather unique in this regard as most RNA dinucleotide steps exhibit larger 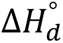 and 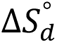 than DNA.^4-5^ The free energy of AA dissociation at 37 °C is roughly 4 kJ mol^-1^ higher in TTg:AA than TTo:AA due to the added stacking stabilization from the 3′-G, and this value is similar to previous measurements of coaxial stacking in 5′-AG-3′ steps of DNA oligonucleotides.^15, 17–18^

### Kinetic analysis of AA dissociation

The observed relaxation rates for AA dissociation kinetics, *k*_*obs*_, are extracted from T-jump measurements with varying initial temperature across the AA melting transition of each sequence (Figs. 5 & S11-S13). Only AA dissociation and G:C fraying are observed below final temperatures *T*_*f*_ = 42 °C for DNA and 52 °C for RNA. At higher temperatures, primer and helper dissociation is observed on 100 µs timescales and is more than 100-fold slower than AA dissociation (Fig. S13). The timescale separation between AA dissociation and primer and helper dissociation is sufficient to treat *k*_*obs*_ with two-state kinetics.^73^ Using TTg:AA as an example,

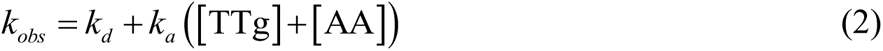

**Figure 5.**
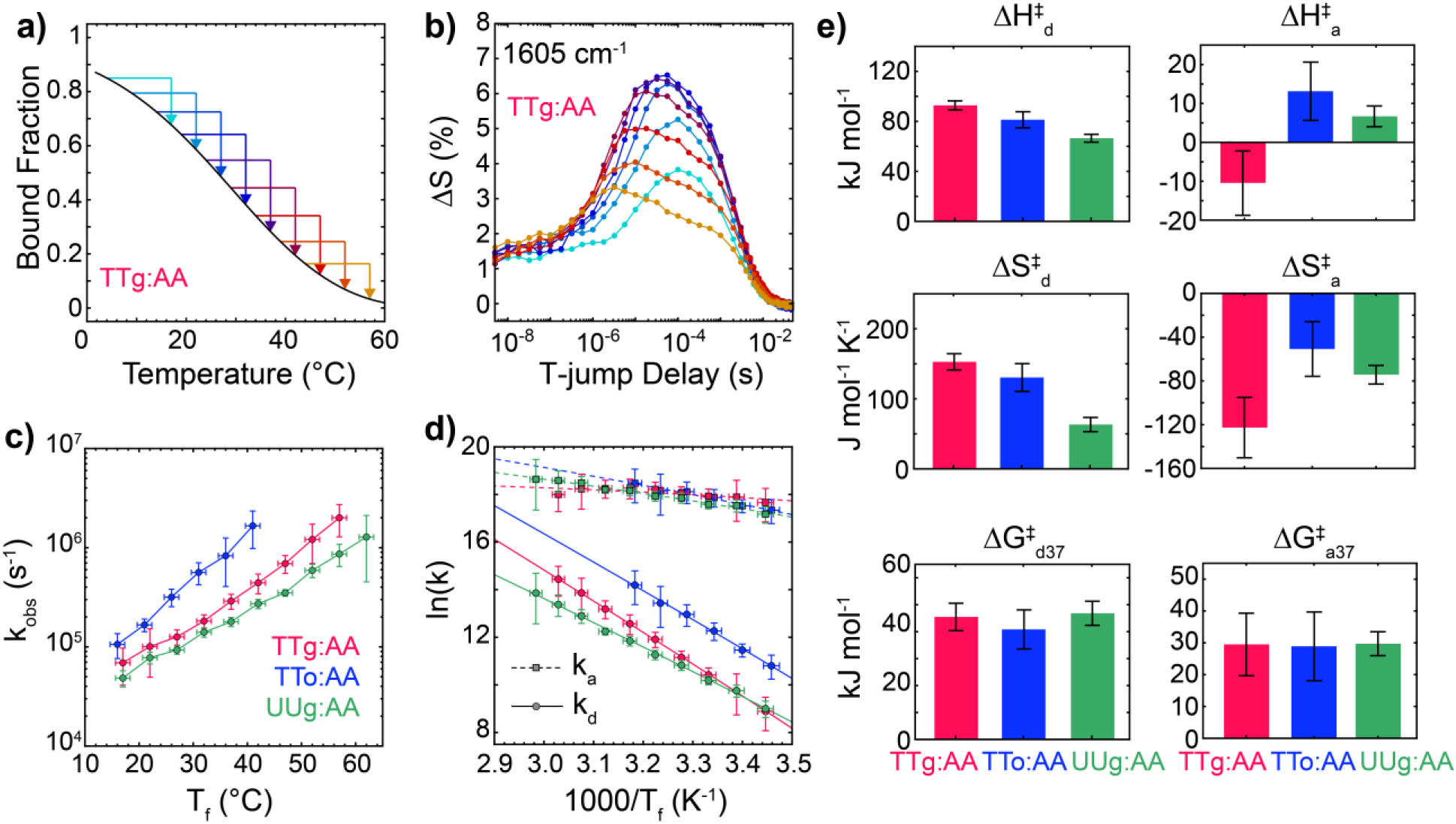
Kinetics of AA association and dissociation from gaps and overhangs. **(a)** Illustration of a series of T-jumps performed from various initial to final temperatures (*T_i_* →*T_f_*) with ΔT∼13 °C across the AA dissociation transition of TTg:AA. **(b)** t-HDVE time traces of TTg:AA probed at the adenine ring mode (1605 cm^-1^) for each temperature condition shown in **(a)**. **(c)** Observed relaxation rate (*k*_*obs*_) for AA unbinding as a function of *T_f_* for all sequences. Rates correspond to the amplitude-weighted mean across the adenine ring mode response (1585 to 1610 cm^-1^) in the rate-domain spectra determined from MEM-iLT.^48–49^ Vertical error bars indicate standard deviation over the rate spectra and horizontal error bars correspond to the measured standard deviation in T-jump magnitude. **(d)** Rate constants for AA association (*k*_*a*_) in M^-1^ s^-1^ and dissociation (*k*_*d*_) in s^-1^ as a function of *T_f_* determined from a two-state analysis of *k*_*obs*_ (eq. 2). Data for all sequences is shown in Fig. S27. **(e)** Enthalpic 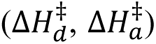, entropic 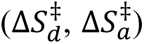, and free energy (37°C, 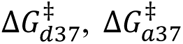) barriers determined from fitting *k*_*d*_ and *k*_*a*_ temperature-trends to a Kramers-like model (eq. 3). Error bars indicate 95% confidence intervals propagated from the fits.

where *k*_*a*_and *k*_*d*_are the rate constants for AA association and dissociation. Since *K*_*d*_ = *k*_*d*_/*k*_*a*_, these rate constants are extracted using the concentrations of template, [TTg], and AA, [AA], determined from the AA dissociation curves in Fig. 4. Due to the broad observable temperature range of AA-gap dissociation transitions, T-jump measurements can be performed at multiple *T_f_* values below *T*_*m*_ where *k*_*obs*_ is dominated by *k*_*a*_. Over this temperature range, the dissociation rate follows Arrhenius behavior and increases from 3 × 10^3^ to 2 × 10^6^ s^-1^ while *k*_*a*_ shows just a minor 2-to-5-fold increase. Such contrasting temperature-dependence of *k*_*a*_ and *k*_*d*_ has consistently been observed in studies of short DNA and RNA oligonucleotides (5 – 20 bp),^6-7, 49, 74–75^ and has been used to characterize the transition-state between hybridized and dissociated states. We fit the temperature trends of *k*_*a*_ and *k*_*d*_ to a Kramers-like model for diffusive barrier crossing where the rate is inversely proportional to the solvent viscosity, *η*. Written for the dissociation rate:

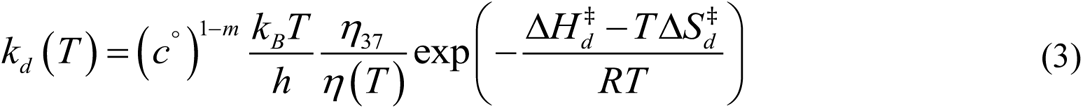

and the same expression applies for association. Here the free energy barrier 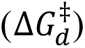 is written in terms of the enthalpic 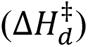 and entropic 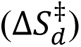 barriers. The temperature-dependent viscosity of D_2_O is taken relative to its value at 37 °C (*η*_37_/*η*).^76^ Also, c° is the standard state concentration of 1 M, and *m* is the reaction molecularity (*m* = 1 for *k*_*d*_ and *m* = 2 for *k*_*a*_). We use a simple form of the Eyring pre-factor (*k*_*B*_*T*/ℎ). This approximation is known to overestimate the frequency of diffusive motion at the barrier for biomolecular folding and binding.^77–78^ While this impacts the absolute magnitudes of 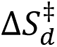 and 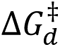, we assume that the true pre-factor is the same across our sequences such that relative values of 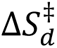 and 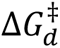 are meaningful. Fitting of our data indicates that the free energy barrier for AA dissociation arises from an enthalpic penalty (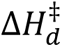 > 50 kJ mol^-1^) that is partially compensated by a positive 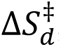, whereas the free energy barrier for AA association 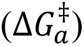 is dominated by an entropic barrier (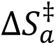 < 0).

The kinetics of AA association and dissociation vary substantially depending on the template. 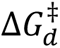 is ∼4.5 kJ mol^-1^ smaller for TTo:AA than TTg:AA at 37 °C, which matches the difference in 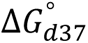, suggesting that the 5-fold faster unbinding of AA from an overhang compared to a gap primarily comes from reduced binding stability. The rate of association or 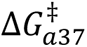 are essentially the same for TTo:AA and TTg:AA but with small differences in its temperature-dependence as quantified by 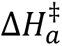. There are also significant differences between the kinetics of AA dissociation in UUg:AA and TTg:AA. *k*_*d*_ increases more sharply with temperature in TTg:AA, as quantified by a ∼25 kJ mol^-1^ larger 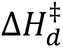 relative to UUg:AA, which likely comes from enhanced stacking interactions in the hybridized state of TTg:AA and is consistent with the thermodynamic results presented above. On the other hand, *k*_*a*_ is similar for each sequence across the measured temperature range.

### Structural properties of AA-gap complexes from molecular dynamics simulations

The energetics of AA hybridization onto overhangs and gaps differs from free strand hybridization due to a combination of differences in the template and AA-gap complex structural configurations relative to canonical duplexes and single-strands. We characterized the differences in conformation between these species using all-atom MD simulations, finding that most configurational differences are localized near the central AA binding site. As observed previously, gap templates adopt bent configurations in high population which reduces hydration of the gap relative to single strands (Figs. S29-S30).^27–28, 79^ AA-gap complexes exhibit greater configurational freedom than canonical duplexes (Figs. 6 & S31-S33), with the largest differences found in backbone and base-stacking structure rather than base pair geometry. TTg:AA exhibits broadened distributions in shift, slide, and twist parameters at the 5′-GA-3′ and 5′-AG-3′ nicks, whereas the stacking geometry of the between adenine bases is nearly identical to B-DNA (ΔDx, ΔDy < 0.4 Å and ΔΩ < 4°). A broader spread of stacking orientations at the nick sites is unsurprising given the greater flexibility in the backbone, particularly for glycosidic bond angles (χ) of the A and G bases on the 3′-side of each nick.^22^ The nick sites enable greater local flexibility in the backbone, and in particular for the glycosidic bond angles (χ) of the A and G bases on the 3′-side of each nick. The greater freedom of the backbone near the nick sites also reduces the effective population of BII backbone configurations across the 5′-CGAAGG-3′ motif (Fig. S34). The fluctuations along slide, twist, shift, and χ coordinates are partially correlated in a given nick site (Fig. 6b). In contrast, the structural fluctuations in the 5′-GA-3′ and 5′-AG-3′ nicks appear largely independent of one another (Fig. 6c).

**Figure 6.**
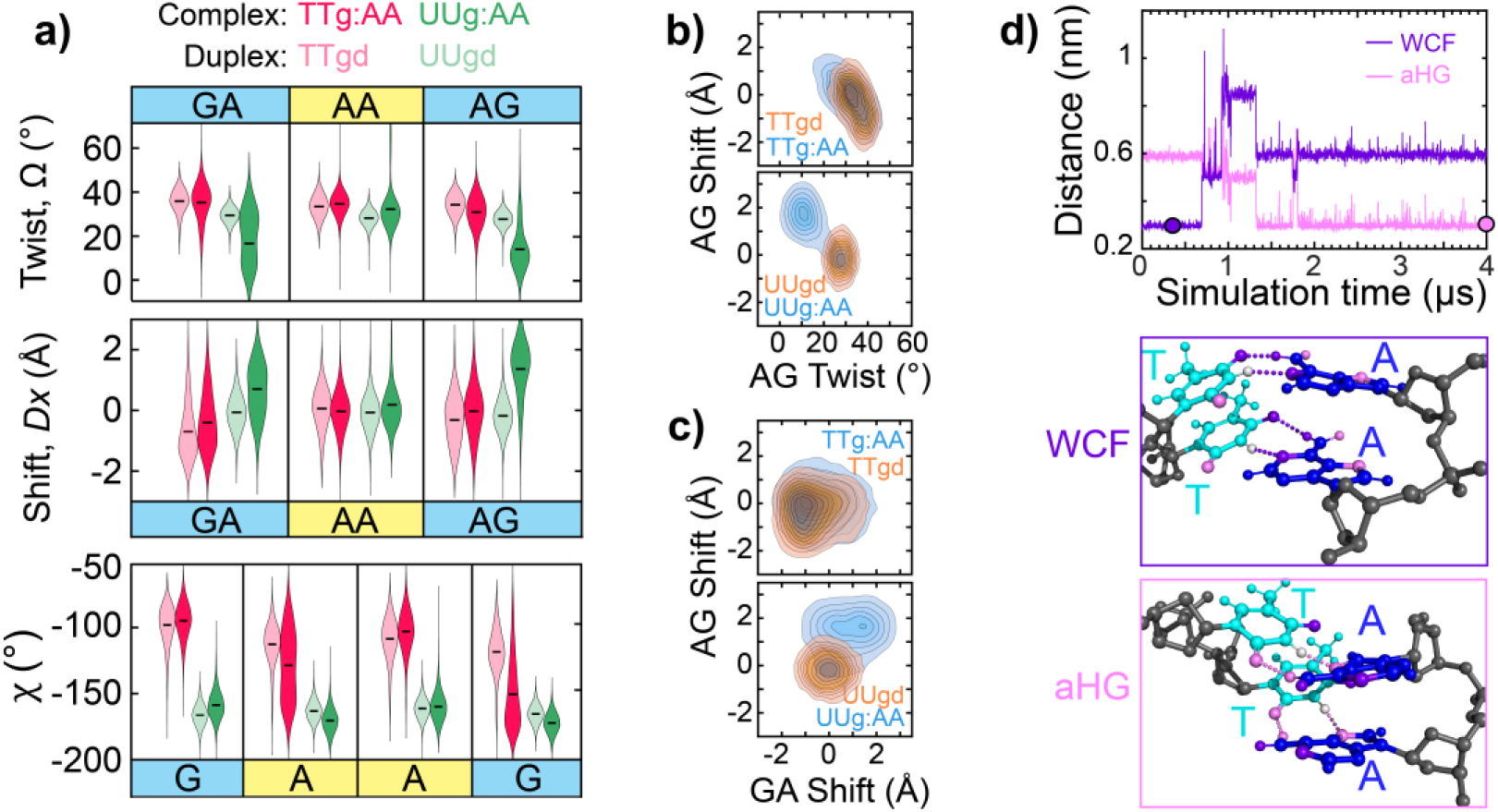
Structural properties of AA-gap complexes from all-atom MD simulations. **(a)** Comparison of structural variation between AA-gap complex and canonical duplex. Violin plots show structural distributions for base pair twist (Ω) and shift (Dx), and glycosidic bond dihedral angle (χ) for the three central base pair steps of TTgd (red, light), TTg:AA (red, dark), UUgd (green, light), and UUg:AA (green, dark). Black horizontal bars indicate the distribution’s median value obtained from a Gaussian kernel estimation using Scott’s rule.^80^ Distributions were generated from 1.5 µs of simulation for each sequence using the bsc1-AMBER force field for DNA and DES-AMBER force field for RNA. Only Watson-Crick-Franklin (WCF) base pair configurations are included in the distributions. Other structural parameters using both force fields are shown in Figs. S31-S33. **(b)** Contour plots showing the correlation of 5′-AG-3′ twist vs. shift. Nine contours with uniform 10% spacing are plotted for each sequence. Diagonally elongated distributions indicate that fluctuations along different base pair coordinates within the same nick site are partially correlated with Pearson correlation coefficients of −0.61 for DNA and −0.56 for RNA. **(c)** Contour plots showing the correlation of 5′-GA-3′ shift vs. 5′-AG-3′ shift. Symmetrical distributions indicate that fluctuations at each nick site are uncorrelated with Pearson correlation coefficients of −0.04 for DNA and 0.05 for RNA. **(d)** 4 µs trajectory of TTg:AA showing the A:T base pairs switching from WCF to an anti-Hoogsteen (aHG) geometry. Trajectories are plotted in terms of the mean WCF (purple) and aHG (pink) distances across each A:T base pair. The WCF distance is defined as the mean of adenine-N6 to thymine-O4 and adenine-N1 to thymine-N3 distances and the aHG distance is the average of adenine-N7 to thymine-N3 and adenine-N6 to thymine-O2 distances. Visualizations of each base pair geometry at time points marked with filled circles indicate WCF distances as purple dashed lines and aHG distances as pink dashed lines.

The structural differences between UUg:AA and an A-RNA duplex are also minor yet distinct from DNA in a few regards. Overall, UUg:AA exhibits larger changes in stacking orientation (twist, shift) and less change in the backbone torsional angles (χ, δ, γ) relative to DNA. Further, the base pairing and stacking geometry of the AA differs with broader distributions of shear, stretch, and buckle values, an increase in mean rise from 3.0 to 3.3 Å, and sub-populations with larger shift and twist values. In contrast to DNA, distinct states are observed along twist and shift coordinates for the 5′-GA-3′ and 5′-AG-3′ and the mean values differ from A-RNA by as much as 20° and 1.5 Å, respectively.

During one of two 4 μs trajectories for TTg:AA, we observed AA switch from canonical Watson-Crick-Franklin (WCF) to anti-Hoogsteen (aHG) base pairing geometry. As shown in Fig. 6d, the 5′-A flips at 700 ns followed by the 3′-A 200 ns afterward, however, both adenines remain poorly aligned with the gap until reaching the final aHG geometry at 1.3 μs. Our 8 µs of total simulation time for this sequence is insufficient to determine the relative stability of WCF and HG configurations, therefore we applied the On-the-fly Probability Enhanced Sampling (OPES) method to calculate the free energy differences for AA dissociation from a WCF or aHG geometry (ΔΔ*F*_*WCF*−*aHG*_ = Δ*F*_*d*37,*aHG*_ − Δ*F*_*d*37,*WCF*_; See Section S5.3). WCF and aHG geometries have nearly the same population and stability at 37 °C with ΔΔ*F*_*WCF*−*aHG*_ ≈ 0 kJ mol^-1^. Although not previously tested for dinucleotide-gap complexes or aHG base pairing, simulations with the bsc1-AMBER force field typically capture accurate relative stabilities between WCF and syn-Hoogsteen base pairing geometries.^81–82^ Simulations with the DES-AMBER force field instead suggest that the WCF geometry is more stable (ΔΔ*F*_*WCF*−*aHG*_ ∼ −8 kJ mol^-1^). For UUg:AA, WCF and aHG geometries have similar stabilities regardless of the employed force field (Fig. S35). The high stability of an aHG geometry may appear unusual, but this has been reported as the dominant base pairing geometry between AA and TT DNA dinucleotides mediated through self-assembling tripyridyl-triazinine capped hydrophobic cages in aqueous solution, so far the only structural characterization of dinucleotide duplexes in aqueous solution.^83^

### Length scaling of A_n_ dissociation from templates

To assess the scaling of dissociation thermodynamics and kinetics for 2-to-4 nucleotides sequences, we experimentally examined the dehybridization of AAA (A3) and AAAA (A4) from DNA gaps and overhangs using the extended templates shown in Fig. 7a. As expected, longer A_n_ oligonucleotides bind with greater stability and show a decrease in both *k*_*d*_and *k*_*a*_(Figs. 7b-d & S27). Both gaps and overhangs exhibit an average increase in 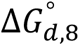 of 2.7-3.0 kJ mol^-1^ per base pair (bp), yet 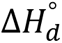 and 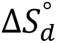 increase more sharply with length for overhangs and become equivalent to the gap for A4 dissociation. The overhang values for 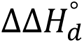, 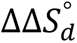, and 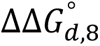 are still well below the 33 kJ mol^-1^ bp^-1^, 94 J mol^-1^ K^-1^ bp^-1^, and 6.7 kJ mol^-1^ bp^-1^ values predicted for appending AA dinucleotide steps by Santa Lucia’s DNA nearest-neighbor (NN) model (Fig. 7e).^4^

**Figure 7.**
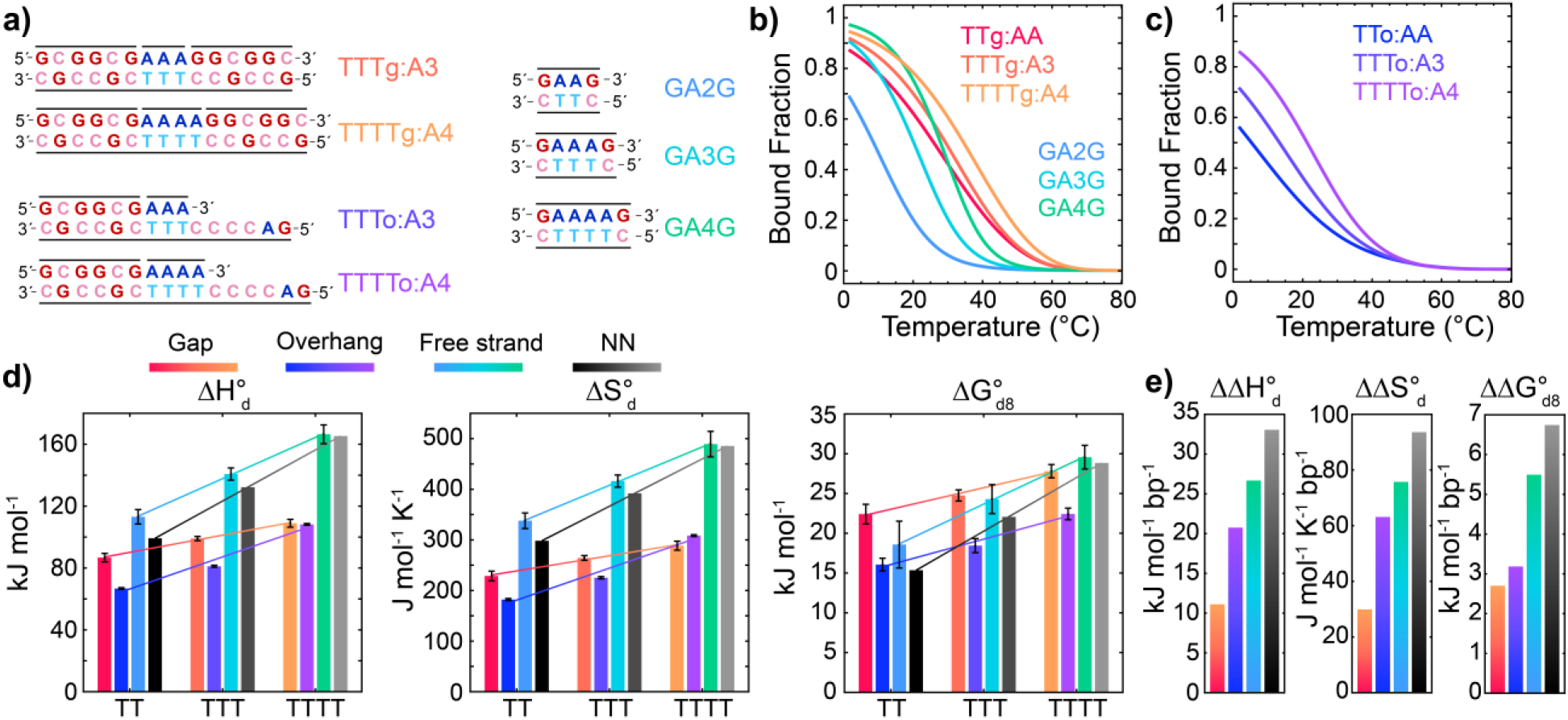
Comparing length-dependent dissociation thermodynamics in gaps, overhangs, and free strands. **(a)** Extended templates for 3 and 4 nucleotide gap (TTTg:A3, TTTTg:A4) and overhang (TTTo:A3, TTTTo:A4) sequences bound to A3 and A4 oligonucleotides. Dissociation of A_n_ (n = 2, 3, 4) segments from gaps and overhangs are compared with dissociation of GA_n_G single-strands with their complement strands. **(b, c)** A_n_ melting curves for gaps and overhangs extracted from global fitting of FTIR and 2D IR temperature series to three-state sequential model (eq. 1). GA_n_G melting curves are shown in **(b)** and were extracted from a two-state model fit to FTIR temperature series (Section S4.1). GA_n_G melting curves were measured at 10 mM oligonucleotide concentration and corrected to 2 mM. **(d)** 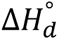, 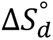, and dissociation free energy at 8 °C 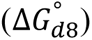 for all sequences. Red-orange bars correspond to DNA A_n_-gap complexes, blue-purple bars correspond to DNA A_n_-overhang complexes, light blue-green bars are for GA_n_G strands, and black-gray indicate GA_n_G dissociation parameters calculated from Santa Lucia’s nearest-neighbor (NN) model and corrected to [Na^+^] = 600 mM.^4, 84^ Values for gap and GA_n_G systems are an average over those determined from IR spectroscopy and ITC (Section S4.2) while overhang values are only from FTIR and 2D IR global fits. Error bars indicate 95% confidence intervals from fits to thermodynamic models. **(e)** Slopes from linear fits of each thermodynamic parameter across A_n_ length (solid lines in **(d)**). Data are presented as the change per appended A:T base pair (bp) in the dissociation enthalpy 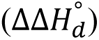,entropy 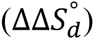, and free energy at 8 °C 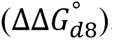. Although AA-gap complex binding is more stable than GA2G, 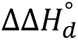, 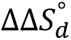, and 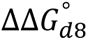 are each more than 2-fold greater in GA_n_G strands than for A_n_ dissociation from gaps.

To test for the origin of different length-scaling dissociation thermodynamics between overhangs, gaps, and the NN model, we compare to dissociation of oligonucleotides 5′-GAAG-3′ (GA2G), 5′-GAAAG-3′ (GA3G), and 5′-GAAAAG-3′ (GA4G) with their respective complement strands (Figs. 7 & S24-S26). 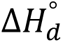 and 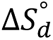 are both greater for GA2G than AA-gap dissociation while 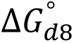 is ∼4 kJ mol^-1^ lower for GA2G. As a result, the GA2G dissociation transition is much sharper than for AA-gap dissociation yet with *T_m_* shifted ∼15 °C lower in temperature. Values of 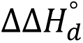, 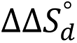, and 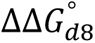 for GA_n_G are more than 2-fold greater than for gap sequences and are much closer the NN model values (Fig. 7e). This leads to nearly equivalent 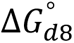 for TTTTg:A4 and GA4G, and binding between free single-strands may progressively become more stable than for binding of longer oligonucleotides onto gaps and overhangs. Additionally, we find that length-scaling of the dissociation free energy barrier 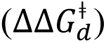 is ∼2.5-fold greater than 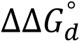 for gaps (Fig. S28), which is a much larger difference than previously reported for dehybridization of free single-strands.^74, 85^ This sharp dependence of 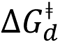 (and *k*_*d*_) on length is important for predicting dissociation kinetics in template systems and also indicates a length-dependence in the free energies of the association transition state and/or the unbound state.^86–87^

Based on previous studies of gap and overhang structure,^23–28^ it is possible that the single-stranded binding region in gaps and overhangs becomes less ordered as the length increases from 2 to 4 nucleotides. However, our all-atom MD simulations indicate essentially no difference in the structural distribution of the canonical unbent T_n_ configurations in gaps, but instead highly bent configurations and alternative stacking geometries are observed for certain gap lengths (Fig. S29). AMBER force fields, including bsc1-AMBER, tend to overestimate base-stacking stability in single-strand DNA as they are parameterized for duplex DNA,^56, 88^ and therefore these results should be interpreted with caution.

## Discussion

We demonstrate an approach for measuring temperature-dependent binding thermodynamics and kinetics of nucleic acids as short as dinucleotides onto gaps and overhangs using IR spectroscopy. By determining accurate melting curves, our method extracts enthalpic and entropic contributions to binding that are typically missing in previous titration studies. Further, to our knowledge we provide the first measurements of dinucleotide association and dissociation timescales (0.5 – 20 µs) through the application of T-jump IR spectroscopy. Although our study is limited to pure A_n_ segments, we identify striking energetic differences between cases of association onto gaps or overhangs with free-strand association that have not been observed previously.

Most notable is that association of AA onto a gap with G:C base pairs on each side is energetically more favorable than association between 5′-GAAG-3′ and 5′-CTTC-3′ (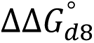 = 4 kJ mol^-1^, Fig. 7d). This distinction may arise from energetic differences in either the bound or dissociated states or a combination of both through factors such as single-strand stacking, optimized coaxial stacking, or non-canonical base pairing. Most association measurements, including the approach in this work, directly probe energetic differences between the bound and dissociated states but not energetic differences between the respective bound or dissociated states of different samples. As a result, we cannot directly dissect the enhanced stability of AA-template complexes but rather infer how the energy of the associated and dissociated states may change between gap, overhang, and free single-strand scenarios.

A clear distinction between gap and free single-strand binding scenarios is present in both the enthalpic and entropic contributions to association. 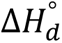 and 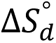 are lower for the gap, and therefore the extra stability of TTg:AA relative to GA2G arises from an entropic benefit. Enhanced binding stability of dinucleotides onto gaps and overhangs may arise from greater base pair ordering and constraints in water dynamics within the single-strand binding region and/or increased configurational flexibility in the AA-template complexes relative to canonical B-DNA, which we observe in MD simulations near the nick sites (Figs. 6a & S30-S31). The former effect decreases the entropy of the dissociated state while the latter increases that of the hybridized state, each of which will decrease 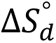 relative to free stand hybridization. The same effect that may increase rigidity and stacking interactions in the gap single-strand will also reduce the enthalpy of the dissociated state. Another reduction in enthalpy may arise from bending about the single-stranded segment in gap templates as observed in our MD simulations (Fig. S29) and reported in previous studies.^26, 28–29, 89–90^ Lastly, our simulations of TTg:AA suggest that AA can bind in WCF or aHG geometries with significant population, although the relative stability depends on the choice of force field (Fig. S35). aHG configurations are likely only relevant to binding of pure A:T or A:U dinucleotides or G:G mismatch pairing where the aHG configuration is a stable option. To date, the only crystallographic studies of dinucleotide binding to a gap or overhang template are for G:C base pairing of modified GG RNA dinucleotides,^91–93^ a scenario where aHG configurations are highly improbable. Unfortunately, we cannot indicate nor disprove the presence of A:T aHG base pairing using the IR and NMR measurements presented in this work, and further structural characterization is necessary to clarify the role of aHG base pairing.

The energetics of dinucleotide association onto gaps and overhangs differs from free-strand hybridization, and these effects stem from underlying dynamics of association and dissociation. Our T-jump IR measurements of DNA and RNA AA association and dissociation kinetics provide a first step toward understanding the structural dynamics. We determine a *k*_*a*_ value of ∼5 × 10^7^ M^-1^ s^-1^ that corresponds to a time constant of 20 µs at 1 mM oligonucleotide concentration. This *k*_*a*_ value is nearly 100-fold slower than an estimated diffusion-limited rate constant (*k*_*diff*_) of 5 × 10^9^ M^-1^ s^-1^ calculated using the translational diffusion coefficients of AA and TTg determined from DOSY NMR measurements and estimated radii of gyration from experimental length-scaling of ssDNA and worm-like chain model of duplex DNA (Section S6).^94–95^ Although faster than association between oligonucleotides of 5 – 20 nucleotide lengths, the 100-fold difference between AA association and diffusion-limited rates suggest that numerous unsuccessful collisions and/or significant structural rearrangement occur prior to successful binding. Association rate constants for longer oligonucleotides are also consistently reported to be 100-1000-fold smaller than *k*_*diff*_,^74^ suggesting that each system’s hybridization transition-state and encounter complex may share certain characteristics. Even for longer oligonucleotides these properties are still poorly understood but are thought to involve some degree of single-strand ordering, a small number of base pair contacts, and potentially rearrangement of water molecules and counterions. In the case of dinucleotide association to gaps and overhangs, water and counterion rearrangement is still necessary, and the ordering of the binding region of the gap or overhang contains a large penalty. The time ordering of these events are unclear for each system and must be investigated further to develop a molecular picture for the dynamics of short oligonucleotide hybridization.

We have primarily focused on short DNA oligonucleotide association, and it is important to assess whether similar characteristics are found for short RNA oligonucleotides, which are more relevant in the context of non-enzymatic replication.^96^ Our results show large differences in energetics and kinetics of AA unbinding from DNA and RNA gaps. 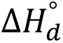 and 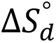 values for UUg:AA are approximately half of those predicted by NN models and obtained for TTg:AA (Fig. S26). For this particular sequence, 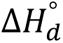 and 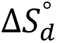 are likely higher in DNA due to enhanced stacking arising from the C5-methyl groups of thymine.^71–72, 97^ However, RNA duplexes are generally more thermodynamically stable than an equivalent sequence DNA duplex due to multiple effects from the C2′-OH groups. Differences in base stacking, hydrogen bonding between C2′-OH groups and water molecules, and greater uptake and ordering of cations and water molecules are all effects that contribute to larger 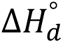, 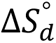, and 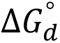 in A-RNA than B-DNA.^98–99^ Therefore, we predict that binding of other RNA dinucleotide sequences will generally be more stable than the equivalent DNA sequence.

Gaps and overhangs are expected to have a qualitatively similar thermodynamic and kinetic impact on DNA and RNA hybridization, but the relative magnitude may generally differ. For instance, single- and double-strand RNA are shown to have a longer persistence length than DNA and the change in strand rigidity upon introducing an overhang or gap may differ as well as the propensity for bending of gapped RNA.^30, 100^ MD simulations of UUg:AA and UUgd indicate that the distribution of stacking geometries at nick sites between primer and helper segments and AA differ from A-RNA while the backbone geometry is relatively unchanged, yet the opposite is found for DNA (Fig. 6a). Such differences in geometry may influence the enthalpy of the AA-gap complex while the enhanced configurational flexibility increases its entropy. Together, the combination of differences between DNA and RNA templates and dinucleotide-template complexes may promote significant contrast in their association energetics and kinetics across variations in dinucleotide sequence and template construction.

Our experimental and computational results indicate multiple structural, energetic, and kinetic factors that differentiate hybridization between short nucleic acids and templates with hybridization between free single-strands, and these differences may play an important role in processes such as non-enzymatic replication and toehold-mediated strand displacement. In addition to various structural and chemical factors, the efficiency of these processes depends upon the binding probability, dissociation rate of the bound complex, and structural dynamics of the complex.^93, 101–103^ An understanding of these factors for hybridization between short nucleic acids and templates is still developing, and this study demonstrates a method for direct measurement of energetics and kinetics for dinucleotide binding that is applicable to a wide range of oligonucleotide sequences and template designs.

## Data Availability

Molecular dynamics and enhanced sampling input files and Python scripts for plotting helical parameter distributions and reweighting OPES trajectories are available at https://github.com/mrjoness/dinucs/. All unbiased and biased MD trajectories, time- and rate-domain t-HDVE data, FTIR temperature series, 2D IR temperature series, NMR temperature series, and ITC data are uploaded to Zenodo and available at 10.5281/Zenodo.7662738.

## Supplementary Data

Supplementary Data are available at NAR Online.

## Acknowledgements

We thank Marco Todisco and Jakob Schauss for helpful discussions and feedback on the manuscript. This work was completed in part with resources provided by the University of Chicago Research Computing Center. We gratefully acknowledge computing time on the University of Chicago high-performance GPU-based cyberinfrastructure supported by the National Science Foundation under Grant No. DMR-1828629.

