## Supplemental Text and Figures for "Direct monitoring of the thermodynamics and kinetics of DNA and RNA dinucleotide dehybridization from gaps and overhangs"

#### Table of Contents

**S1. Spectroscopic characterization of AA and Primer/Helper dissociation……………......S2**
S1.1 FTIR-monitored titrations of $A_n$ binding at 1 °C
S1.2 FTIR temperature series
S1.3 2D IR temperature series
S1.4 Temperature-dependent $^1$H NMR of AA dissociation from a gap

**S2. Temperature-jump IR spectroscopy……………………………………………………..S12**

**S3. Thermodynamic modelling of IR and NMR temperature series data…………………S16**
S3.1 Independent model for $A_n$ dissociation and primer and helper dissociation
S3.2 Sequential unbinding model
S3.3 Fitting of $^1$H NMR temperature series data
S3.4 Global fitting of FTIR and 2D IR temperature series

**S4. Thermodynamics of free strand binding……………….…………………………..…S29**
S4.1 FTIR temperature series of GAAG, GAAAG, and GAAAAG
S4.2 ITC of free strand binding and $A_n$ binding to gaps
S4.3 Length-scaling of hybridization thermodynamics and kinetics

**S5. Molecular dynamics simulations…………………………………………………........S35**
S5.1 Structural comparison of gap single-strand regions and free single-strands
S5.2 AA-gap complex and duplex structural parameters
S5.3 Calculation of relative WCF to anti-HG base pair stability with enhanced sampling

**S6. Estimation of diffusion-limited rate constant for AA binding to gap……………….…S44**

#### S1. Spectroscopic characterization of AA and primer/helper dissociation

##### S1.1 FTIR-monitored titrations of $A_n$ binding at 1 °C

Short oligonucleotides ($A_n$, with $n$ = 2,3,4) were titrated against gap and overhang templates in order to determine the fraction of bound $A_n$ in a 1:1 mixture used for temperature-dependent measurements. Titrations were performed at 1 °C, where the fraction of bound primer and helper is assumed to be ~1, and $A_n$ association can be modeled as a two-state equilibrium.

$$\mathrm{X} + \mathrm{A_n} \rightleftharpoons \mathrm{X:A_n} \tag{S1}$$

Here X is the template and X:$A_n$ is the bound $A_n$-template complex. Upon addition of $A_n$, a reduction in absorbance is observed near 1665 $cm^{-1}$ with a corresponding gain centered near 1695 $cm^{-1}$ (Fig. S1). These changes come from thymine (T) and guanine (G) carbonyl bands and are consistent with an both an increase A:T base pairing and stacking of guanine nucleotides.(1) The second component from singular value decomposition (SVD) component over the frequency window from 1650 to 1720 $cm^{-1}$ was used to describe the concentration of X:$A_n$ and was fit to a general two-state binding expression (eq. S2).(2)

$$[\mathrm{X:A_n}] = 0.5\left(K_\mathrm{d} + c_{\mathrm{A_n}} + c_\mathrm{Temp} - \sqrt{\left(K_\mathrm{d} + c_{\mathrm{A_n}} + c_\mathrm{Temp}\right)^2 - 4c_{\mathrm{A_n}}c_\mathrm{Temp}}\right) \tag{S2}$$

$$\text{where } c_{\mathrm{A_n}} = [\mathrm{X:A_n}] + [\mathrm{A_n}] \text{ and } c_\mathrm{Temp} = [\mathrm{X:A_n}] + [\mathrm{X}]$$

An initial measurement was made with [$A_n$] = 0, so the minimum of the 2nd SVD component was set to 0 prior to fitting. In practice, an amplitude scaling factor and $K_d$ were used as fit parameters and the normalized fit was used to represent fraction of bound X:$A_n$.

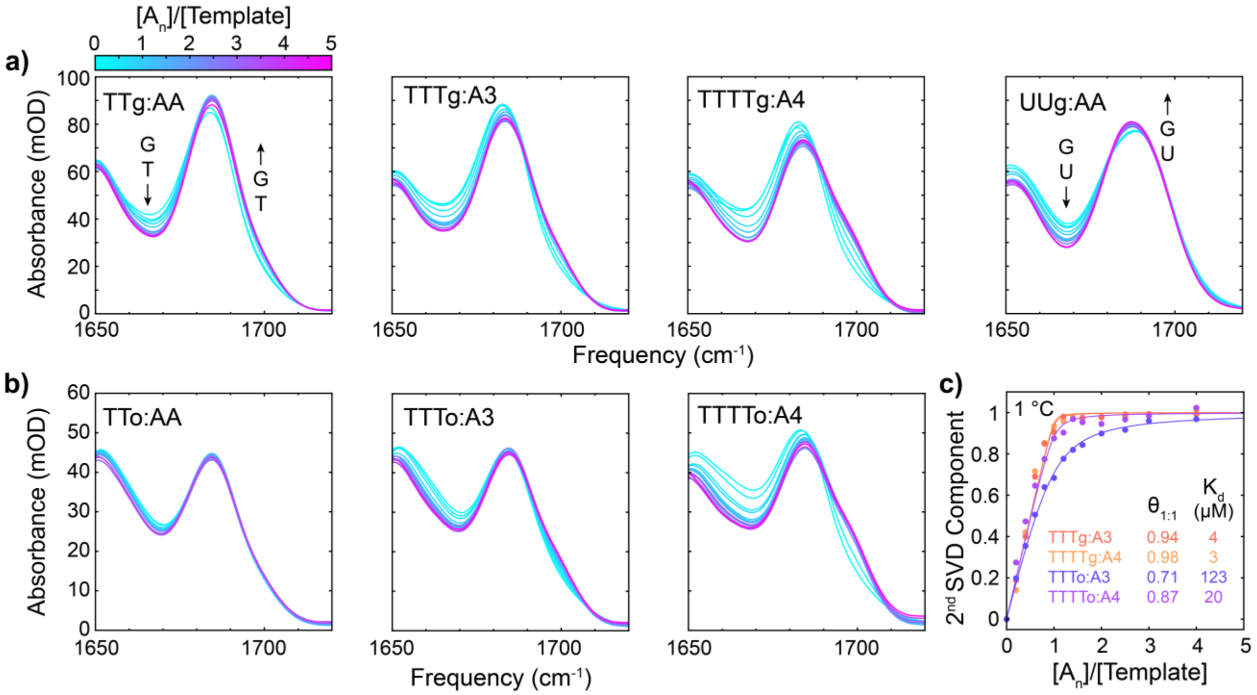


**Figure S1. FTIR-monitored titrations of $A_n$ binding at 1 °C. (a)** FTIR spectra as a function of molar ratio between $A_n$ and their gap template. The template is at a concentration of 1 mM for all spectra. **(b)** FTIR spectra for titrations with overhang templates. **(c)** Titration curves for TTTg:A3, TTTTg:A4, TTTo:A3, and TTTTo:A4 determined from the second component of singular value decomposition (SVD) of FTIR-monitored titration data. SVD was performed over the 1650 to 1720 $cm^{-1}$ spectral range. Solid lines correspond to fits to a two-state binding model (eq. S2). Fraction of X bound to $A_n$ at a 1:1 molar ratio ($\theta_{1:1}$) and dissociation constants ($K_d$) extracted from the fits are listed.

**Table S1.** Thermodynamic parameters for $A_n$ dissociation at 1 °C determined from FTIR-monitored titrations[a,c]

| Complex | $K_d$ (μM) | $\boldsymbol{\theta_{1:1}}$[b] |
|---|---|---|
| **TTg:AA** | 12 ± 10 | 0.89 ± 0.06 |
| **TTTg:A3** | 4 ± 4 | 0.94 ± 0.07 |
| **TTTTg:A4** | 3 ± 3 | 0.98 ± 0.08 |
| **UUg:AA** | 147 ± 40 | 0.70 ± 0.07 |
| **TTo:AA** | 330 ± 150 | 0.54 ± 0.18 |
| **TTTo:A3** | 123 ± 38 | 0.71 ± 0.08 |
| **TTTTo:A4** | 20 ± 16 | 0.87 ± 0.11 |

[a]All values determined in deuterated pH* 6.8 400 mM SPB buffer with $c_{\text{Temp}}$ = 1 mM. [b]Fraction of X bound to $A_n$ at a 1:1 molar ratio. [c]Errors determined from 95% confidence intervals of titration curve fit parameters.

##### S1.2 FTIR temperature series

Additional temperature-dependent linear spectral changes overlap with the sigmoidal dissociation transitions of interest in FTIR temperature series. These linear changes are particularly significant in the region from 1600 to 1700 $cm^{-1}$, which contains vibrational bands from all of the nucleobases, whereas the guanine and adenine ring modes from 1550 to 1600 $cm^{-1}$ exhibit minimal interference. In general, such linear spectral changes may arise from both structural and non-structural effects of the nucleic acid. For example, solvation dynamics and the strength of hydrogen bonding between the nucleobase and solvent are temperature-dependent and lead to spectral changes in the FTIR spectra of free nucleobases.(3) Such changes are better characterized for the Amide I band of proteins.(4) Another non-structural source of absorbance changes may result from systematic error in subtraction of the $D_2O$ bend-libration combination band background from the raw data. However, linear temperature-dependent spectral changes may also arise from noncooperative weakening or loss of base pairing that is most commonly observed from terminal base pairing fraying in oliognucleotides.(1,5) In a two-state treatment of spectroscopic melting curves, these linear changes are subtracted from melting curves through baseline fitting,(6,7) and the presence of multiple melting transitions in this work complicates baseline removal.

It is worth noting that an additional non-linear temperature-dependent spectral change is observed from 1 to 25 °C in TTo as shown at 1625, 1665, and 1695 $cm^{-1}$ (Fig. S2). It is possible that this change is reporting of the unfolding a hairpin formed with contacts between the 5′-GA-3′ end of the overhang and 5′-TC-3′ segment at the center of the template strand, requiring displacement of a G:C base pair between primer and template strands. Due to overlapping temperature range of apparent hairpin unfolding in TTo:AA and AA dissociation in TTo:AA, it is unclear what population of hairpin exists at low temperature in TTo:AA. The clear signatures of AA dissociation in temperature-dependent FTIR, 2D IR, and T-jump IR measurements suggest that the AA-overhang complex is more populated than the hairpin (Figs. S2, S8, S12).

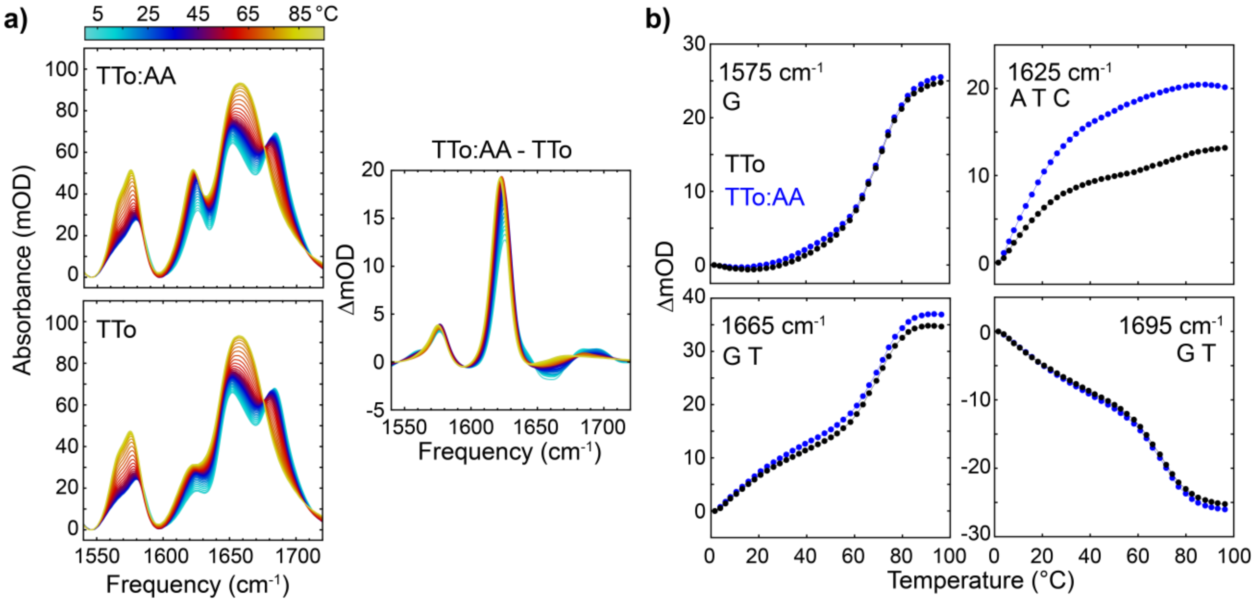


**Figure S2. Temperature-dependent FTIR spectra of DNA overhang templates. (a)** FTIR temperature series for TTo and TTo:AA from 1 to 96 °C. The right panel shows the differences spectra between TTo:AA and TTo at each temperature. **(b)** Change in absorbance as a function of temperature at select FTIR frequencies for TTo (black) and TTo:AA (blue).

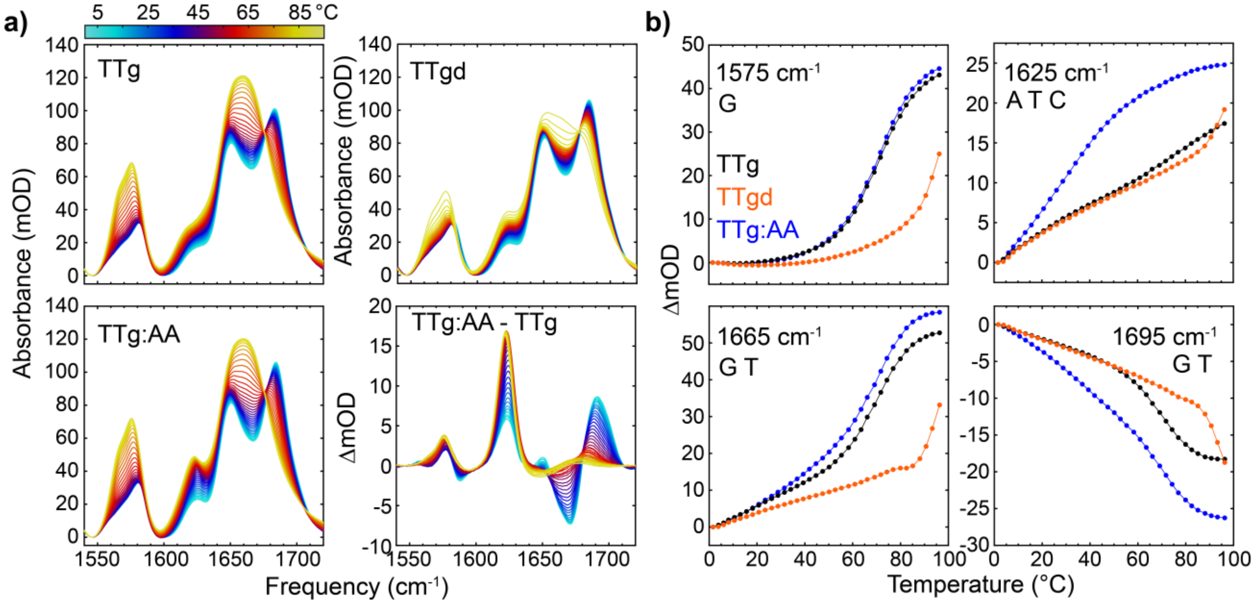


**Figure S3. Temperature-dependent FTIR spectra of DNA gap templates. (a)** FTIR temperature series for TTg, TTg:AA, and a fully complementary duplex 5′-GCGGCGAAGGCGGC-3′/5′-GCCGCCTTCGCCGC-3′ (TTgd) from 1 to 96 °C. The bottom-right panel shows the differences spectra between TTg:AA and TTg at each temperature. **(b)** Change in absorbance as a function of temperature at select FTIR frequencies for TTg (black), TTg:AA (blue), and TTgd (orange).

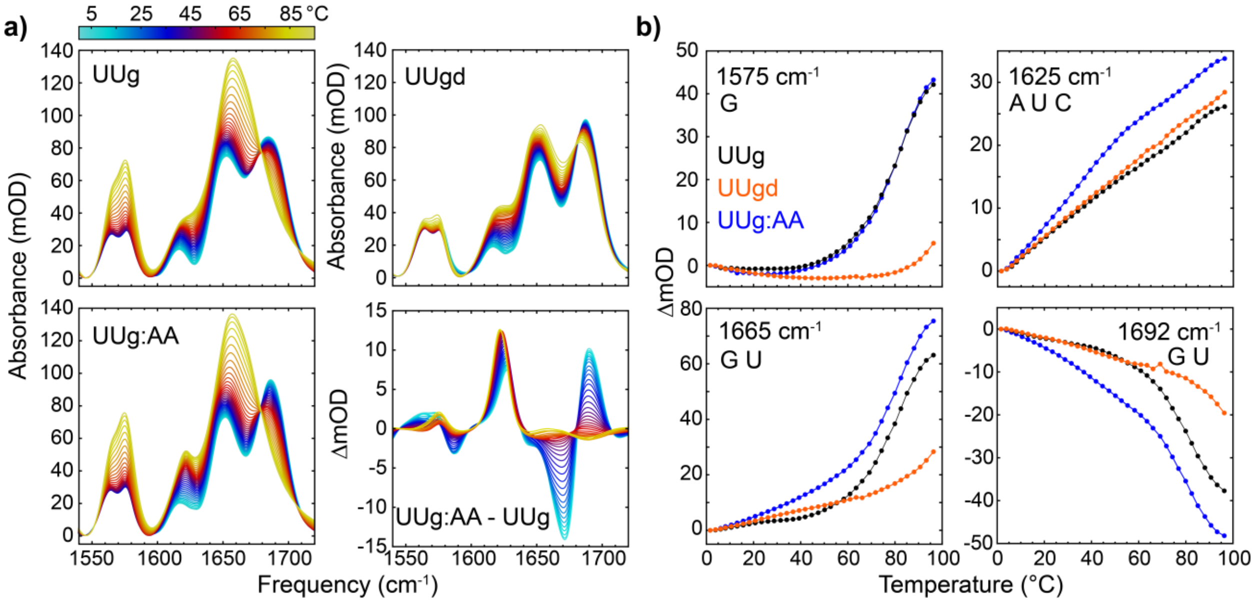


**Figure S4. Temperature-dependent FTIR spectra of RNA gap templates. (a)** FTIR temperature series for UUg, UUg:AA, and a fully complementary duplex 5′-GCGGCGAAGGCGGC-3′/5′-GCCGCCUUCGCCGC-3′ (UUgd) from 1 to 96 °C. The bottom-right panel shows the differences spectra between UUg:AA and UUg at each temperature. **(b)** Change in absorbance as a function of temperature at select FTIR frequencies for UUg (black), UUg:AA (blue), and UUgd (orange).

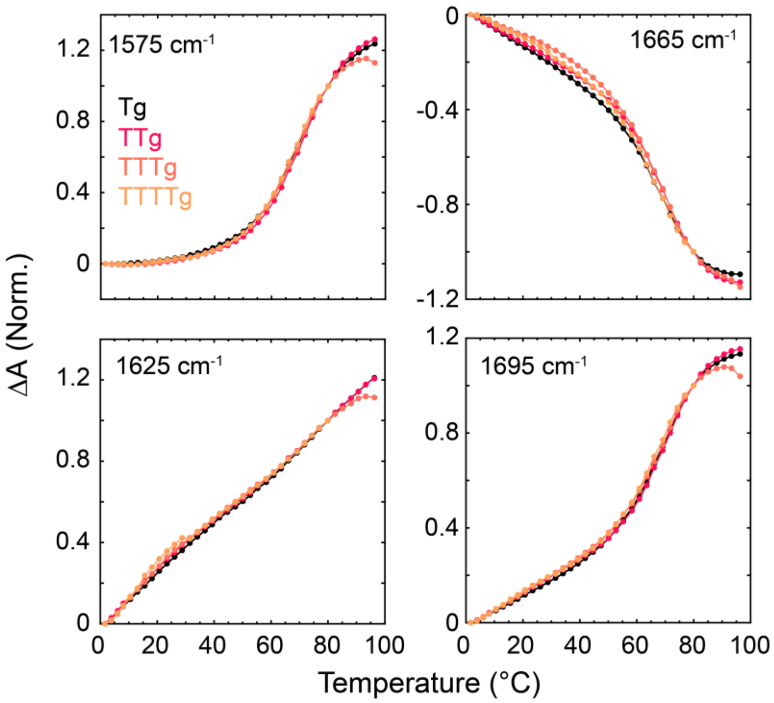

**Figure S5. Gap size effect on primer and helper melting transition.** Normalized temperature-dependent FTIR absorption change relative to 1 °C (ΔA) at 1575, 1625, 1665, and 1695 $cm^{-1}$ for DNA gap templates. Tg corresponds a single T gap with identical duplex regions to TTg, TTTg, and TTTTg. The temperature-dependent ΔA is nearly identical for all gap templates.

##### S1.3 2D IR temperature series

Two-dimensional IR (2D IR) temperature series were acquired as an additional measurement of AA dissociation and primer and helper dissociation. Overall, the 2D IR spectra contain information on vibrational anharmonicity, lineshape, and coupling between vibrations that cannot be extracted from FTIR spectra, and these factors provide added sensitivity to changes in base pairing and stacking structure.(1,8-10) Another important factor is that the 2D IR signal is proportional to the fourth order of the transition dipole moment ($\mu_i^4$) for each oscillator *i*. As a result, the third-order signal of the $D_2O$-bend-libration combination band, which must be subtracted from raw FTIR spectra, has negligible third-order signal amplitude relative to the 2D IR signal of the nucleic acid. Further details of 2D IR spectroscopy have been described elsewhere.(11,12)

Fig. S6a shows 2D IR spectra of TTg:AA, TTo:AA, and UUg:AA at 1, 43, and 91 °C. All peaks present as positive (red-yellow) and negative (blue) doublets corresponding to ground-state bleach (GSB) and excited-state absorption (ESA) transitions. GSB peaks along the diagonal correspond to the same fundamental vibrational transitions observed in FTIR whereas cross-peaks report on coupling between vibrations. Difference spectra between 43 and 1 °C as well as 91 and 43 °C are shown in Fig. S6b and roughly correspond to the spectral changes of AA and template dissociation, respectively. The 43 – 1 °C spectra are similar for each sequence and characterized by an increase in amplitude of the diagonal guanine (1575 $cm^{-1}$) and adenine (1625 $cm^{-1}$) ring mode peaks as well as a decrease in amplitude of the overlapping guanine and thymine carbonyl bands (1680-1700 $cm^{-1}$), and these changes are consistent with those observed in the FTIR spectra. A reduction in cross-peak amplitude between 1690 and 1650 $cm^{-1}$, 1690 and 1625 $cm^{-1}$, and 1690 and 1585 $cm^{-1}$ bands are observed. The first two sets of cross-peaks may arise from coupling between thymine vibrational modes on the same nucleobase or from interbase couplings between guanine and cytosine whereas the third cross-peak comes from coupling between guanine carbonyl and ring modes on the same nucleobase.(1,8,13) There is also an increase in amplitude of the cross-peak between guanine carbonyl and ring modes at 1665 and 1575 $cm^{-1}$.

Figs. S7-S8 compare temperature-dependent 2D IR spectral changes of AA dissociation with those from template (black) and complement duplexes (orange) over the 1 to 43 °C. The signal change along the diagonal at 1625 $cm^{-1}$ is significantly lower or negligible in the template and complement duplexes relative to the AA-template complexes. However, the templates and complement duplexes do show signal change that indicate a loss or weakening of G:C base pairing or G stacking. For TTg, UUg, and TTo, the weakening of G:C base pairing may indicate a small increase in unbound primer and helper population or an increase in terminal base pair fraying. TTgd and UUgd have negligible dehybridized population at 43 °C and their spectral changes are likely dominated by an increase of terminal base pair fraying. The temperature-dependent integrated signal change over select regions are shown for each sequence. In contrast to the FTIR data (Figs. S2-S4), there are only minor changes in the template and complement duplexes relative to the AA-template complexes in the 1625 $cm^{-1}$ region.

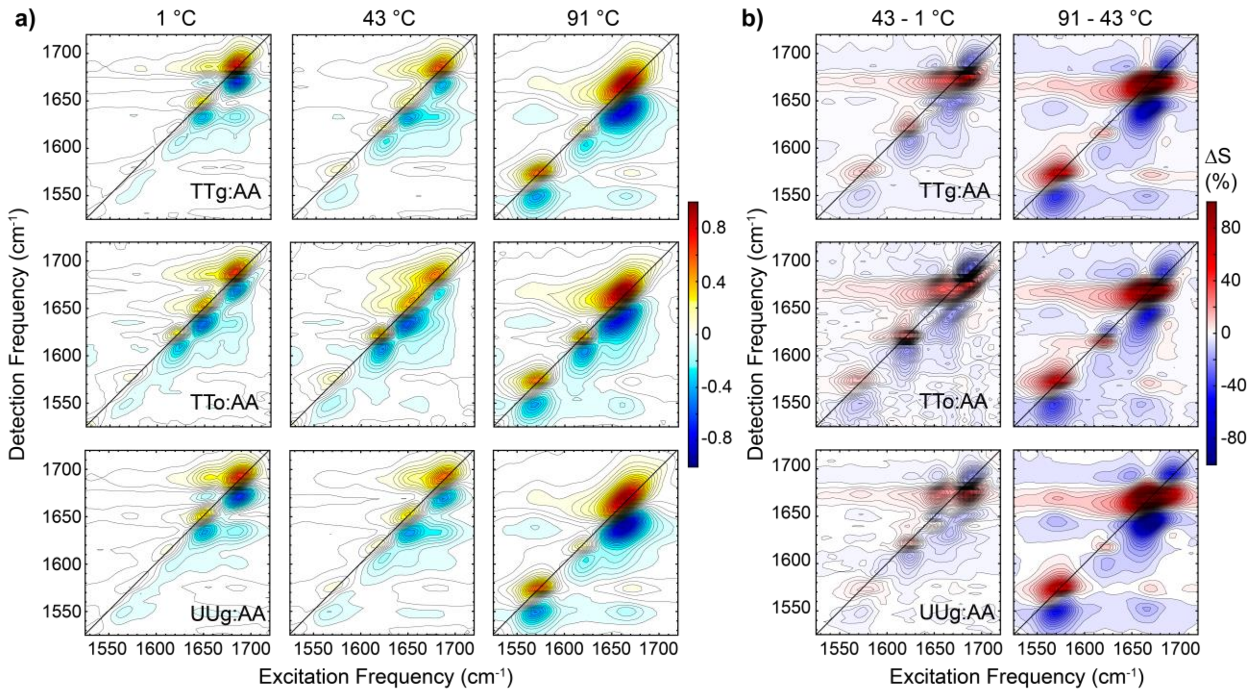


**Figure S6. Temperature-dependent 2D IR spectroscopy of AA and template dissociation. (a)** 2D IR spectra of TTg:AA, TTo:AA, and UUg:AA at 1, 43, and 91 °C taken with parallel (ZZZZ) pulse polarization and at a waiting time of 150 fs. Yellow-red amplitude is positive and blue is negative. 25 contours are plotted with uniform spacing. The solid line indicates diagonal frequencies. **(b)** Difference spectra between (left) 43 and 1 °C and (right) 91 and 43 °C for each sequence. Data are plotted in percent change relative to the maximum of the lower temperature spectrum (ΔS). 25 contours are plotted with uniform ΔS=3.6% spacing.

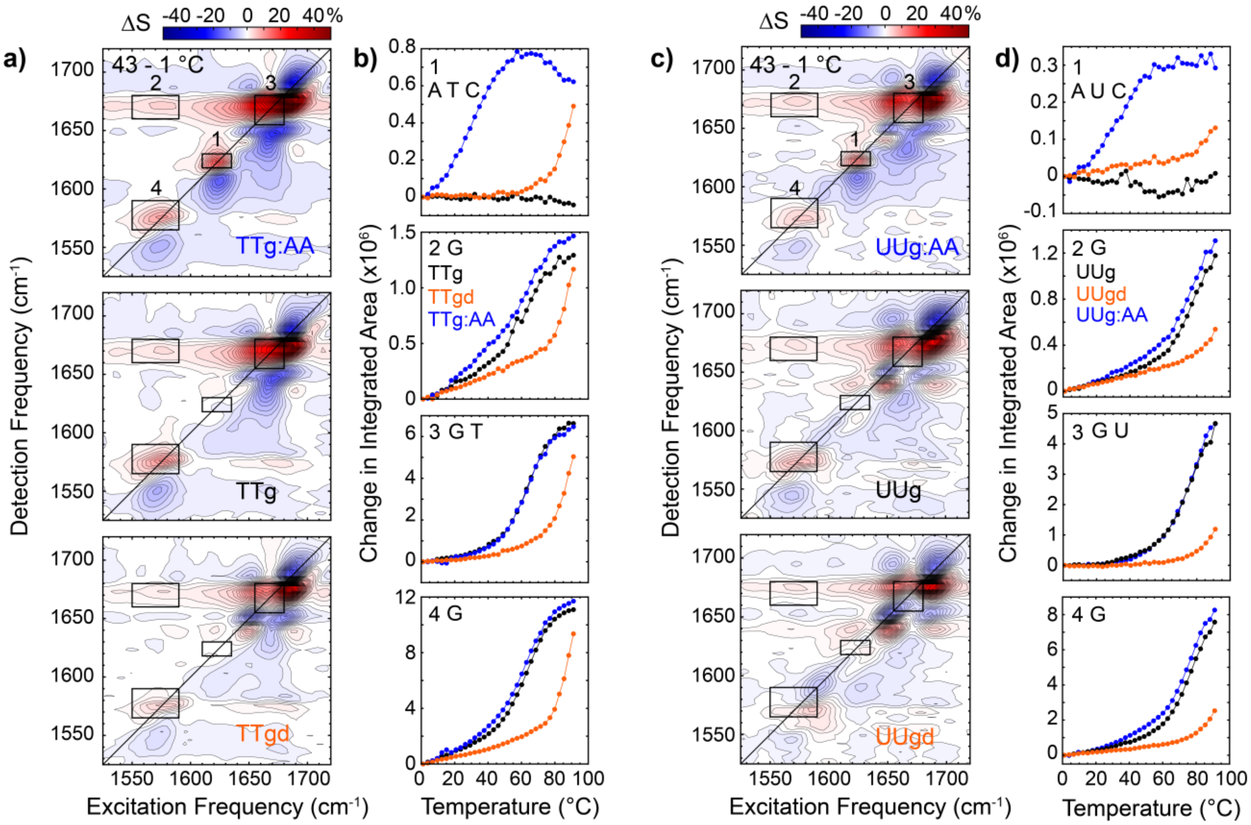


**Figure S7. Temperature-dependent 2D IR spectral changes of DNA and RNA gap templates. (a)** 2D IR difference spectra between 43 and 1 °C for TTg:AA, TTg, and TTgd. **(b)** Temperature-dependent integrated amplitudes over the regions marked in **(a)** plotted relative to the spectrum at 1 °C. **(c-d)** Corresponding temperature-dependent 2D IR data for UUg:AA, UUg, and UUgd.

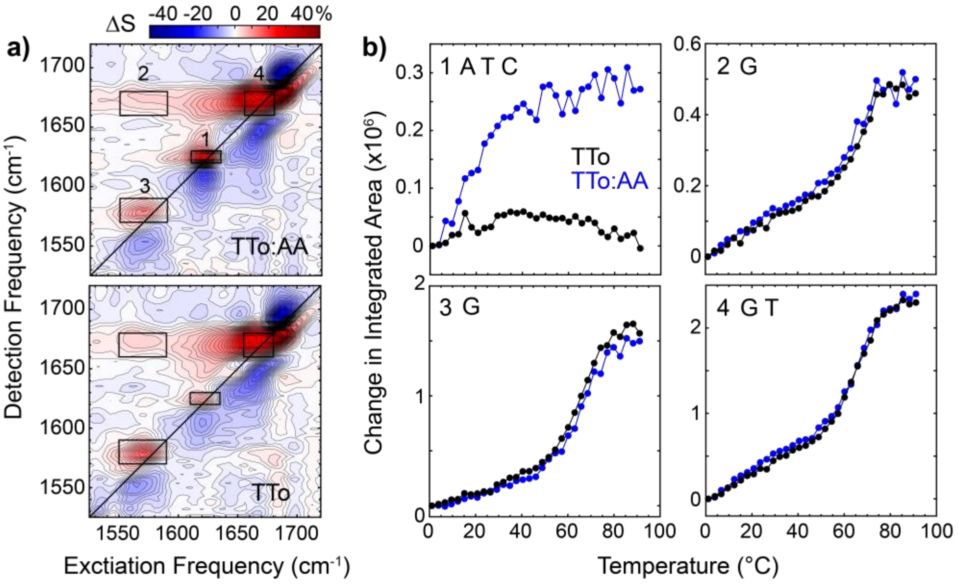


**Figure S8. Temperature-dependent 2D IR spectral changes of TTo and TTo:AA. (a)** 2D IR difference spectra between 43 and 1 °C for TTo:AA, TTo. **(b)** Temperature-dependent integrated amplitudes over the regions marked in **(a)** plotted relative to the spectrum at 1 °C.

#### S1.4 Temperature-dependent $^1$H NMR of AA dissociation from a gap

We performed temperature-dependent $^1$H NMR measurements to directly compare with our IR measurements of AA dissociation in TTg:AA. $^1$H NMR temperature series have previously been used to extract nucleic acid binding thermodynamics and contain base-specific information in short oligonucleotides.(14,15) Numerous resonances are sensitive to dissociation of AA and/or primer and helper dissociation, and a few signals may be used to selectively track each process. The H2 and H8 protons of adenine specifically probe the AA dissociation equilibrium while H5 and H6 protons of cytosine will primarily be sensitive to primer and helper dissociation. Many of the peaks are broad and overlap at low temperature, likely due to slow tumbling of the AA-gap complex, and become more narrow and separated as the temperature increases. Due to fast AA association and dissociation, chemical shifts of the four aromatic adenine protons from 7.6 to 8.4 ppm report on the relative population of bound and unbound states (Fig. S9). From 10 to 50 °C, identification of the aromatic adenine peaks is hindered by overlap with resonances from aromatic protons on guanine, thymine, and cytosine, but the chemical shift of two adenine peaks (labeled A3 and A4) can be determined over most of the temperature range through comparison with temperature-dependent spectra of free AA and TTg. The temperature-dependence of A3 and A4 chemical shift follows a single sigmoidal transition that reports on the AA binding equilibrium and exhibits a midpoint and width similar to the IR-monitored transition (Fig. 2). Due to the congestion of aromatic peaks from 7.6 to 8.4 ppm, we use the H5 cytosine protons located from 5.2 to 6.2 ppm to monitor the primer and helper binding equilibrium. Two H5 cytosine signals (C1 and C2), that appear insensitive to AA dissociation, are distinguished from H1′ peaks of the deoxyribose groups by TOCSY cross-peaks to H6 cytosine signals (Figs. S9-S10) and exhibit sigmoidal temperature-dependent trends in chemical shift that report on primer and helper dissociation.

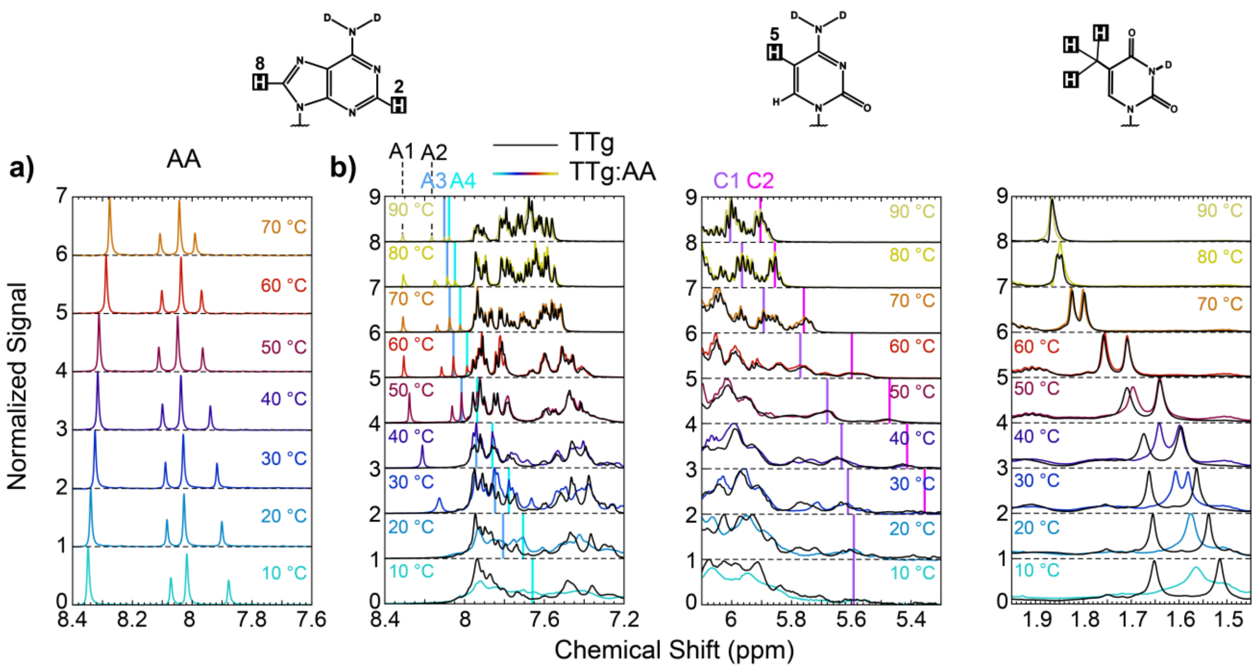


**Figure S9. Temperature-dependent $^{1}$H NMR spectra of free AA and gap template. (a)** Normalized aromatic proton spectra of AA. **(b)** Normalized $^{1}$H NMR spectra of TTg:AA from 10 °C to 90 °C. (Left) Frequency window from 8.4 to 7.6 ppm that contains H8/H2 signals of adenine, H8 signals of guanine, and H6 signals of thymine and cytosine. Each adenine proton signal is labeled A1-A4 in the 90 °C spectrum, and the A3 and A4 chemical shift values are marked at each temperature with dark and light blue vertical lines, respectively. The spectra of TTg are shown in black and those of TTg:AA are colored. A3 and A4 chemical shifts are indicated with vertical lines. (Center) Frequency window from 6.2 to 5.2 ppm that contains H1′ signals from the 2′-deoxyribose moiety of each nucleotide and H5 signals from cytosine. Two cytosine H5 signals are denoted C1 and C2 and marked with purple and magenta vertical lines, respectively, at each temperature. (Right) Frequency window from 1.45 to 1.95 ppm window showing the methyl protons of thymine. All oligonucleotides are present at 1 mM concentration in pH* 6.8 400 mM SPB in $D_2O$.

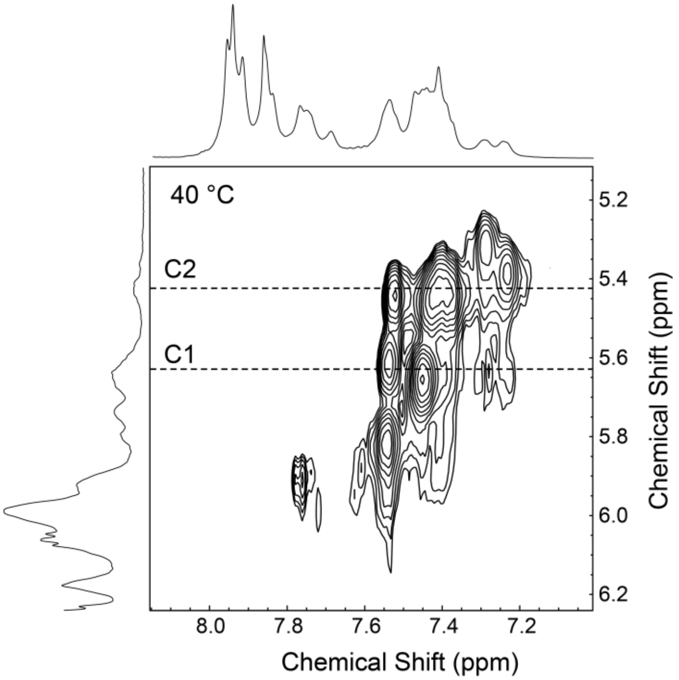


**Figure S10. Assignment of cytosine H5 protons with $^1$H-$^1$H Total correlated spectroscopy (TOCSY) measurements in TTg:AA.** TOCSY cross-peaks between the 7.2 – 8.0 ppm and 5.1 – 6.2 ppm frequency windows due to *J*-coupling between cytosine H5 (5.1-6.0 ppm) and H6 (7.2 – 8.0 ppm) atoms. $^1$H NMR spectra over each region are shown to the left and top. The TOCSY cross-peaks are used to differentiate cytosine H5 from H1′ resonances.

#### S2. Temperature-jump IR spectroscopy

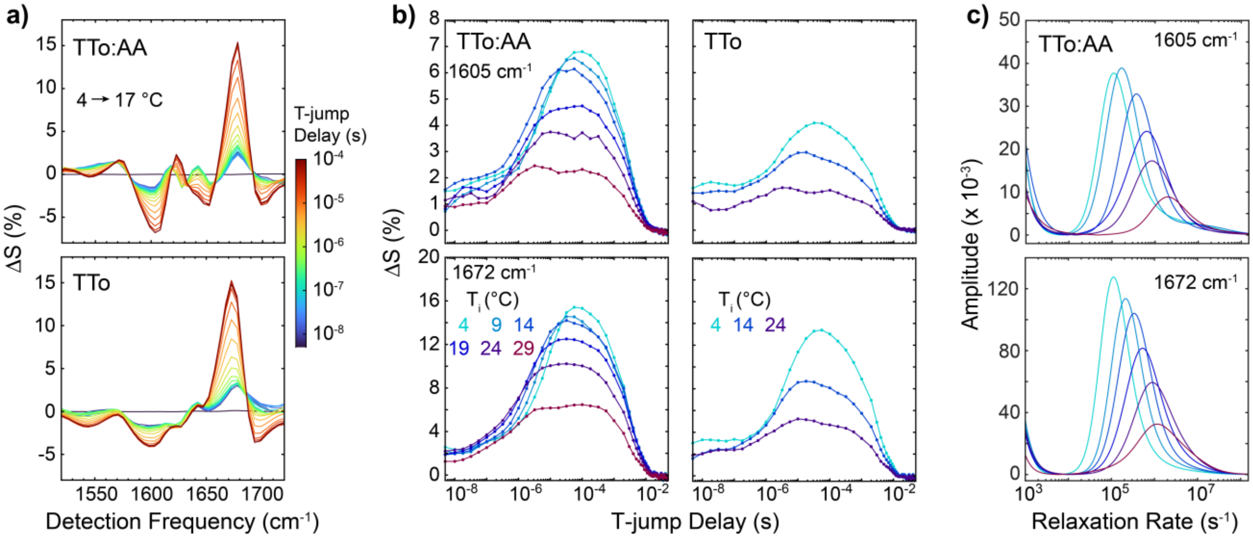


**Figure S11. T-jump measurements of AA dissociation in TTo:AA. (a)** t-HDVE spectra of TTo:AA (top) and TTo (bottom) for a T-jump from 4 to 17 °C. Spectra are plotted as the change relative to the maximum of the initial temperature spectrum ($\Delta S(t) = S(t)/\max(S(T_i))$) **(b)**

t-HDVE responses probed at 1605 (top) and 1672 cm$^{-1}$ (bottom) for multiple temperatures indicated on the plot. **(c)** Rate-domain traces at each temperature for TTo:AA probed at 1605 and 1672 cm$^{-1}$ obtained from MEM-iLT.(16,17)

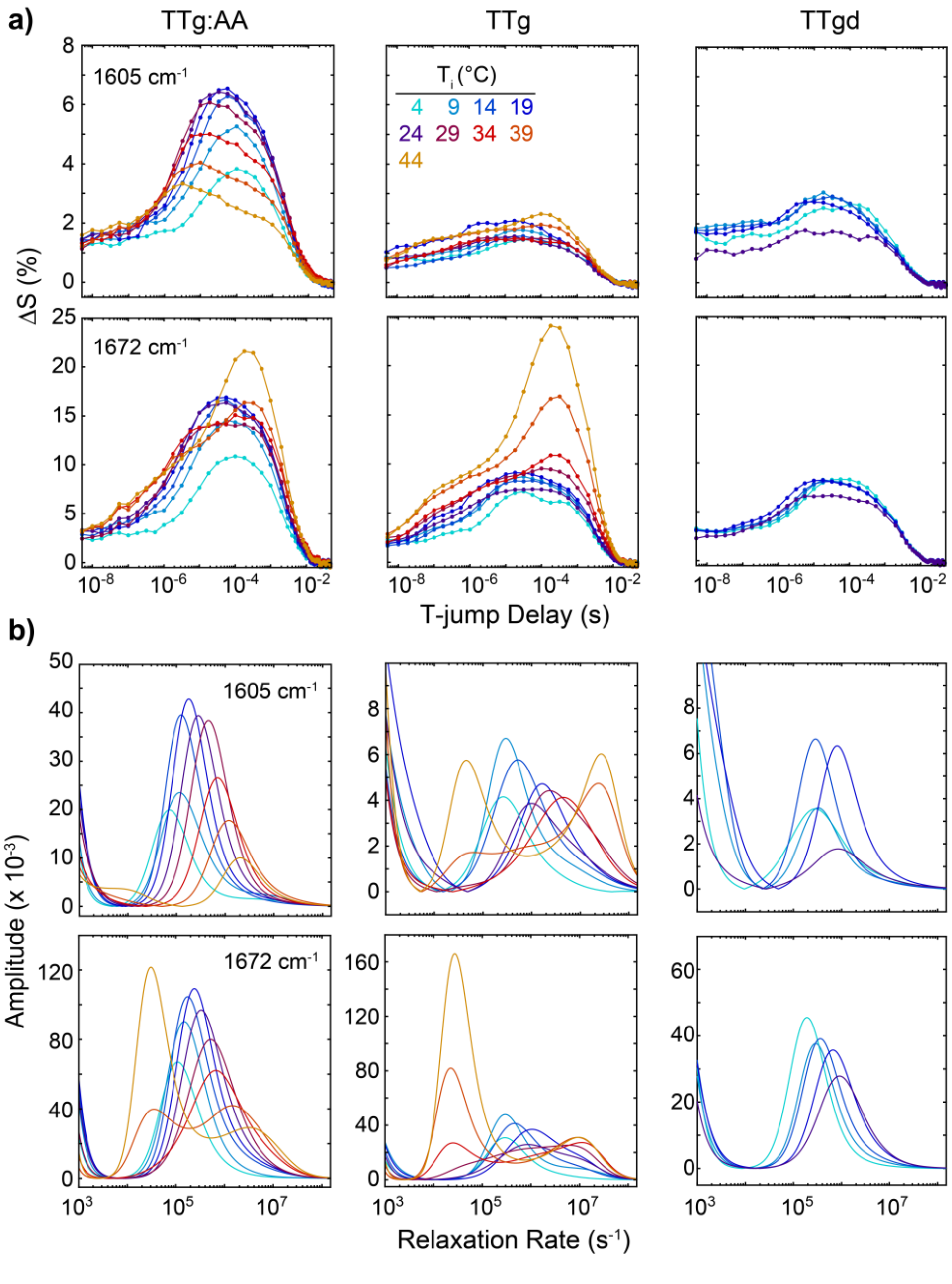


**Figure S12**. **Temperature-dependent T-jump measurements of DNA gap sequences. (a)** t-HDVE time-domain responses of TTg:AA, TTg, and TTgd probed at 1605 (top) and 1672 cm$^{-1}$ (bottom) for multiple temperatures indicated on the plot. **(b)** Rate-domain traces for each sequence and temperature probed at 1605 and 1672 cm$^{-1}$ obtained from MEM-iLT.

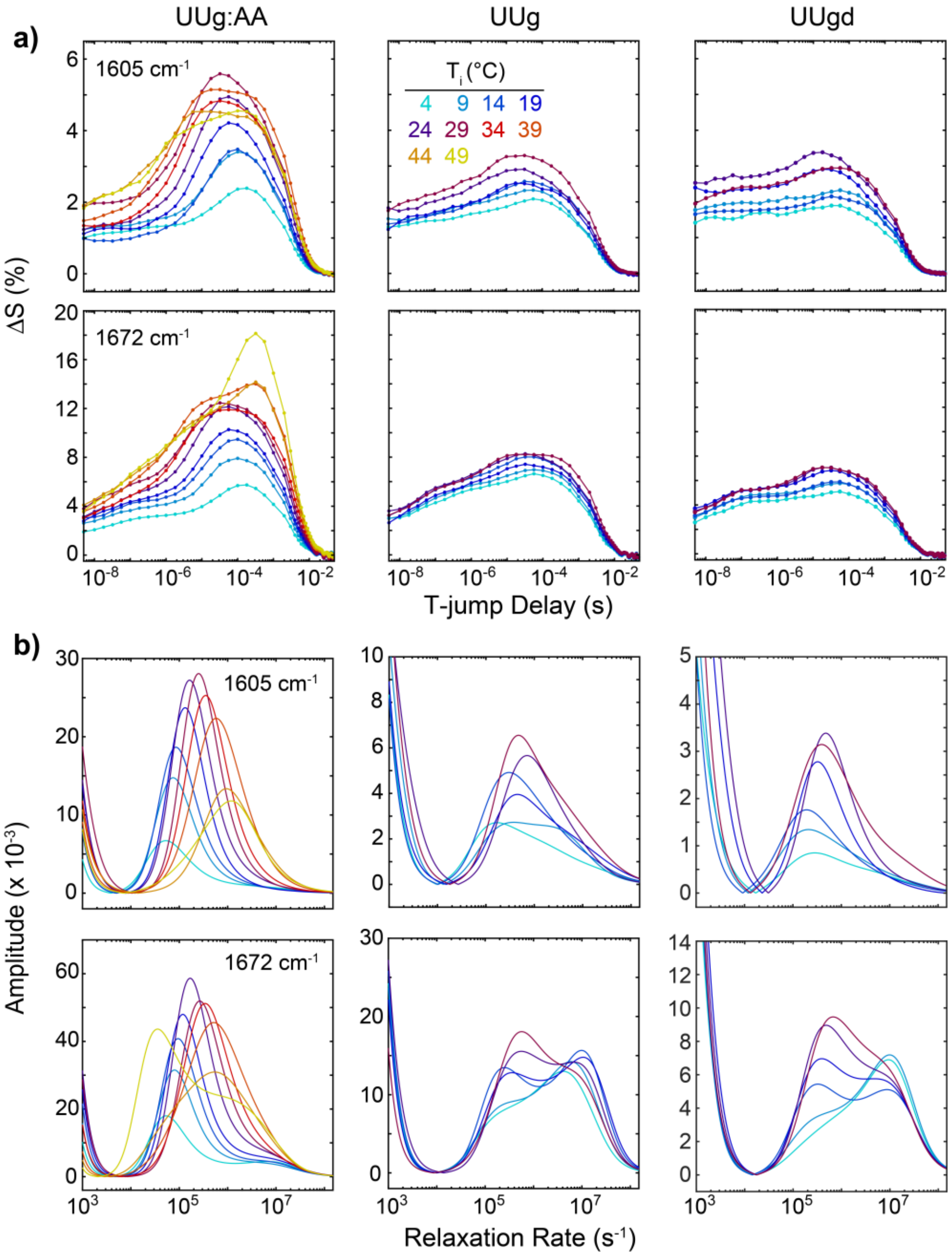


**Figure S13**. **Temperature-dependent T-jump measurements of RNA gap sequences. (a)** t-HDVE time-domain responses of UUg:AA, UUg, and UUgd probed at 1605 (top) and 1672 $cm^{-1}$ (bottom) for multiple temperatures indicated on the plot. **(b)** Rate-domain traces for each sequence and temperature probed at 1605 and 1672 $cm^{-1}$ obtained from MEM-iLT.

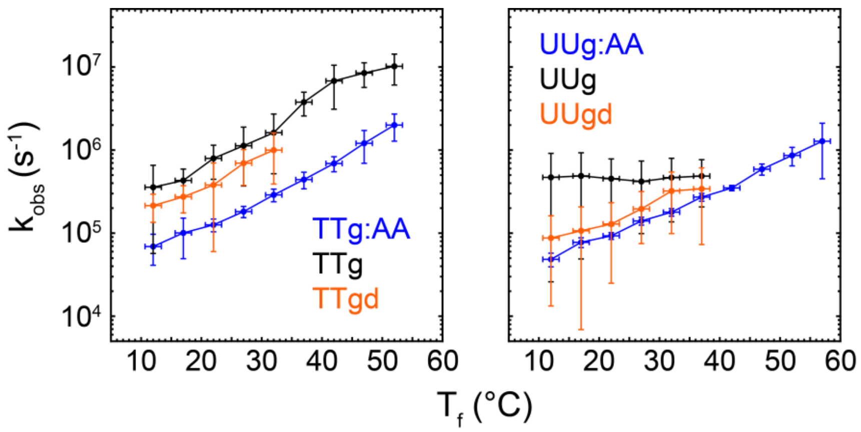


**Figure S14**. **Observed T-jump rate for AA dissociation and template response.** Temperature-dependent observed rates for (left) DNA and (right) RNA gap and duplex sequences obtained from amplitude-weighted mean of rate-domain spectra. Rates for TTg, TTgd, UUg, and UUgd were determined over the 1585-1690 $cm^{-1}$ window while those for TTg:AA and UUg:AA were determined from the 1585-1610 $cm^{-1}$. Vertical error bars indicate standard deviation over the specified frequency range of the rate spectra and horizontal error bars correspond to the measured standard deviation in T-jump magnitude.

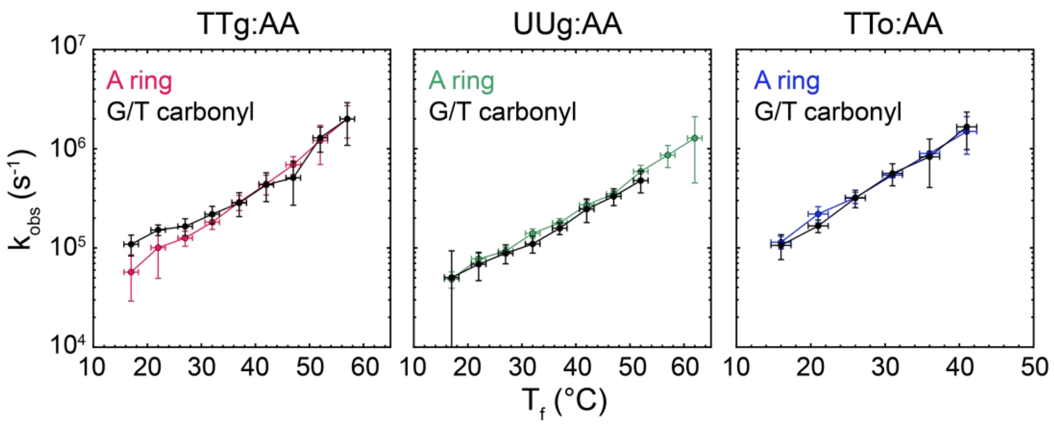


**Figure S15**. **Comparison of observed T-jump rate obtained from different spectral regions.** Observed rates are determined from the amplitude-weighted mean of rate-domain spectra for TTg:AA, UUg:AA, and TTo:AA across the adenine (A) ring mode (1585-1610 $cm^{-1}$) and guanine (G) and thymine (T) carbonyl mode (1660 to 1685 $cm^{-1}$) spectral regions. Vertical error bars indicate standard deviation over the specified frequency range of the rate spectra and horizontal error bars correspond to the measured standard deviation in T-jump magnitude.

#### S3. Thermodynamic modelling of IR and NMR temperature series data

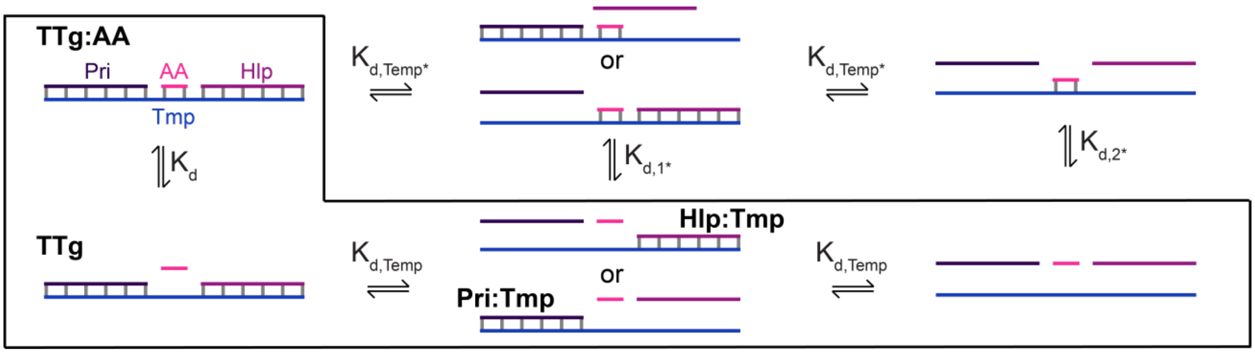


**Figure S16. Schematic of coupled binding equilibria for AA-template complexes using TTg:AA as an example.** Direct binding of AA with the template strand (Tmp), primer:template complex, and helper:template complex is negligible due to the much greater binding stability of the primer (Pri) and helper (Hlp) relative to AA. The full unbinding of the complex proceeds first by AA dissociation (bottom-left) followed by dissociation of Pri and Hlp at higher temperatures.

The $A_n$-template complex (X:$A_n$) formed between a short oligonucleotide ($A_n$) and the gap or overhang template (X) exhibits two dissociation equilibria as a function of temperature. In general, there may be multiple thermodynamic paths for unbinding where either $A_n$, primer, or helper strands dissociate first (Fig. S16). For all complexes studied in this work, $K_{\mathrm{d}} >> K_{\mathrm{d,Temp1*}}$ & $K_{\mathrm{d,Temp2*}}$ over the measured temperature range due to use of G:C-rich 6-mer primer (Pri) and helper (Hlp) strands that bind to the template strand (Tmp) much more strongly than $A_n$. For the same reason, $K_{\mathrm{d,1*}}$ & $K_{\mathrm{d,2*}} >> K_{\mathrm{d,Temp1}}$ & $K_{\mathrm{d,Temp2}}$, and $A_n$ binding to Tmp will be negligible population over the measured temperature range. Additionally, primer and helper strands have nearly identical binding stability such that $K_{\mathrm{d,Temp1}} \sim K_{\mathrm{d,Temp2}} \sim K_{\mathrm{d,Temp1*}} \sim K_{\mathrm{d,Temp2*}} = K_{\mathrm{d,Temp}}$. Due to these conditions, we can treat the dissociation thermodynamics using just $K_{\mathrm{d}}$ & $K_{\mathrm{d,Temp}}$ equilibria.

$$\mathrm{X{:}A}_n \xrightleftharpoons{K_{\mathrm{d}}} \mathrm{X} + \mathrm{A}_n \qquad \text{(S3a)}$$

$$\mathrm{X} \xrightleftharpoons{K_{d,\mathrm{Temp}}} \mathrm{Pri{:}Tmp + Hlp} \xrightleftharpoons{K_{d,\mathrm{Temp}}} \mathrm{Pri + Hlp + Tmp} \quad \text{(Gap)} \qquad \text{(S3b)}$$

$$\mathrm{X} \xrightleftharpoons{K_{d,\mathrm{Temp}}} \mathrm{Hlp{:}Tmp + Pri} \xrightleftharpoons{K_{d,\mathrm{Temp}}} \mathrm{Pri + Hlp + Tmp} \quad \text{(Gap)} \qquad \text{(S3c)}$$

$$\mathrm{X} \xrightleftharpoons{K_{\mathrm{d,Temp}}} \mathrm{Pri + Tmp} \quad \text{(Overhang)} \qquad \text{(S3d)}$$

#### S3.1 Independent model of $A_n$ dissociation and primer and helper dissociation

The $A_n$ and primer and helper melting curves are typically well-separated in temperature ($K_d >> K_{d,Temp}$) for the sequences studied in this work. Therefore, it may be adequate to treat these processes as independent equilibria with the following equilibrium constants.

$$K_d = \frac{[X][A_n]}{[X{:}A_n]} \tag{S4a}$$

$$K_{d,Temp} = \frac{[Pri{:}Tmp][Hlp]}{[X]} = \frac{[Pri][Tmp]}{[Pri{:}Tmp]} = \frac{[Pri][Hlp{:}Tmp]}{[X]} = \frac{[Hlp][Tmp]}{[Hlp{:}Tmp]} \text{ (Gap)} \tag{S4b}$$

$$K_{d,Temp} = \frac{[Pri][Tmp]}{[X]} \text{ (Overhang)} \tag{S4c}$$

$A_n$ dissociation is described using a two-state model for non-self-complementary oligonucleotides where $K_d$ is related to the fraction of bound $A_n$ ($\theta_{A_n}$).

$$K_d = \frac{\left(1-\theta_{A_n}\right)^2 c_{A_n}}{2\theta_{A_n}} \tag{S5}$$

$$\text{where, } \theta_{A_n} = \frac{2[X{:}A_n]}{c_1} \quad \text{and } c_1 = 2[X{:}A_n]+[X]+[A_n]$$

Eq. S5 assumes a 1:1 molar ratio between X and $A_n$, which is most valid at low temperature when [X] >> [Pri], [Hlp], [Tmp]. Then, eq. S5 can be arranged to express $\theta_{A_n}$ in terms of $K_d$.

$$\theta_{A_n} = 1 + \frac{K_d - \sqrt{K_d^2 + 2c_1 K_d}}{c_1} \tag{S6}$$

The temperature-dependence of $\theta_{A_n}$ and $K_d$ are determined by the enthalpy ($\Delta H_d^\circ$) and entropy ($\Delta S_d^\circ$) for dissociation of $A_n$.

$$K_d(T) = \exp\left[-\frac{\Delta H_d^\circ}{RT} + \frac{\Delta S_d^\circ}{R}\right] \tag{S7}$$

The dissociation temperature ($T_m$), which is defined as the temperature where $\theta_{A_n}$ = 0.5, can be re-cast in terms of $\Delta H_d^\circ$ and $\Delta S_d^\circ$.

$$T_m = \frac{\Delta H_d^\circ}{\Delta S_d^\circ - R\ln\left(c_1/4\right)} \tag{S8}$$

We neglect the temperature-dependence of $\Delta H_d^\circ$ and $\Delta S_d^\circ$, which is set by the change in heat capacity ($\Delta C_P$) between states. $\Delta C_P$ has previously been measured for dehybridization of DNA oligonucleotides to give a length-scaling of ~0.18 kJ mol$^{-1}$ K$^{-1}$ bp$^{-1}$ but it also depends on sequence, ionic strength, and temperature.(18-20) It is also unclear whether coaxial stacking and changes in gap or overhang conformation following $A_n$ unbinding will significantly change $\Delta C_P$.

If the template is an overhang (eq. S3c) rather than gap, then primer unbinding can be treated identically to dissociation of $A_n$.

$$\theta_{\text{Temp}} = 1 + \frac{K_{\text{d,Temp}} - \sqrt{K_{\text{d,Temp}}^2 + 2c_2 K_{\text{d,Temp}}}}{c_2} \tag{S9}$$

where, $c_2 = [\text{X}] + [\text{Pri}] + [\text{Tmp}]$

$$T_{m,\text{Temp}} = \frac{\Delta H_{d,\text{Temp}}^\circ}{\Delta S_{d,\text{Temp}}^\circ - R\ln\left(c_2/4\right)} \tag{S10}$$

Primer and helper dissociation from a gap template involves a network of bimolecular equilibria with nearly identical unbinding constants represented by $K_{\text{d,Temp}}$. Unbinding of primer and helper strands may be cooperative or non-cooperative. For completely non-cooperative unbinding, where unbinding of the primer and helper from the template strand are independent of one another, the total fraction of bound primer and helper ($\theta_{\text{Temp,NC}}$) may be approximated as the product of each binding fraction.

$$\theta_{\text{Temp,NC}} = \theta_{\text{Temp}}^2 \tag{S11}$$

In the opposite limit of high cooperativity, the primer and helper may unbind from the template strand in an all-or-nothing fashion. In our case, [Tmp]=[Pri]=[Hlp], leading to:(21)

$$K_{\mathrm{d,TempC}} = \frac{\left(1-\theta_{\mathrm{Temp,C}}\right)^3 c_2^2}{9\theta_{\mathrm{Temp,C}}} \quad \text{(S12)}$$

$$\text{where, } \theta_{\mathrm{Temp,C}} = \frac{3[\mathrm{X}]}{c_2} \text{ and } c_2 = 3[\mathrm{X}]+[\mathrm{Pri}]+[\mathrm{Hlp}]+[\mathrm{Tmp}]$$

Arranging in terms of $\theta_{\mathrm{Temp,C}}$ gives a cubic polynomial:

$$c_2^2\theta_{\mathrm{Temp,C}}^3 - 3c_2^2\theta_{\mathrm{Temp,C}}^2 + \left[3c_2^2 + 9K_{\mathrm{d,TempC}}\right]\theta_{\mathrm{Temp,C}} - c_2^2 = 0 \quad \text{(S13)}$$

$\theta_{\mathrm{Temp,C}}$ may be solved for by reducing eq. S13 to a depressed cubic polynomial and applying Cardano's method and Vieta's substitution.(22)

$$\theta_{\mathrm{Temp,C}} = \sqrt[3]{W} - \frac{p}{3\sqrt[3]{W}} \quad \text{(S14a)}$$

$$W = -\frac{q}{2} + \sqrt{\frac{q^2}{4} + \frac{p^3}{27}} \quad \text{(S14b)}$$

$$\text{where, } p = \frac{9K_{\mathrm{d,TempC}}}{c_2^2} \text{ and } q = -\frac{9K_{\mathrm{d,TempC}}}{c_2^2}$$

As for the bimolecular equilibria, the melting temperature ($T_{m,\mathrm{Temp,C}}$) is defined as the temperature where $\theta_2$ is 0.5.

$$T_{m,\mathrm{TempC}} = \frac{\Delta H^\circ_{d,\mathrm{TempC}}}{\Delta S^\circ_{d,\mathrm{TempC}} - R\ln\left(c_2^2/36\right)} \quad \text{(S15)}$$

The non-cooperative approximation (eq. S11) is likely suitable for the sequences studied in this work because the binding regions for primer and helper strands are separated by at least a two nucleotide gap. We also find that the primer and helper melting profile is identical for gap sizes of one to four nucleotides (Fig. S5), supporting that coaxial stacking interactions between primer and helper strands are negligible.

##### S3.2 Sequential dissociation model

In reality, the $K_{\mathrm{d}}$ and $K_{\mathrm{d,Temp}}$ equilibria are coupled to a degree depending on the sequence of each oligonucleotide. Coaxial stacking of the primer and helper with $A_n$ stabilizes both X:$A_n$ as well as binding of the primer and helper to the template strand, and $A_n$ unbinding or primer and helper unbinding will reduce the binding stability of the other. Therefore, we can treat the processes in eqs. S3abc and eqs. S3a,d as sequential equilibria.

For sequential bimolecular equilibria, the concentration of each species follows a 4th order polynomial, and we find that the solution to this polynomial is not stable over the full temperature range that covers both melting transitions. Therefore we approximate $K_{\mathrm{d,Temp}}$ as a unimolecular equilibrium between X and a "dissociated" template (dX).

$$\mathrm{X{:}A_n} \underset{}{\overset{K_\mathrm{d}}{\rightleftharpoons}} \mathrm{X + A_n} \underset{}{\overset{K_\mathrm{d,Temp}}{\rightleftharpoons}} \mathrm{dX + A_n} \tag{S16}$$

where each equilibrium constant may be expressed using a van't Hoff form as in eq. S7.

$$K_\mathrm{d}(T) = \frac{[\mathrm{X}][\mathrm{A_n}]}{[\mathrm{X{:}A_n}]} = \exp\left[-\frac{\Delta H_d^\circ}{RT} + \frac{\Delta S_d^\circ}{R}\right] \tag{S17a}$$

$$K_\mathrm{d,Temp}(T) = \frac{[\mathrm{dX}]}{[\mathrm{X}]} = \exp\left[-\frac{\Delta H_{d,Temp}^\circ}{RT} + \frac{\Delta S_{d,Temp}^\circ}{R}\right] \tag{S17b}$$

The concentrations of each species can be determined in terms of the equilibrium constants as well as the total concentration of template strand ($c_\mathrm{Temp}$) and $A_n$ ($c_{\mathrm{A_n}}$), which are known values and independently controlled in experiments.

$$c_{\mathrm{A_n}} = [\mathrm{X{:}A_n}] + [\mathrm{A}_n] \tag{S18a}$$

$$c_\mathrm{Temp} = [\mathrm{X{:}A_n}] + [\mathrm{X}] + [\mathrm{dX}] \tag{S18b}$$

By re-arranging eq. S18b in terms of [dX] and using eqs. S17a and S17b, we can express [dX]:

$$[\mathrm{dX}] = c_\mathrm{Temp} - \frac{[\mathrm{dX}]}{K_\mathrm{d,Temp}} - \frac{[\mathrm{dX}][\mathrm{A_n}]}{K_\mathrm{d}K_\mathrm{d,Temp}} \tag{S19}$$

$[A_n]$ is determined from equating $[X{:}A_n]$ from eqs. S18a and S18b and using eq. S17b.

$$[\mathrm{A}_n] = c_{\mathrm{A}_n} - c_\mathrm{Temp} + \frac{[\mathrm{dX}]}{K_\mathrm{d,Temp}} + [\mathrm{dX}] \tag{S20}$$

Substituting eq. S20 into eq. S19 and simplifying leads to a quadratic polynomial in terms of [dX].

$$\left(\frac{1+K_{\mathrm{d,Temp}}}{K_{\mathrm{d}}K_{\mathrm{d,Temp}}^{2}}\right)[\mathrm{dX}]^{2}+\left(1+\frac{K_{\mathrm{d}}+c_{\mathrm{A}_{n}}-c_{\mathrm{Temp}}}{K_{\mathrm{d}}K_{\mathrm{d,Temp}}}\right)[\mathrm{dX}]-c_{\mathrm{Temp}}=0 \tag{S21a}$$

where,

$$[\mathrm{dX}]=\frac{-K_{\mathrm{d,Temp}}\left(K_{\mathrm{d,Temp}}K_{\mathrm{d}}+K_{\mathrm{d}}-c_{\mathrm{Temp}}+c_{\mathrm{A}_{n}}\right)+K_{\mathrm{d,Temp}}\sqrt{\left(K_{\mathrm{d}}K_{\mathrm{d,Temp}}+K_{\mathrm{d}}+c_{\mathrm{A}_{n}}-c_{\mathrm{Temp}}\right)^{2}+4c_{\mathrm{Temp}}\left(1+K_{\mathrm{d,Temp}}\right)}}{2\left(1+K_{\mathrm{d,Temp}}\right)} \tag{S21b}$$

The fractions of intact X ($\theta_{\mathrm{Temp}}$) and X:A$_{\mathrm{n}}$ ($\theta_{\mathrm{A_n}}$) are computed separately for each equilibrium and under the condition $c_{\mathrm{A_n}}=c_{\mathrm{Temp}}$, which is used for all temperature-dependent FTIR and NMR measurements.

$$\theta_{\mathrm{Temp}}=\frac{[\mathrm{X}]}{c_{\mathrm{Temp}}}=1+\frac{K_{\mathrm{d,Temp}}\left(K_{\mathrm{d,Temp}}K_{\mathrm{d}}+K_{\mathrm{d}}\right)-K_{\mathrm{d,Temp}}\sqrt{\left(K_{\mathrm{d}}K_{\mathrm{d,Temp}}+K_{\mathrm{d}}\right)^{2}+4c_{\mathrm{Temp}}\left(1+K_{\mathrm{d,Temp}}\right)}}{2\left(1+K_{\mathrm{d,Temp}}\right)} \tag{S22a}$$

$$\theta_{\mathrm{A_n}}=\frac{[\mathrm{X{:}A_n}]}{c_{\mathrm{A_n}}}=\frac{c_{\mathrm{Temp}}^{2}\left(1-\theta_{\mathrm{Temp}}\right)^{2}}{c_{\mathrm{A_n}}K_{\mathrm{d}}K_{\mathrm{d,Temp}}\left(K_{\mathrm{d,Temp}}+c_{\mathrm{Temp}}\left(1-\theta_{\mathrm{Temp}}\right)\right)} \tag{S22b}$$

The melting temperature for AA dissociation, $T_m$, is defined as the temperature where half of the A$_{\mathrm{n}}$ is bound to the template.

$$\Delta G_{d}^{\circ}=-RT_{m}\ln\left(c_{\mathrm{A}_{n}}/4\right) \tag{S23}$$

##### S3.3 Fitting of $^{1}$H NMR temperature series data

The temperature-dependence trends in chemical shift of A3, A4, C1, and C2 resonances (Fig. 2) were globally fit to the sequential three-state model (eq. S16). A3 and A4 peak chemical shifts were used for the first component ($\theta_{\mathrm{A_n}}$) and those of C1 and C2 for the second component ($\theta_{\mathrm{Temp}}$). Linear temperature-dependent changes in chemical shift also contribute to each peak and are corrected for using baseline fits. The high-temperature baselines of A3 and A4 are constrained to those measured for free AA (Fig. S9a). The extracted melting curves and thermodynamic parameters for the AA-gap complex and template dissociation quantitatively agree with those determined from global fitting of FTIR and 2D IR temperature series (Fig. 4) as well as ITC measurements (Figs. S26-S27).

##### S3.4 Global fitting of FTIR and 2D IR temperature series

Global fitting is applied to describe multiple melting components present in the FTIR and 2D IR temperature series (Fig. 2). Global fitting describes the temperature-dependence at each frequency with a set of shared fitting parameters, which in this case are $\Delta H^{\circ}_{d,\mathrm{i}}$ and $\Delta S^{\circ}_{d,\mathrm{i}}$ for $i$ components. The frequency- and temperature-dependent absorption data ($A(\omega,\mathrm{T})$) is described by the sum of amplitude-weighted melting components $\theta_{\mathrm{A_n}}$ and $\theta_{\mathrm{Temp}}$ plus an offset $c$.

$$A(\omega,T)=C(T)E(\omega)^{t}=\begin{bmatrix}\theta_{\mathrm{A}_n}(T)\\ \theta_{\mathrm{Temp}}(T)\\ 1\end{bmatrix}\begin{bmatrix}a(\omega)\\ b(\omega)\\ c(\omega)\end{bmatrix}^{t} \tag{S24}$$

Equation S24 is expressed in a general form where $\theta_{\mathrm{A_n}}$ and $\theta_{\mathrm{Temp}}$ can be independent (eqs. S6, S11, and S14) or dependent through a sequential model (eqs. S22ab). The component spectra $a$ and $b$ correspond to difference FTIR and 2D IR spectra associated with $\mathrm{A_n}$ unbinding and primer and helper unbinding, respectively (Figs. 4, S19-S20). The value of $\theta_{\mathrm{A_n}}$ at 1 °C determined from FTIR titrations (Figs. 1 & S1) was included in the fitting to constrain $\theta_{\mathrm{A_n}}$. This constraint is unnecessary for many sequences, but was particularly important to extract accurate melting curves for UUg:AA and TTo:AA (Fig. S20).

As is often done for global fitting of time-dependent spectra,(23,24) gradient-based alternating least squares is applied so that the objective function ($F$) to be optimized during fitting only contains the parameters of C.

$$F=\left\|\left(I-C(T,n)C(T,n)^{+}\right)\Psi\right\|_{2} \tag{S25}$$

$$\text{where } E(\omega)^{T}=C^{+}(T)\Psi$$

$C^{+}$ indicates the Moore-Penrose pseudoinverse of C. 2D IR spectra are reshaped from an m x n matrix, where m and n are the number of $\omega_1$ and $\omega_3$ points, respectively, to a vector of length m*n for fitting as described in previous application of global lifetime fitting to 2D spectra.(25) After fitting, the extracted component spectra are reshaped into an m x n matrix.

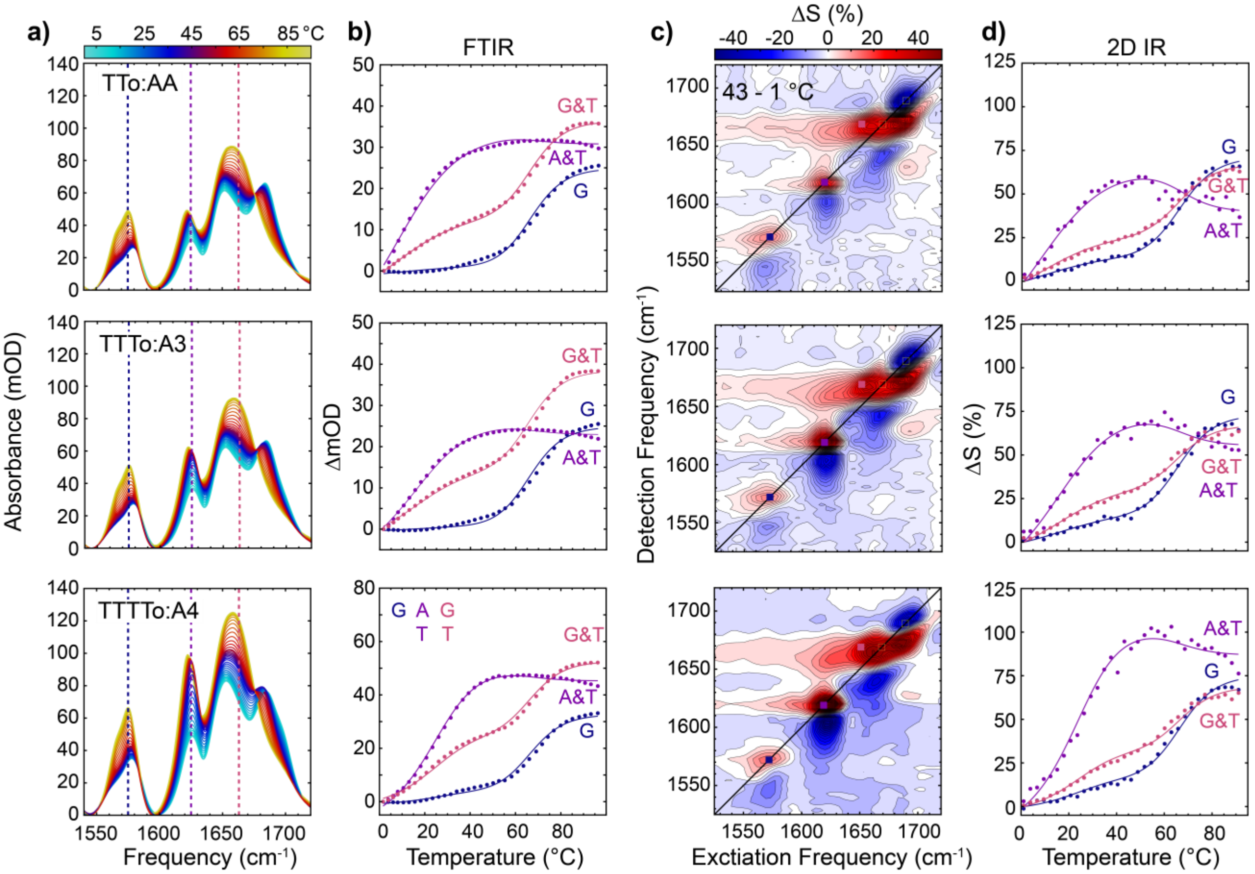


**Figure S17. Global fitting of temperature-dependent IR spectra for overhang sequences. (a)** FTIR spectra of TTo:AA, TTTo:A3, and TTTTo:A4 from 1 to 96 °C in ~2.6 °C steps. **(b)** Temperature-dependent change in absorption relative to 1 °C for select frequencies. Frequencies are indicated in **(a)** with vertical dashed lines. **(c)** 2D IR difference spectra between 43 and 1 °C. Spectra are plotted in units of percentage change relative to maximum value of the 1 °C spectrum (ΔS). 25 contours with uniform 2% spacing are plotted for each sequence. **(d)** Temperature-dependent spectral change relative to the spectrum at 1 °C at select frequencies indicated by colored squares on the 2D IR spectra in **(c)**. Solid lines in **(b)** and **(d)** correspond to global fits to a three-state sequential model (eq. S22a,b).

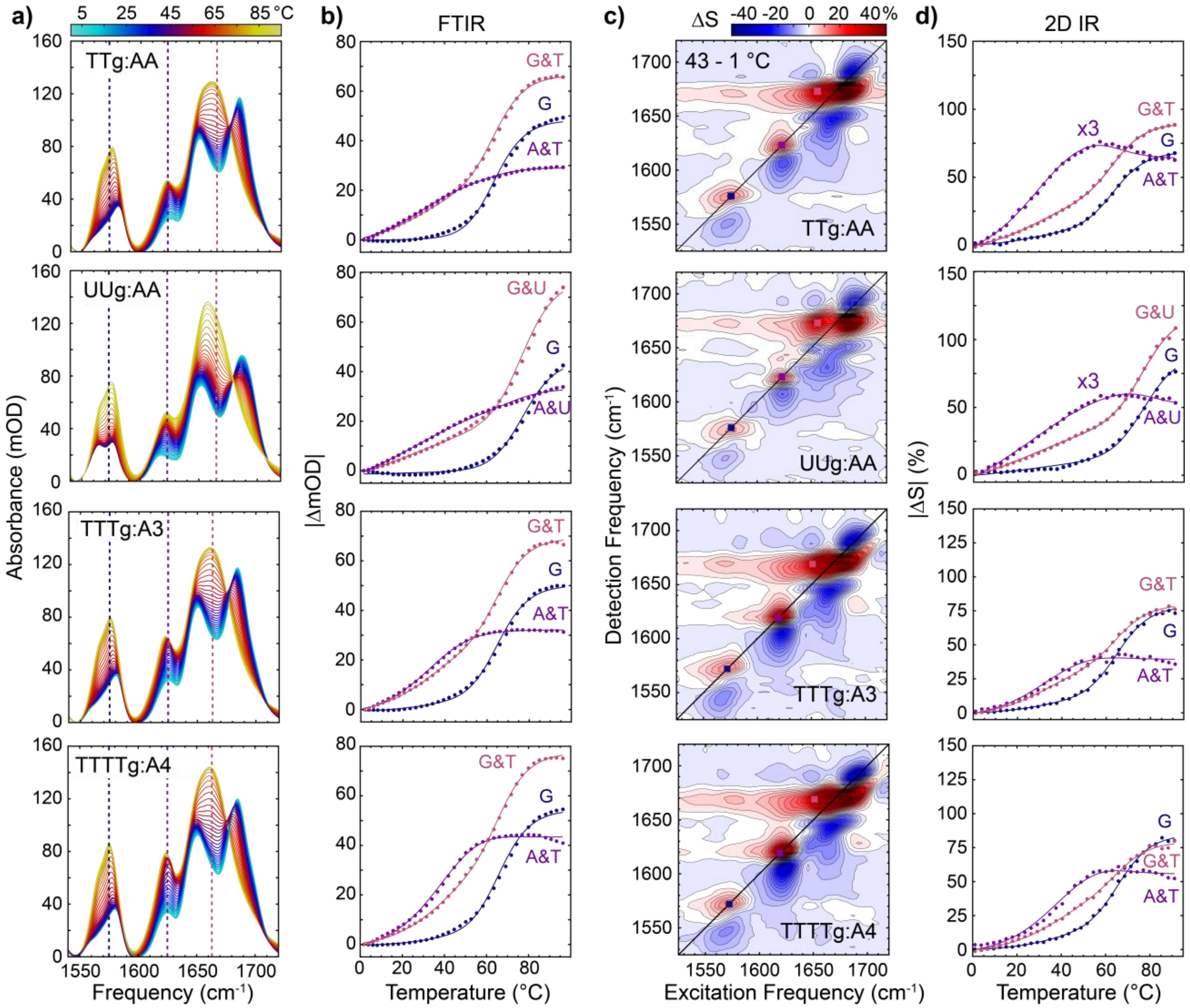


**Figure S18. Global fitting of temperature-dependent IR spectra for gap sequences. (a)** FTIR spectra of TTg:AA, UUg:AA, TTTg:A3 and TTTTg:A4 from 1 to 96 °C in ~2.6 °C steps. **(b)** Temperature-dependent change in absorption relative to 1 °C for select frequencies. Frequencies are indicated in **(a)** with vertical dashed lines. **(c)** 2D IR difference spectra between 43 and 1 °C. Spectra are plotted in units of percentage change relative to maximum value of the 1 °C spectrum (ΔS). 25 contours with uniform 2% spacing are plotted for each sequence. **(d)** Temperature-dependent spectral change relative to the spectrum at 1 °C at select frequencies indicated by colored squares on the 2D IR spectra in **(c)**. Solid lines in **(b)** and **(d)** correspond to global fits to a three-state sequential model (eq. S22a,b).

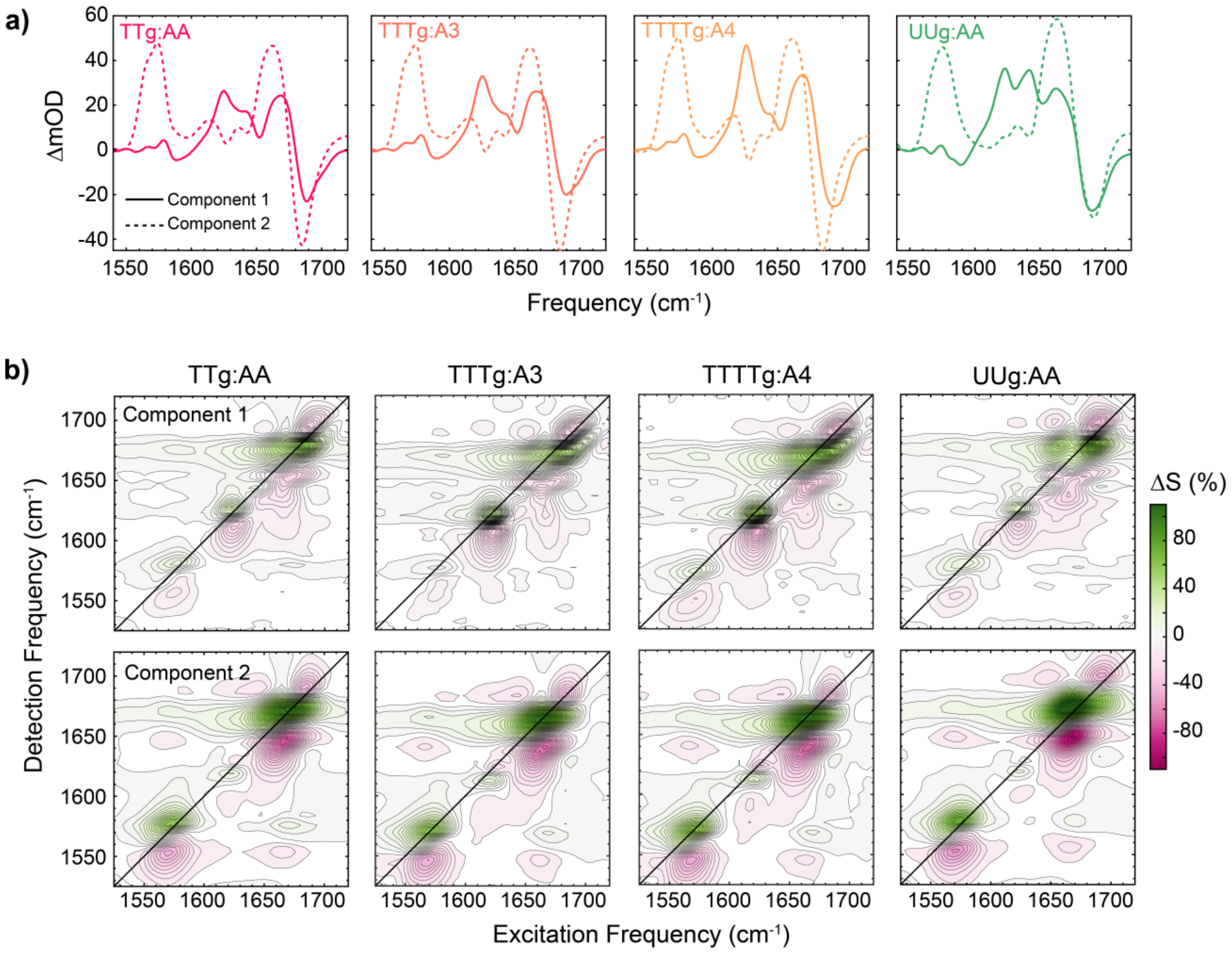


**Figure S19. FTIR and 2D IR component spectra from thermodynamic global fitting of $A_n$-gap complex temperature series. (a)** (solid lines) first and (dashed lines) second FTIR spectral components extracted from global fitting of $A_n$-gap complex FTIR and 2D IR temperature series (Figs. 2 & S17) to a three-state sequential model (eq. S22a,b). **(b)** (top) first and (bottom) second 2D IR spectral components plotted in terms of signal percent change (ΔS). 25 contours with uniform 2% spacing are plotted for each sequence.

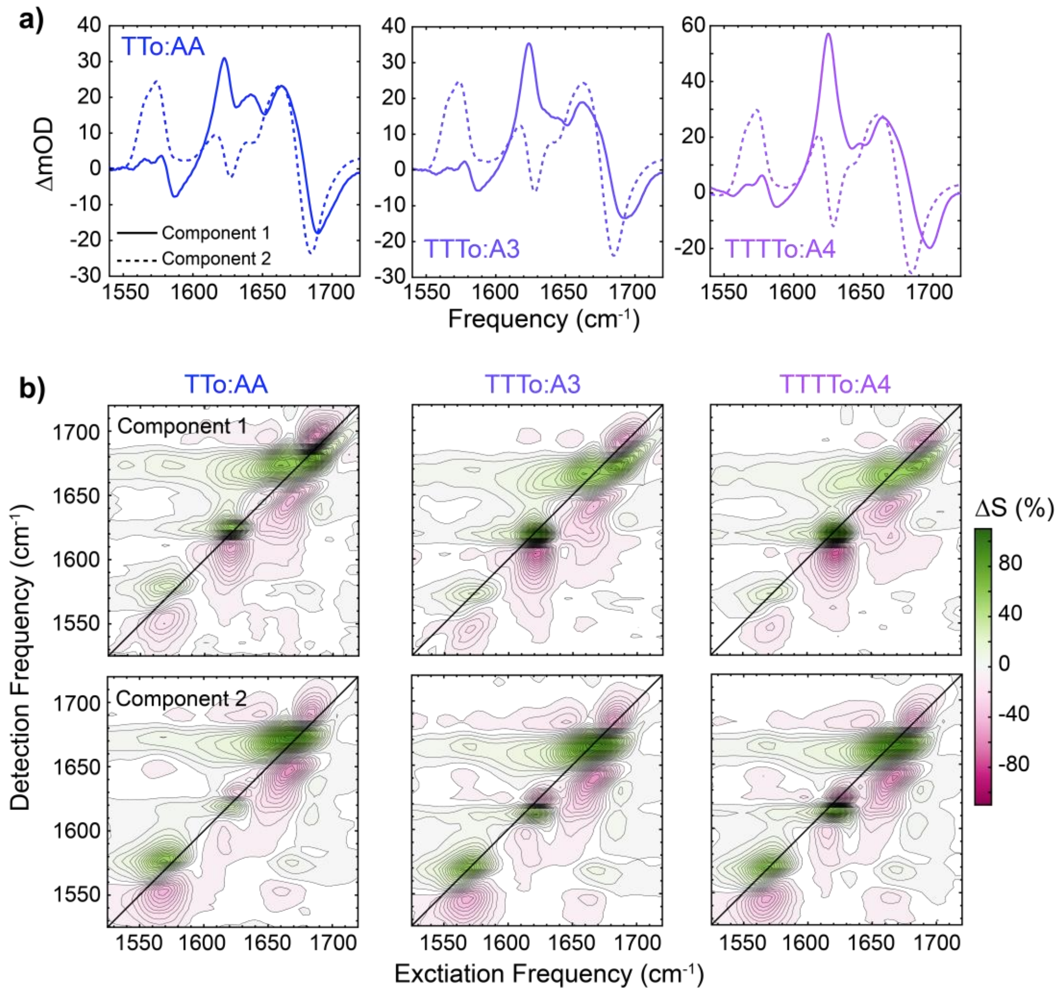


**Figure S20. FTIR and 2D IR component spectra from thermodynamic global fitting of $A_n$-overhang complex temperature series. (a)** (solid lines) first and (dashed lines) second FTIR spectral components extracted from global fitting of $A_n$-overhang complex FTIR and 2D IR temperature series (Fig. S18) to a three-state sequential model (eq. S22a,b). **(b)** (top) first and (bottom) second 2D IR spectral components plotted in terms of signal percent change (ΔS). 25 contours with uniform 2% spacing are plotted for each sequence.

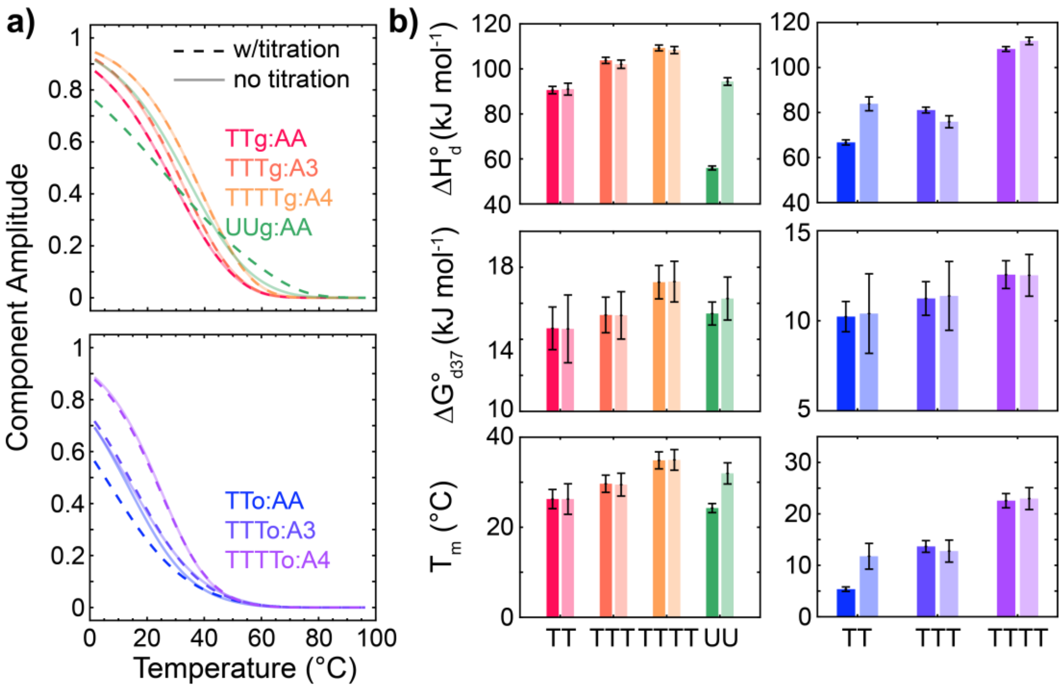


**Figure S21. Impact of low-temperature titration constraint on thermodynamic global fitting.** **(a)** First component melting curves from global fitting of FTIR and 2D IR temperature series to a sequential model with (dark) and without (light) a constraint on the amplitude at 1 °C from FTIR-monitored titrations (Figs. 1 and S1). **(b)** Enthalpy ($\Delta H_d^\circ$), free energy at 37 °C ($\Delta G_{d37}^\circ$), and melting temperature ($T_m$) from fits with (dark) and without (light) a constraint on the amplitude at 1 °C from FTIR-monitored titrations. Error bars are determined from 95% confidence intervals in fit parameters.

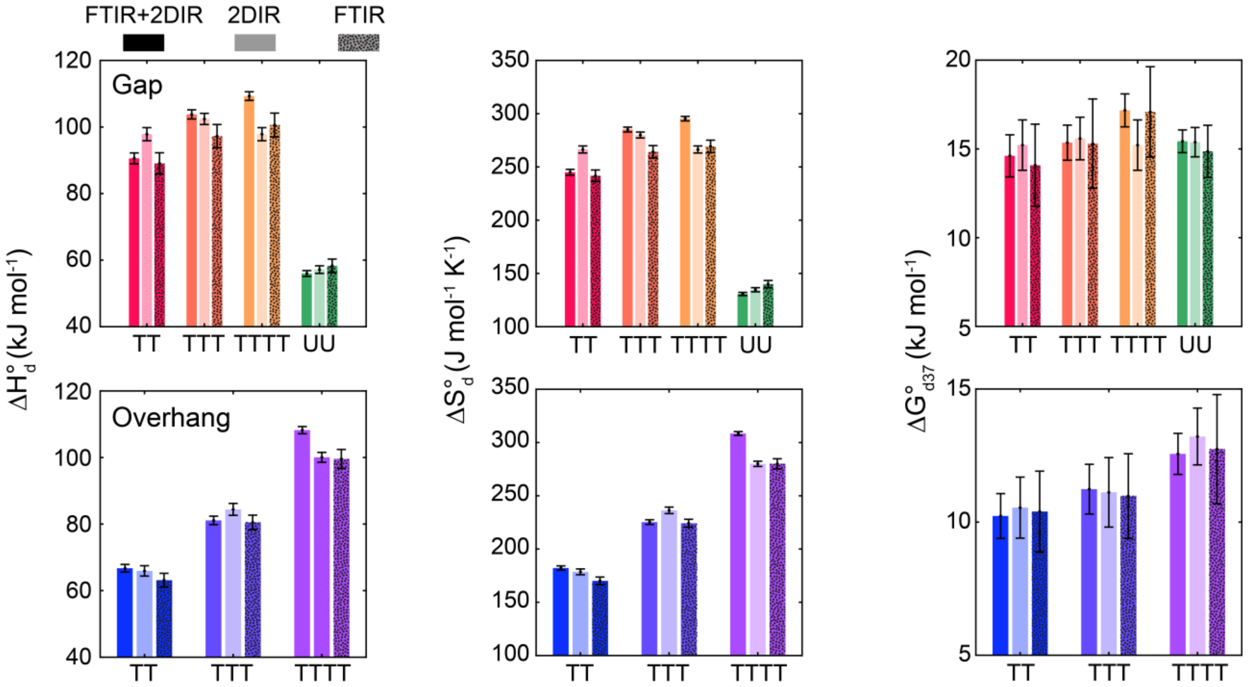


**Figure S22. Comparison of $A_n$ dissociation thermodynamics extracted from global fitting with FTIR or 2D IR data or both together.** $\Delta H^{\circ}_{d}$ (left), $\Delta S^{\circ}_{d}$ (center), and $\Delta G^{\circ}_{d37}$ (right) obtained from global fitting to a three-state sequential model (Section S3.2). Error bars are determined from 95% confidence intervals in fit parameters.

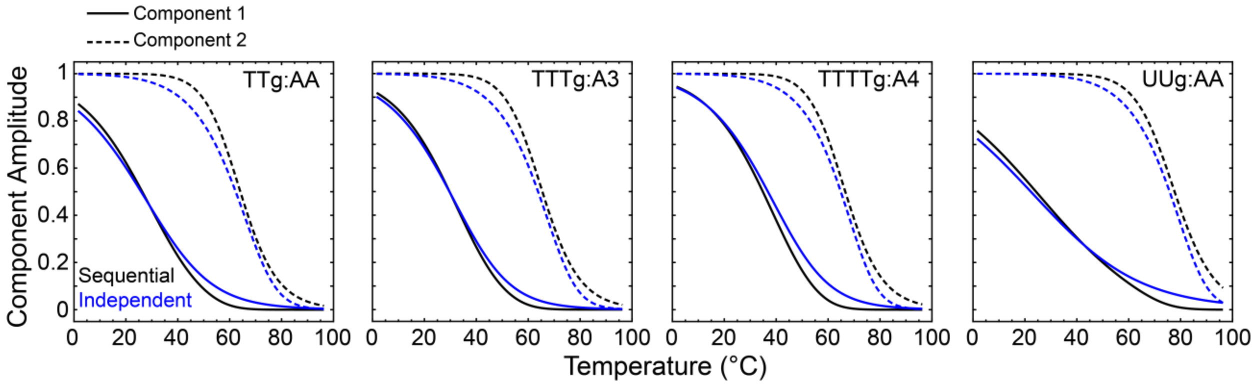


**Figure S23. Comparison of fitting with a sequential model and independent component model.** Melting curves extracted from global fits of FTIR and 2D IR temperature series using a sequential model (black lines, Section S3.2) and an independent model (blue lines, Section S3.1). Component 2 (dashed lines) is treated as a series of non-cooperative bimolecular equilibria (eq. S11) in the independent model and as a unimolecular equilibrium in the sequential model, which leads to different melting curve shapes.

### S4. Thermodynamics of free strand binding

#### S4.1 FTIR temperature series of GA2G, GA3G, and GA4G

For comparison with the DNA gaps and overhangs, we measured hybridization thermodynamics of GA2G, GA3G, and GA4G (Fig. 7). FTIR temperature series reveal a single melting transition and melting curves are extracted as the 2$^{nd}$ SVD component using the frequency window from 1550 to 1720 cm$^{-1}$. The 2$^{nd}$ SVD components can be fit ($V_{Fit}^{(2)}$) to a two-state thermodynamic model identical to eq. S6 but with the inclusion of upper ($L_D$) and lower ($L_S$) baselines to extract the bound duplex fraction ($\theta_D$).(6,7)

$$V_{Fit}^{(2)} = \theta_D \left( L_D - L_S \right) + L_S \quad \text{(S26)}$$

$$\text{where, } L_D = m_D T + b_D \text{ and } L_S = m_S T + b_S$$

Even at an oligonucleotide concentration of 10 mM, the binding stability for GA2G and GA3G is too weak to fit an upper baseline in the measured temperature range. Therefore, the upper baseline slope ($m_D$) fit for GA4G is fixed for GA3G and GA2G. Using the mean value of $\Delta H_d^{\circ}$ and $\Delta S_d^{\circ}$ from FTIR and ITC (Table S2), melting curves at an effective concentration of 2 mM were calculated using eq. S6 to compare with the $A_n$ dissociation curves (Fig. 7).

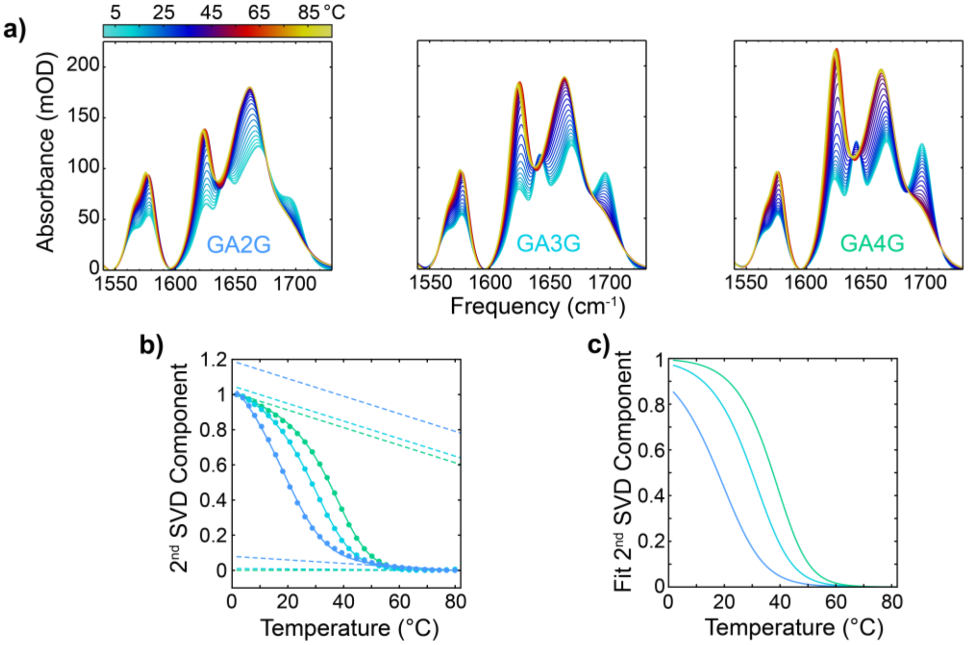

**Figure S24. Temperature-dependent FTIR measurements of free single-strands.** **(a)** FTIR temperature series of GA2G, GA3G, and GA4G sequences shown in Fig. 7**.** Measurements were performed with an equimolar strand ratio and a total oligonucleotide concentration of 10 mM in pH* 6.8 400 mM SPB. **(c)** $2^{nd}$ SVD components from FTIR temperature series fit to a two-state model (solid lines) containing upper and lower baselines (dashed lines). The upper baseline of all sequences was fixed to that determined from a fit the GA4G data. **(d)** Duplex fraction melting curves from baseline-removal of fit $2^{nd}$ SVD components.

#### S4.2 ITC of free strand binding and $A_n$ binding to gaps

ITC measurements were performed to directly measure $\Delta H_d^\circ$ for the hybridization processes. Following each injection, the power required to re-equilibrate the sample and reference cells was monitored as a function of time. The duration of the 2 μL injections were varied depending on the sample in order to avoid saturation. The power profile of each injection was integrated over time to obtain the change in heat from the injection (ΔQ). Curves of ΔQ vs molar ratio of titrant and cell oligonucleotide were fit to a single site binding model(26) with free parameters $\Delta H_d^\circ$, association constant ($K_a$), and reaction stoichiometry ($n$).

$$\Delta Q_i = Q_i - Q_{i-1} + \frac{V_{inj}}{V_o}\left[\frac{Q_i + Q_{i-1}}{2}\right] \tag{S27}$$

$$Q_i = \frac{nM_{c,i}V_o\Delta H^\circ}{2}\left[1+\frac{T_{c,i}}{nM_{c,i}}-\frac{1}{nK_aM_{c,i}}-\sqrt{\left(1+\frac{T_{c,i}}{nM_{c,i}}+\frac{1}{nK_aM_{c,i}}\right)^2-\frac{4T_{c,i}}{nM_{c,i}}}\right] \tag{S28}$$

$Q_i$ is the total heat exchanged after injection $i$ of titrant. $M_{c,i}$ and $T_{c,i}$ are the concentration of cell oligonucleotide and titrant oligonucleotide, respectively, in the cell after injection $i$. $V_o$ is the initial volume of sample in the cell, which was 200 μL for all measurements, and $V_{inj}$ is the volume of titrant used in each injection (2 μL). The sample conditions and injection settings for each measurement are listed in Table S2.

**Table S2.** Experimental conditions used for ITC measurements. All samples were prepared in 400 mM pH 6.8 SPB.

| Sequence | T (°C) | Syringe Strand | [Syringe] (μM) | Cell Strand | [Cell] (μM) | Injection Duration (s) | Injection Interval (s) |
|---|---|---|---|---|---|---|---|
| | | | | **Gap** | | | |
| TTg:AA | 8 | AA | 12000 | TTg | 700 | 8 | 180 |
| TTTg:A3 | 8 | AAA | 2500 | TTTg | 150 | 8 | 180 |
| TTTTg:A4 | 8 | AAAA | 1000 | TTTTg | 60 | 4 | 180 |
| UUg:AA | 8 | AA | 7000 | UUg | 500 | 8 | 180 |
| | | | | **Free strands** | | | |
| GA2G | 5 | CTTC | 40000 | GAAG | 3000 | 210 | 390 |
| GA3G | 5 | CTTTC | 5000 | GAAAG | 500 | 30 | 210 |
| GA4G | 5 | CTTTTC | 700 | GAAAAG | 50 | 4 | 180 |

**Table S3.** Thermodynamic parameters for $A_n$ dissociation from gaps and overhangs as well as dissociation between $GA_nG$ and its complement. All samples were prepared in 400 mM pH 6.8 SPB. Error bars correspond to 95% confidence from fitting data to thermodynamic models.

| | | FTIR + 2D IR | | | | | ITC | |
|---|---|---|---|---|---|---|---|---|
| Sequence | | $\Delta H^\circ_d$ (kJ mol$^{-1}$) | $\Delta S^\circ_d$ (J mol$^{-1}$ K$^{-1}$) | $\Delta G^\circ_{d8}$ (kJ mol$^{-1}$) | $\Delta G^\circ_{d,37}$ (kJ mol$^{-1}$) | $T_m$ (°C) | $\Delta H^\circ_d$ (kJ mol$^{-1}$) | $\Delta G^\circ_{d8}$ (kJ mol$^{-1}$) |
| $A_n$-Gap[a] | TTg:AA | 91 ± 2 | 245 ± 6 | 21.7 ± 1.1 | 14.6 ± 1.2 | 26 ± 2 | 83 ± 5 | 23.1 ± 1.3 |
| | TTg:AA (NMR) | 79 ± 5 | 203 ± 20 | 21.5 ± 4.5 | 15.1 ± 5.5 | 28 ± 3 | - | - |
| | TTTg:A3 | 104 ± 2 | 285 ± 5 | 23.6 ± 0.9 | 15.4 ± 1.0 | 30 ± 2 | 94 ± 2 | 25.8 ± 0.7 |
| | TTTTg:A4 | 109 ± 2 | 295 ± 4 | 26.2 ± 0.9 | 17.2 ± 0.9 | 35 ± 2 | 109 ± 4 | 29.4 ± 0.9 |
| | UUg:AA | 56 ± 1 | 131 ± 3 | 19.2 ± 0.6 | 15.4 ± 0.6 | 24 ± 1 | 51 ± 1 | 23.5 ± 0.4 |
| $A_n$-Toe[a] | TTo:AA | 67 ± 1 | 182 ± 4 | 16.1 ± 0.8 | 10.2 ± 0.9 | 5 ± 1 | - | - |
| | TTTo:A3 | 81 ± 2 | 225 ± 4 | 18.4 ± 0.9 | 11.2 ± 0.9 | 14 ± 1 | - | - |
| | TTTTo:A4 | 108 ± 2 | 308 ± 4 | 22.4 ± 0.7 | 12.5 ± 0.8 | 23 ± 1 | - | - |
| Free[b,c] strands | GA2G | 124 ± 5 | 377 ± 37 | 18.1 ± 5.7 | 7.2 ± 5.8 | 18 ± 2 | 102 ± 4 | 19.0 ± 0.3 |
| | GA3G | 155 ± 3 | 461 ± 21 | 25.2 ± 3.4 | 11.8 ± 3.5 | 30 ± 1 | 127 ± 5 | 23.4 ± 0.4 |
| | GA4G | 181 ± 3 | 532 ± 16 | 31.1 ± 2.5 | 15.7 ± 2.6 | 37 ± 1 | 152 ± 10 | 28.0 ± 0.6 |

[a]Determined from global fitting to three-state sequential model (eq. S16). [b]Prepared at 10 mM total oligonucleotide concentration. [c]Determined with two-state fitting of 2$^{nd}$ SVD component from FTIR temperature series (Fig. S24).

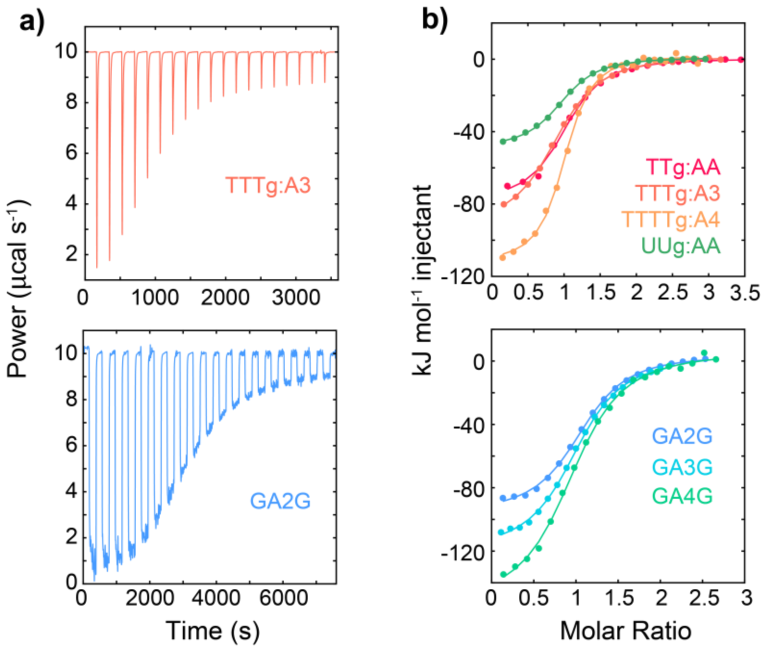


**Figure S25. Isothermal titration calorimetry of gap templates and free single-strand DNA. (a)** Raw ITC thermograms for TTTg:AA and GA2G showing the measured power vs. time. Slower injection rates were used to avoid signal saturation is more concentrated samples (Table S2). **(b)** Integrated ITC thermograms for AA binding onto gaps (top) and binding between free oligonucleotides (bottom). Data are fit to a two-state model (solid lines, eq. S26). Sample conditions for each measurement are listed in Table S2.

##### S4.3 Length-dependent hybridization thermodynamics and kinetics

Temperature-dependent T-jump IR measurements of A3 and A4 dissociation were performed to assess how length-scaling of hybridization between a short segment and gap or overhangs differs from free oligonucleotides. Both $k_d$ and $k_a$ decrease with increasing strand length for gaps and overhangs in the measured temperature range. It is well established that $k_d$ decreases and $\Delta H_d^{\ddagger}$, $\Delta S_d^{\ddagger}$, and $\Delta G_d^{\ddagger}$ increase with length for dissociation between free strands.(27,28) For gaps, $\Delta G_{d37}^{\ddagger}$ increases more sharply than $\Delta G_{d37}^{\circ}$ ($\Delta\Delta G_{d37}^{\ddagger}$, 3.5 vs. 1.0 kJ mol$^{-1}$ bp$^{-1}$) and $\Delta G_{a37}^{\ddagger}$ increases by 2.3 kJ mol$^{-1}$ bp$^{-1}$ (Fig. S28). $\Delta G_a^{\ddagger}$ is typically reported to remain constant or slightly increase ($\Delta\Delta G_{a37}^{\ddagger}$ < 1 kJ mol$^{-1}$ bp$^{-1}$) with DNA strand length,(28,29) although a recent study found a reduction in $\Delta G_a^{\ddagger}$ with increasing RNA strand length.(30) Therefore, our trends for gap DNA indicate that a length-dependence in the free energy of the transition-state or template state contribute to $\Delta G_{d37}^{\ddagger}$ in addition to an increase in thermodynamic stability of the hybridized complex state. Similar, yet more modest trends are observed for overhangs with

$\Delta\Delta G^{\ddagger}_{d37}$ = 2.8 kJ mol$^{-1}$ bp$^{-1}$ and $\Delta\Delta G^{\ddagger}_{a37}$ = 1.8 kJ mol$^{-1}$ bp$^{-1}$. Together, these results suggest that distinct thermodynamic effects from gaps and overhangs stabilize AA binding beyond what may be predicted from canonical single-strand hybridization yet also reduce the energetic benefit of extending the binding segment from 2 to 4 nucleotides.

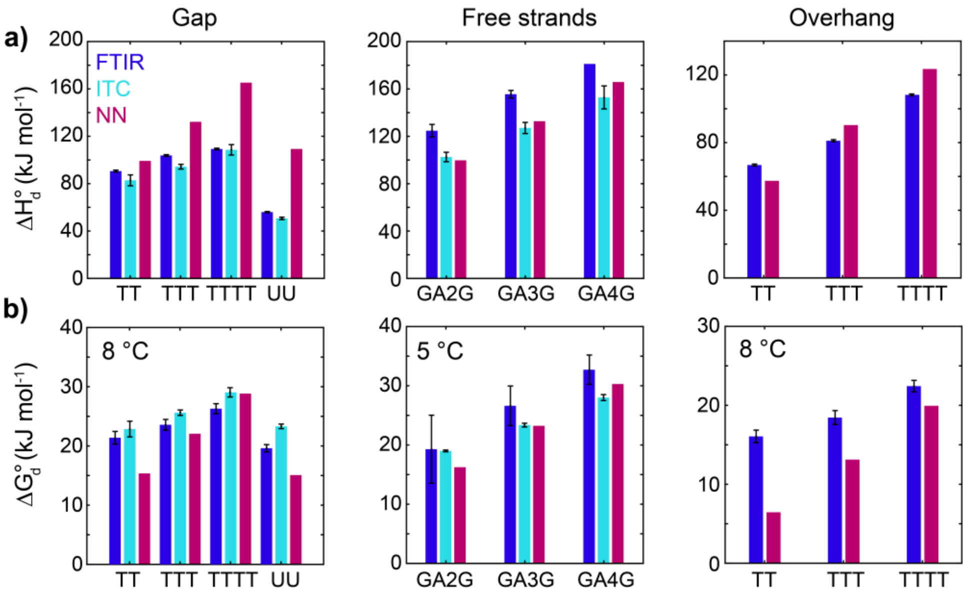


**Figure S26. Dissociation thermodynamics for free single-strands and gaps from isothermal titration calorimetry with comparison to FTIR and nearest-neighbor (NN) model. (a)** Unbinding enthalpy ($\Delta H^{\circ}_{d}$) and **(b)** free energy ($\Delta G^{\circ}_{d}$) at listed temperatures determined from FTIR temperature series (blue) and ITC (cyan) and calculated using DNA and RNA NN models corrected to [Na$^{+}$] = 600 mM.(31-34) For gaps and free single-strands, NN calculations are performed on GA$_{n}$G segments whereas GA$_{n}$ segments are used for comparison with overhangs. Initiation parameters are applied in both cases.

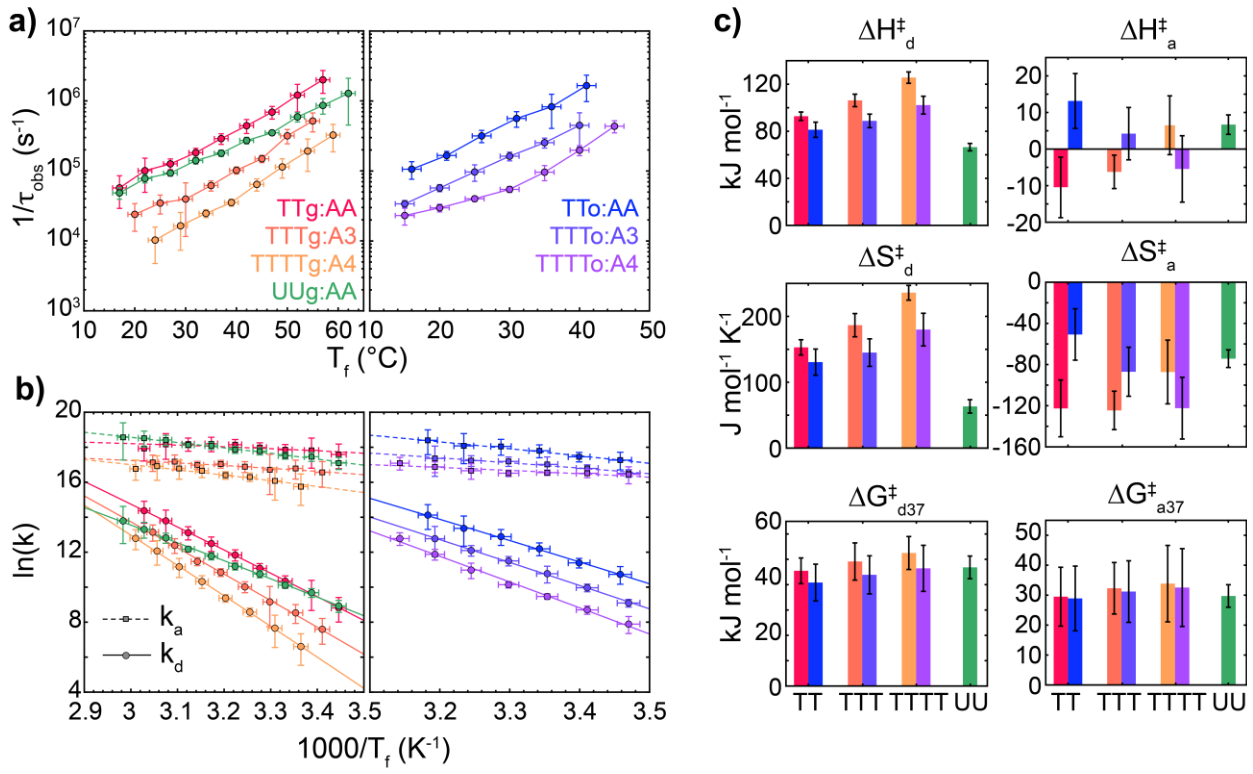


**Figure S27. Length-scaling of kinetics for $A_n$ association and dissociation from gaps and overhangs (a)** Observed relaxation rate ($k_{obs}$) for AA dissociation as a function of $T_f$ for all sequences. Rates correspond to the amplitude-weighted mean across the adenine ring mode response (1585 to 1610 cm$^{-1}$) in the rate-domain spectra determined from MEM-iLT.(16,17) Vertical error bars indicate standard deviation over the rate spectra and horizontal error bars correspond to the measured standard deviation in T-jump magnitude. **(b)** Rate constants for AA association ($k_a$) and dissociation ($k_d$) as a function of $T_f$ determined from a two-state analysis of $1/\tau_{obs}$ (eq. 3). **(c)** Enthalpic ($\Delta H^{\ddagger}_{d,}$, $\Delta H^{\ddagger}_{a}$), entropic ($\Delta S^{\ddagger}_{d}$, $\Delta S^{\ddagger}_{a}$), and free energy (37 °C, $\Delta G^{\ddagger}_{d37}$, $\Delta G^{\ddagger}_{a37}$) barriers determined from fitting $k_d$ and $k_h$ temperature-trends to a Kramers-like model (eq. 3). Error bars indicate 95% confidence intervals propagated from the fits.

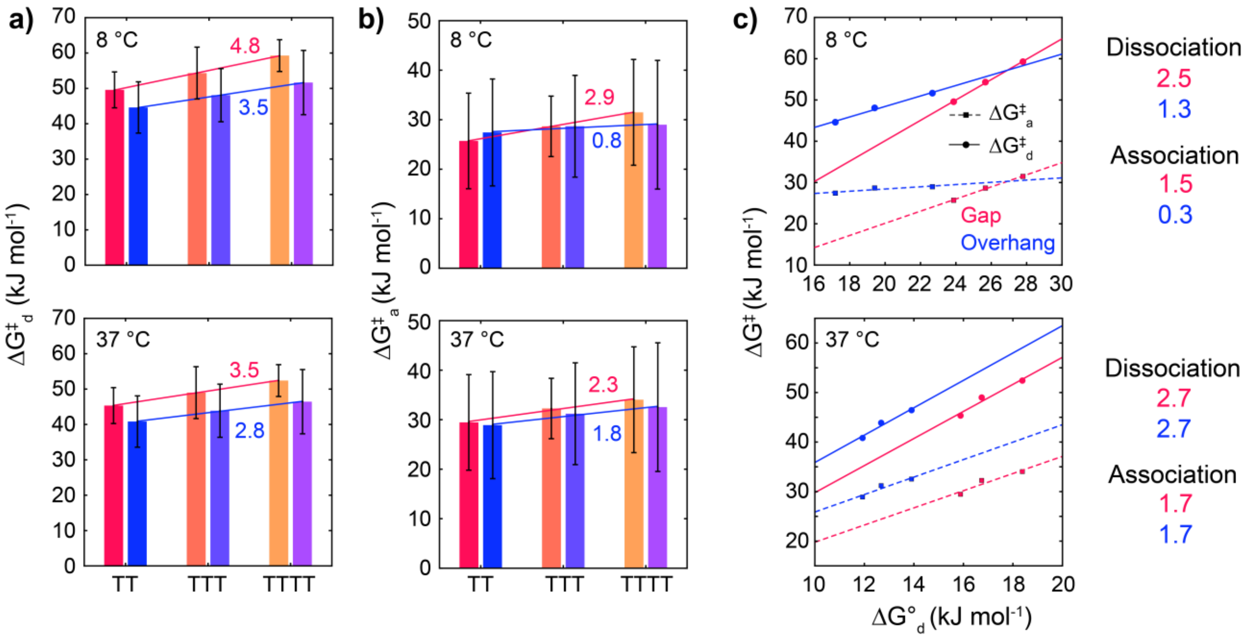


**Figure S28**. **Length-dependence of dehybridization and hybridization free energy barriers.** **(a)** $\Delta G_d^{\ddagger}$ at 8 and 37 °C for DNA gap (red-yellow) and overhang (blue-purple) sequences. Red and blue solid lines indicate linear fits to gap and overhang strand length trends, respectively. The slope of gap and overhang fit lines are listed above and below each line, respectively. Error bars are determined from 95% confidence intervals of fits to eq. 4. **(b)** Same plots for $\Delta G_a^{\ddagger}$ at 8 and 37 °C. **(c)** Correlation plots between $\Delta G_d^{\ddagger}, \Delta G_a^{\ddagger}$ and $\Delta G_d^{\circ}$ at 8 and 37 °C for DNA gap and overhang length series. Solid and dashed lines indicate linear fits with the respective slopes reported on the right.

### S5. Molecular dynamics simulations

#### S5.1 Structural comparison of gap single-strand regions and free single-strands

In addition to comparing the AA-gap complexes with canonical duplexes, we examine the structure of DNA gap-templates and their corresponding free single-strand oligonucleotides. It is important to note that DNA AMBER force fields are primarily parameterized for duplex structure, and therefore the structure of single-strands should be interpreted with caution. That being said, we find striking similarity between the TTg single-strand region and 5′-CTTC-3′ in terms of the distance between cytosine nucleobases on each side of the gap (C-to-C) and root-mean-square deviation (RMSD) relative to B-DNA (Fig. S29). The C-to-C distance distributions of each are peaked at 1 nm, identical to B-DNA, and the most probable RMSD is only 1 Å. Both the high C-to-C distance and low RMSD distribution regions are identical in TTg and 5′-CTTC-3′, suggesting that the single-strand segment is no more ordered than in free single-strand DNA. Overall, these single-strand regions are well stacked and resemble their structure in B-DNA. TTg instead contains a sub-population of bent configurations with greater RMSD and lower C-to-C distance. The bending is centered at the TT step where stacking is disrupted, but both CT and TC stacks remain intact and show increased overlap with the adjacent G:C base pair. Similar bending configurations have previously been proposed for 1- and 2-nucleotide DNA gaps.(35-43) Bending is found to a lesser and greater extent in TTTg and TTTTg, respectively. For TTTg, configurations with C-to-C distance of 1 nm and RMSD of 0.25 nm show disruption of the stacking between the central and 5′-T, and these thymines instead each stack with a flanking guanine (Fig S29). TTTTg adopts a highly bent configuration with a C-to-C distance of 0.9 nm where the stacking interactions between the 5′/3′ and central Ts are fully disrupted. However, configurations with a C-to-C distance of 1.3 nm and RMSD of 0.28 nm show similar alternative stacking to the minor state of TTTg. Overall, it is evident that the bendability and stacking arrangement of gapped DNA may depend on the length of the gap.

The structure and dynamics of the water network around the T nucleobases may differ in a gap or overhang template relative to free single-strands due to excluded volume and hydrophobic interactions from the flanking guanine nucleobases. We evaluated the radial distribution functions (RDF) between each thymine nucleobase and the surrounding water molecules and observe a reduction in water molecule density in the gap of TTg relative to CT2C (Fig. S30a). A similar reduction is found for TTTg and TTTTg at the thymine nucleobases at the ends of the gap (Fig. S30b,c). Beyond a mean T N3,O2,O4 to water oxygen (N3,O2,O4···O) distance of 0.8 nm, the reduction in water density primarily arises from excluded volume in TTg due to the flanking guanine nucleobases (Fig. S30d). However, there is also a reduction in probability at the 0.55 nm peak that corresponds to water molecules hydrogen bonded to thymine. The reduction of water molecules directly associated with thymine in TTg may still result from excluded volume interactions resulting from bending of the strand but may also be a consequence of the hydrophobic guanine nucleobases. Previous work employing MD simulations on single nucleotides gaps showed that residence times of water molecules within the gap cavity is slowed to similar values as those in minor groove of DNA.(38) Together, the gap appears to reduce the density of water molecules near thymine as well as constrain their motion, which may influence both $\Delta H_d^\circ$ and $\Delta S_d^\circ$ as these water molecules are displaced upon binding.

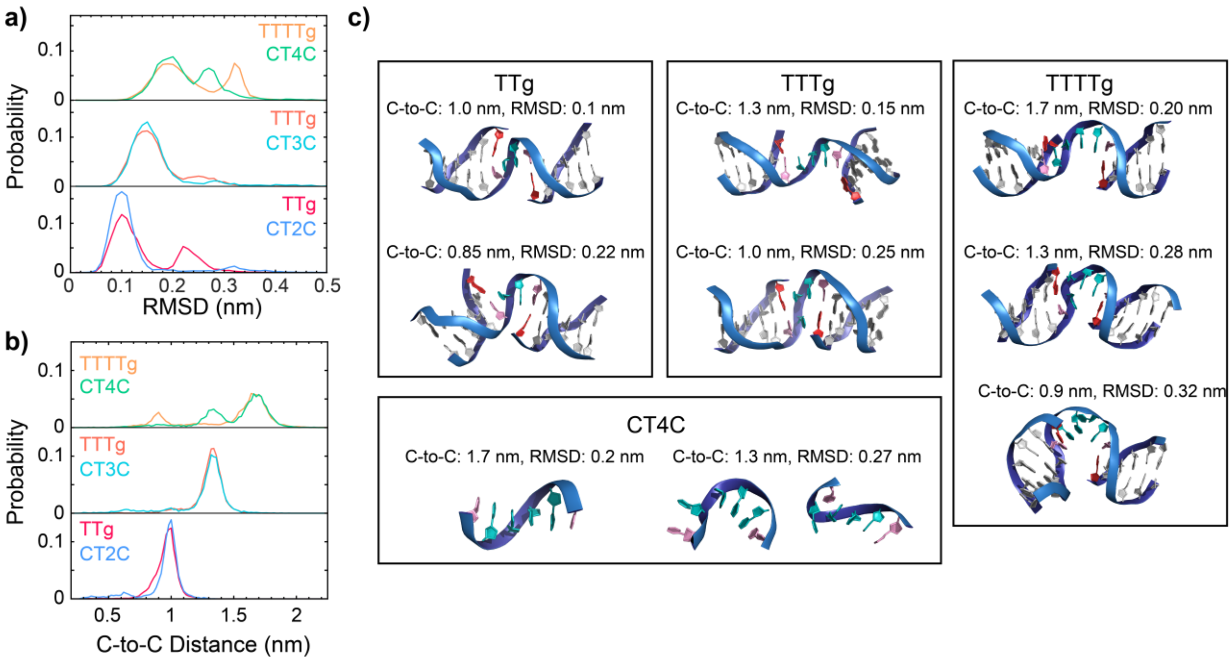


**Figure S29. C-to-C distance in gap templates and free single-strands from all-atom MD simulations. (a)** Probability distributions of root-mean-square deviation (RMSD) from B-DNA from 1 µs simulation time of gap templates (TTg, TTTg, TTTTg) and 0.2 µs simulation of free single-strand DNA (CT2C, CT3C, CT4C). **(b)** Probability distributions of center-of-mass distance between cytosine nucleobases adjacent to the $T_n$ regions (C-to-C) of DNA gap and $CT_nC$ sequences. Distributions are determined from 1 µs and 0.2 µs simulation time for gap templates and $CT_nC$ sequences, respectively, using the bsc1-AMBER force field at 37 °C. **(c)** Representative structures of TTg, TTTg, TTTTg, and CT4C at various C-to-C/RMSD values. There is significant overlap between the gap template and free single-strand distributions at the respective lowest RMSD or highest C-to-C distance maximum for each length, yet configurations with alternative stacking geometries or bending about the gap center are observed with increased RMSD and/or reduced C-to-C distance.

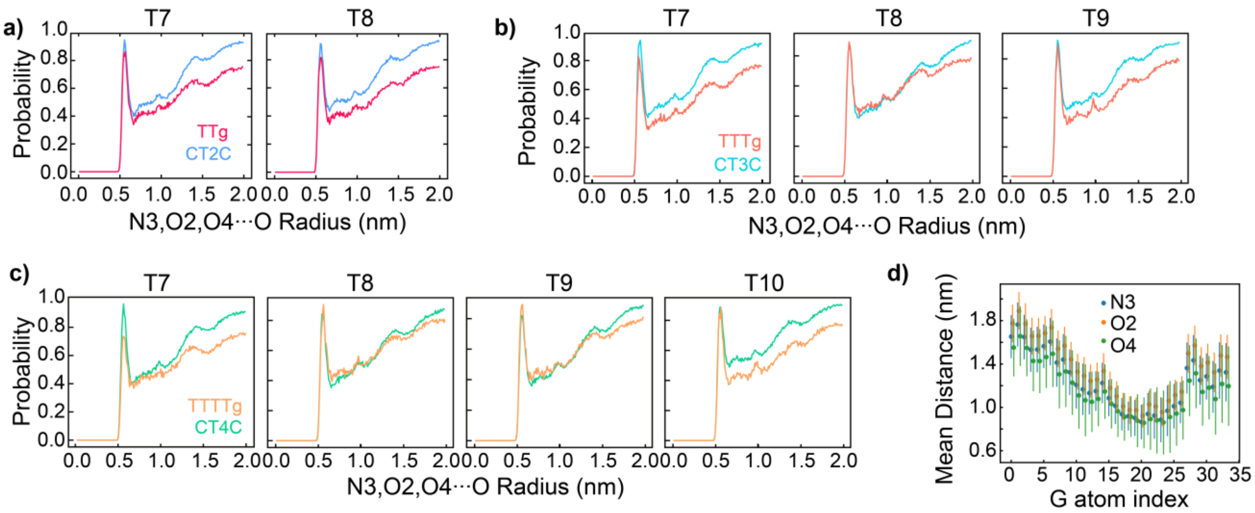


**Figure S30. Assessment of hydration in gap templates.** Radial distribution functions for the average over T N3, O2, and O4 distances to all water oxygen atoms. Each panel corresponds to a different thymine nucleobase in **(a)** TTg and C2TC, **(b)** TTTg and CT3C and **(c)** TTTTg and CT4C. **(d)** Mean distance between thymine N3, O2, and O4 atoms with each atom on adjacent G nucleotides in TTg. Error bars correspond to the standard deviation in distance across all simulation frames. Thymines at the edges of the gap exhibit a reduction in water probability from 0.5 – 0.7 nm relative to free single strands. The loss in water probability for >0.7 nm likely results from the excluded volume of adjacent guanine nucleobases.

#### S5.2 AA-gap complex and duplex structural parameters

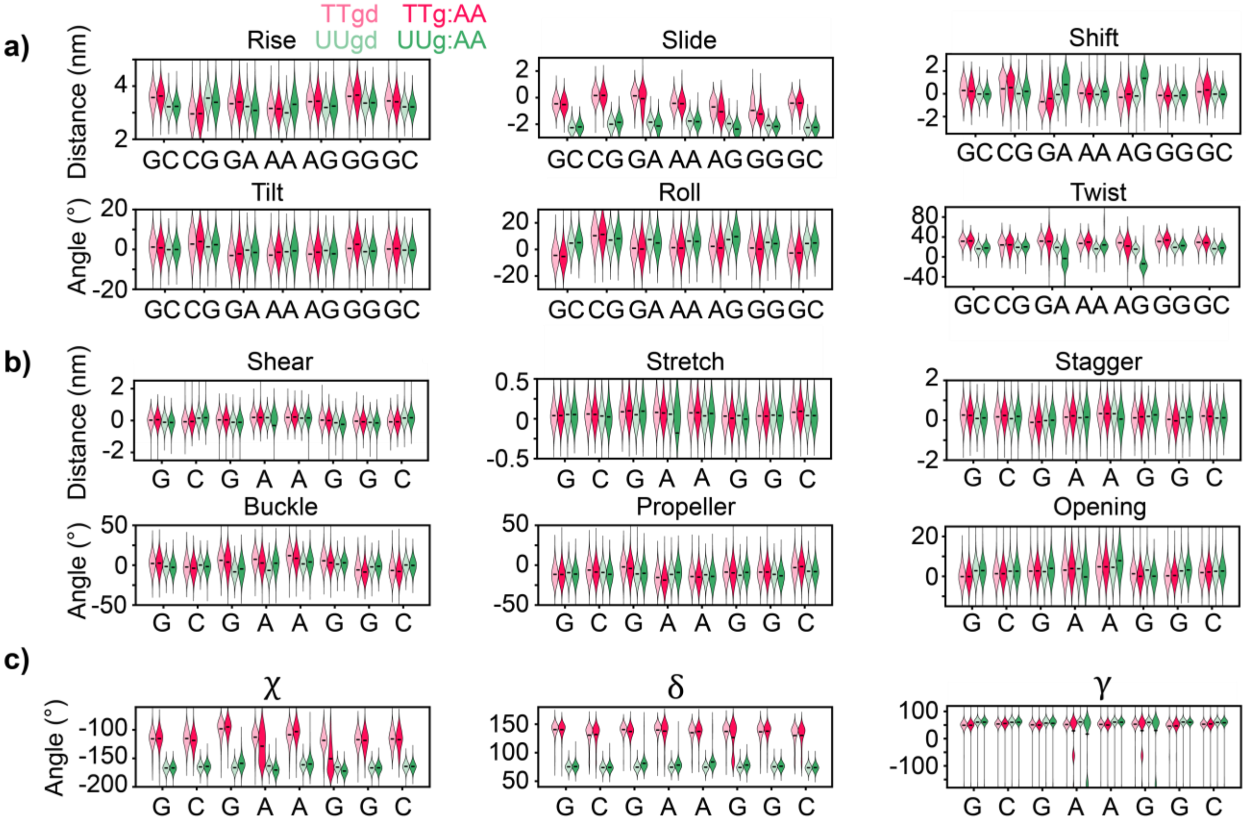


**Figure S31. Comparison of structural parameters in AA-gap complexes and duplex DNA and RNA.** Violin plots of **(a)** inter-base pair, **(b)** intra-base pair, and **(c)** select torsional angle distributions of central base pair steps and nucleotides in TTgd, TTg:AA, UUgd, and UUg:AA Black horizontal bars indicate the median value for each distribution determined from a Gaussian kernel estimation using Scott's rule.(44) Distributions were generated from 1.5 μs of simulation time for each sequence using the bsc1-AMBER force field for DNA sequences and DES-AMBER force field RNA sequences. Only trajectories of WCF base pair configurations are included the distributions. Relative to the canonical duplex, complexes show greatest differences in stacking geometry (twist, shift, slide) at the nick site. Larger differences in base pair (stretch, shear, opening, stagger) and stacking (shift, twist, rise) geometry are observed for RNA than DNA.

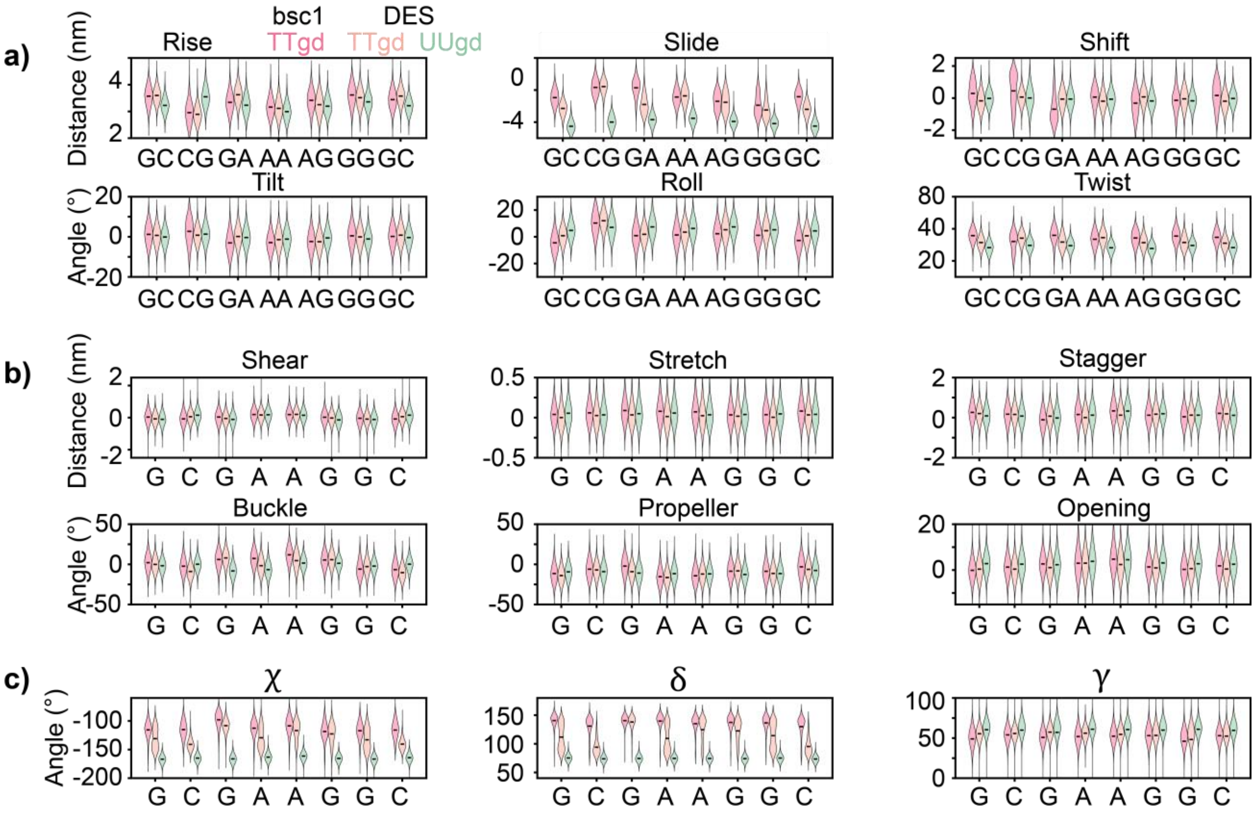


**Figure S32. Comparison of structural parameters obtained for duplex DNA using bsc1-AMBER and DES-AMBER force fields and duplex RNA using DES-AMBER.** Violin plots of **(a)** inter-base pair, **(b)** intra-base pair, and **(c)** select torsional angle distributions of central base pair steps and nucleotides in TTgd and UUgd. Black horizontal bars indicate the median value for each distribution determined from a Gaussian kernel estimation using Scott's rule.(44) Distributions were generated from 1.5 μs of simulation time for each sequence. Only trajectories of WCF base pair configurations are included the distributions. Substantial differences in mean stacking and backbone parameters are found between bsc1-AMBER and DES-AMBER force fields for DNA. Stacking (slide, twist, role) and backbone (χ, δ) parameters from DES-AMBER DNA simulations often have mean values in between those found from bsc1-AMBER DNA simulations and DES-AMBER RNA simulations.

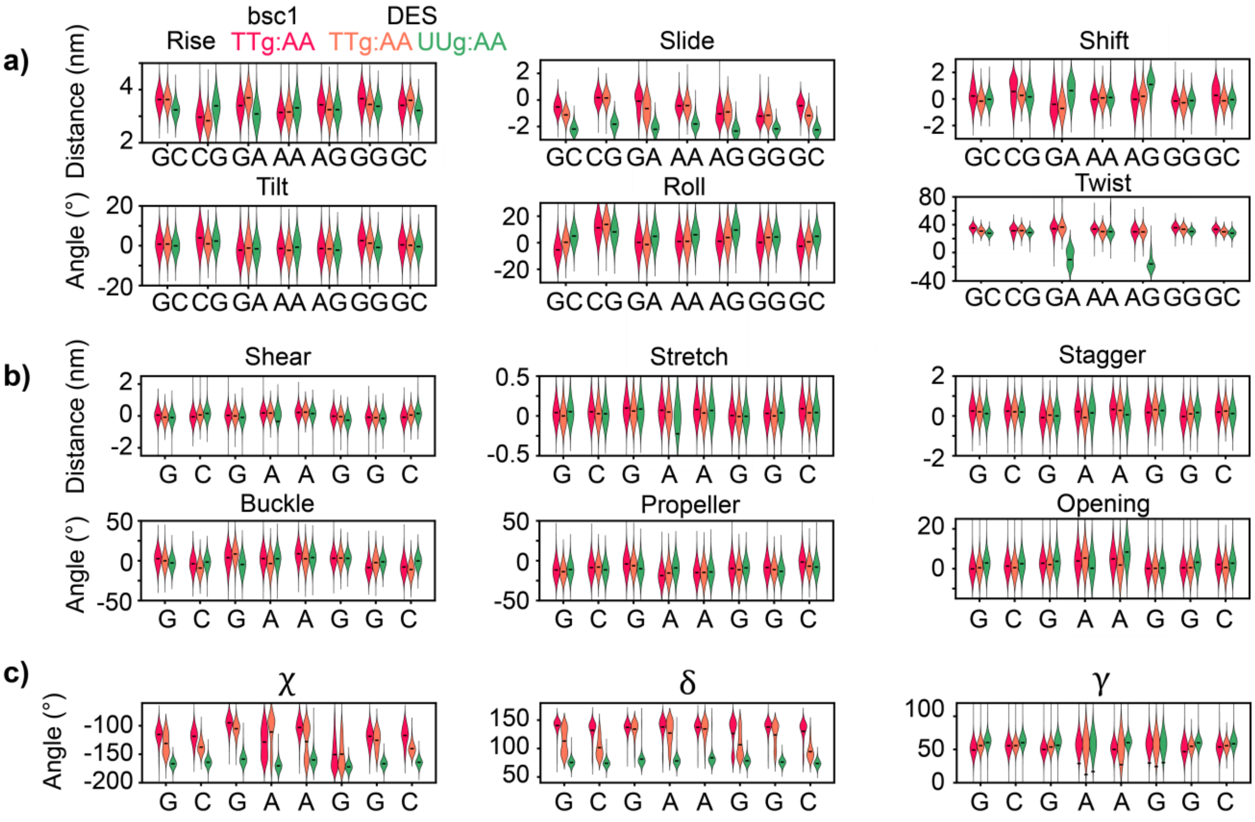


**Figure S33. Comparison of structural parameters obtained for TTg:AA using bsc1-AMBER and DES-AMBER force fields and UUg:AA using DES-AMBER.** Violin plots of **(a)** inter-base pair, **(b)** intra-base pair, and **(c)** select torsional angle distributions of central base pair steps and nucleotides in TTg:AA and UUg:AA. Black horizontal bars indicate the median value for each distribution determined from a Gaussian kernel estimation using Scott's rule.(44) Distributions were generated from 1.5 μs of simulation time for each sequence. Only trajectories of WCF base pair configurations are included the distributions. Substantial differences in the mean and distribution width of stacking (slide, shift) and backbone (χ, δ) parameters are found between bsc1-AMBER and DES-AMBER force fields for DNA.

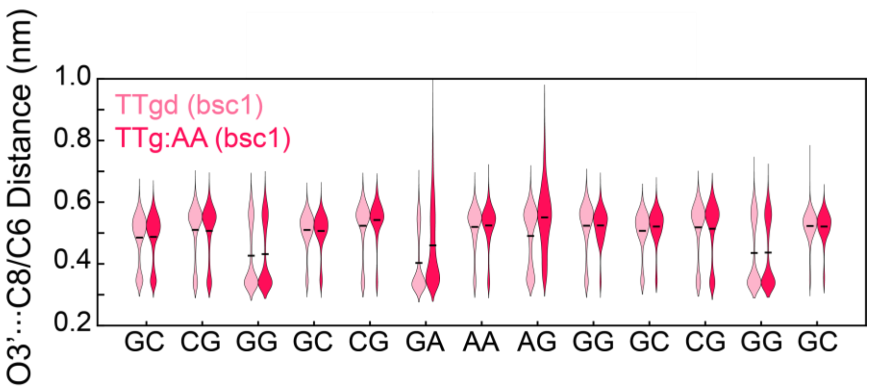

**Figure S34. Comparison of effective BI and BII population in DNA duplex and AA-gap complex.** Violin plots of the distributions for O3′···C8 distances in purine-purine or pyrimdine-purine steps and O3′···C6 distances in purine-pyrimidine steps. Black horizontal bars indicate the median value for each distribution determined from a Gaussian kernel estimation using Scott's rule.(44) TTgd shows two well-separated states for each step centered at ~0.5 nm and ~0.35 nm that correspond to BI and BII states, respectively. The greater backbone freedom of TTg:AA at GA and AG steps leads to a broadened distribution and greater median O3′···C8 distance than in TTgd.

##### S5.3 Calculation of relative WCF to anti-HG base pair stability with enhanced sampling

The On-the-fly Probability Enhanced Sampling (OPES)(45,46) was used to drive transitions and determine relative free energies between the canonical WCF dinucleotide configuration and an alternative anti-Hoogsteen (aHG) geometry. Like conventional Metadynamics,(47,48) OPES accelerates sampling along a pre-defined collective variable and leverages the deposited bias to recover equilibrium thermodynamic quantities. However, by reweighting on-the-fly, OPES optimizes the bias according to the ratio between target and Boltzmann probabilities leading to more efficient biasing and faster convergence. The biasing potential is expressed as follows

$$V(x) = -\frac{1}{k_B T} \ln\left(\frac{p^{tg}(x)}{p(x)}\right) \tag{S29}$$

Where $x$ is a set of collective variables, $p^{tg}$ is the target distribution, and $p$ is the Boltzmann distribution. As a biasing coordinate, we designed a one-dimensional collective variable $S$ that assigns a degree of similarity to the WCF state (near 1) and aHG state (near -1). $S$ is constructed as the difference between two contact map functions ($c_{WCF}$ , $c_{aHG}$ ).

$$S = c_{WCF} - c_{aHG} \tag{S30}$$

Where each contact map is the sum of four Q switching functions(49) corresponding to the four hydrogen bonds pairs that define the WCF (A N6 to T O4 and A N1 to T N3) and aHG states (A N7 to T N3 and A N6 to T O2).

$$c = \sum_{i}^{4} \frac{1}{4} \left( \frac{1}{1 + \exp\left[ \beta \left( r_i - \lambda r_i^0 \right) \right]} \right) \quad \text{(S31)}$$

We used a smoothing factor $\beta$ of 50, scaling factor $\lambda$ of 1.5, and a reference distance $r_i^0$ equal to the equilibrium hydrogen bond distance of each pair. To prevent the dinucleotide from fully dissociating or getting trapped in an unphysical configuration, we added a restraining potential to the center-of-mass separation between T and A nucleotides and to the center-of-mass separation within stacked GA and AG steps with 0.75 nm and 0.6 nm cutoffs, respectively. To prevent the dinucleotide from unstacking, we added a restraining potential on the eRMSD with respect to the crystal structure.(50) All additional biases were reweighted along with bias deposited along $S$ to ensure correct free energy comparisons.

Representative WCF and aHG configurations were identified from a 4 μs equilibrium DNA simulation and used as seeds for multi-walker OPES simulations (Fig. S35, 2 WCF seeds and 2 aHG seeds). Sampling was performed for ~400 ns per walker with bias factors ranging from 10 to 25. The collective variables and OPES biasing protocol were computed with PLUMED v2.8.(51,52) Reweighting was performed along $m_{WCF}$ and $m_{HG}$ which correspond to the mean distance between the four hydrogen bonds that define each configuration. Any configurations where $m_{WCF} <$ 0.32 nm was assigned to WCF basin and $m_{HG} <$ 0.32 nm was assigned to aHG. Relative Helmholtz free energies were calculated from the difference in probability densities between these two states as follows:

$$\Delta\Delta F_{WCF-aHG} = -RT \ln\left( \frac{p_{WCF}}{p_{aHG}} \right) \quad \text{(S32)}$$

Python scripts adapted from the OPES Github page (https://github.com/invemichele/opes) were used for reweighting.

WCF and aHG geometries have nearly the same population and stability at 37 °C with $\Delta\Delta F_{WCF-aHG}$= -2 - 0 kJ mol$^{-1}$ depending on the bias factor (Fig. S35b). Simulations with the DES-AMBER force field instead suggest that the WCF geometry is more stable ($\Delta\Delta F_{WCF-aHG}$ = -12 – -8 kJ mol$^{-1}$). The relative stability of aHG and WCF base pair geometries is not extensively characterized for DES-AMBER and may not be as reliable as bsc1-AMBER, but we use this result to illustrate that the aHG population is highly sensitive to force fields even within the AMBER family. The population of aHG base pairing is also significant for UUg:AA where similar results are obtained for using both DES-AMBER relative and bsc1-AMBER force fields. However, it should be noted that the aHG configuration was not observed in unbiased simulations of UUg:AA. Furthermore, because we designed and optimized our collective variable based on the aHG transition observed for TTg:AA, it may be suboptimal to drive aHG transitions for UUg:AA.

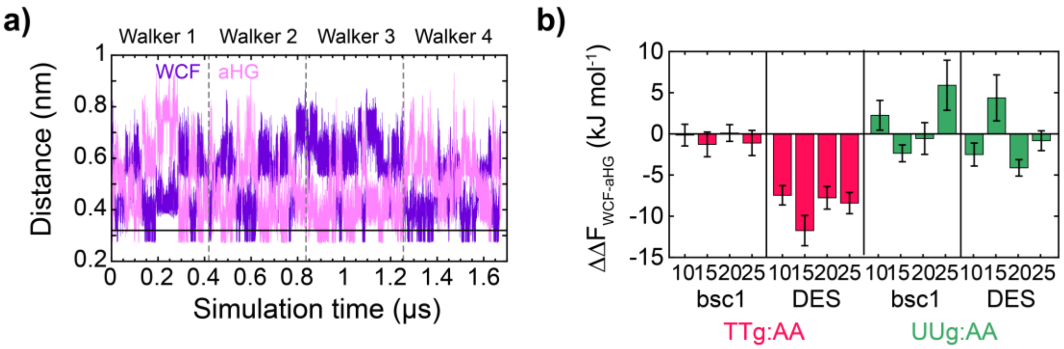


**Figure S35. Enhanced sampling of transition between WCF and aHG base pair configurations in AA-gap complexes. (a)** Trajectories of mean WCF (purple) and aHG (pink) distance shown for multiple walkers employing OPES. WCF and aHG distances are defined in the caption of Fig. 6. **(b)** Differences in Helmholtz free energy between WCF and aHG geometries ($\Delta\Delta F_{WCF-aHG}$) for TTg:AA and UUg:AA using the DES-AMBER and bsc1-AMBER force fields. Values of $\Delta\Delta F_{WCF-aHG}$ are shown using biasing factors of 10, 15, 20, 25. Error bars are determined from 10 block averages across trajectories.

#### S6. Estimation of diffusion-limited rate constant for AA binding to gap

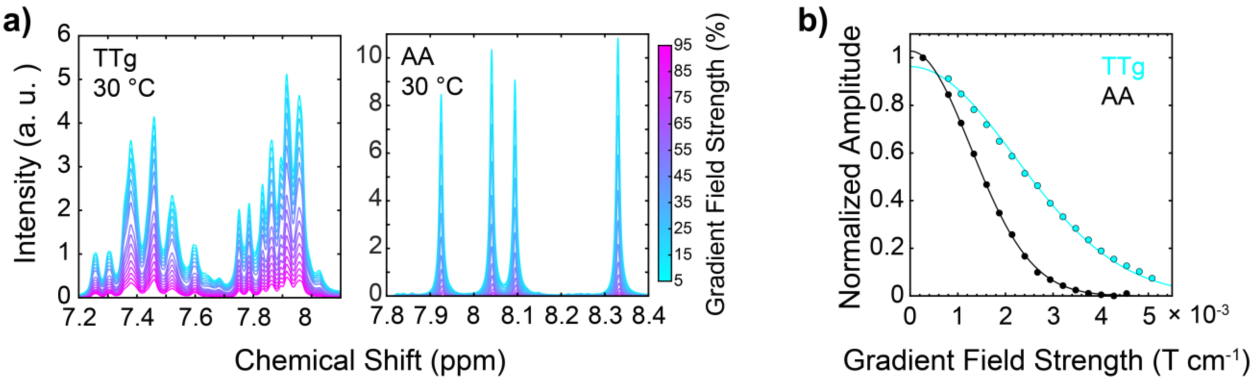


**Figure S36. Measuring the diffusion coefficient of TTg and AA with diffusion-ordered spectroscopy (DOSY) NMR spectroscopy. (a)** Aromatic $^1$H NMR spectra of (left) TTg and (right) free AA shown from 5 to 95% of the maximum magnetic gradient field strength (5.35 × 10$^{-3}$

T cm$^{-1}$). Spectra were acquired at 30 °C employing a diffusion time (Δ) of 100 ms and gradient pulse length (δ) of 4 ms. **(b)** Decay of normalized integrated signal over the aromatic $^1$H regions spectral regions shown in **(a)** as a function of gradient field strength. Solid lines correspond to fits to the Stejskal-Tanner equation (eq. S34). Translational diffusion coefficients of 92 ± 8 and 273 ± 10 μm$^2$ s$^{-1}$ were obtained for TTg and AA, respectively, where errors correspond to 95% confidence intervals from fits.

We estimate the diffusion-limited rate constant ($k_{diff}$) of AA-gap complex formation by treating TTg and AA as uniformly reactive spheres with distinct radii (*r*) and transitional diffusion coefficients (*D*).(53)

$$k_{diff} = 4\pi\left(D_{AA} + D_{TT_g}\right)\left(r_{AA} + r_{TT_g}\right) \quad \text{(S33)}$$

We use $r_{AA}$ = 4.7 Å, which comes from the experimentally measured length-scaling of radius of gyration ($R_g$) for polyA at [Na$^+$] = 500 mM.(54) We approximate $r_{TT_g}$ = 13.6 Å using the length-scaling of $R_g$ from the worm-like chain model of duplex DNA.(55) $D_{AA}$ and $D_{TT_g}$ were determined from diffusion-ordered spectroscopy (DOSY) NMR measurements of AA and TTg at 30 °C (Fig. S36). $^1$H NMR spectra were measured as a function of gradient magnetic field strength (*G*) from 5 to 95% of the maximum field strength (5.35 × 10$^{-3}$ T cm$^{-1}$). The signal decay (*S*) is fit to the Stejskal-Tanner equation to determine *D*.(56)

$$S(G) = S(0)\exp\left[-D\gamma^2 G^2 \delta^2 \left(\Delta - \delta/3\right)\right] \quad \text{(S34)}$$

The diffusion time (Δ) and gradient pulse length (δ) were set to 100 and 4 ms, respectively. γ corresponds to the gyromagnetic ratio of $^1$H. Fit values of $D_{AA}$ and $D_{TT_g}$ are 273 ± 10 and 92 ± 8 μm$^2$ s$^{-1}$, respectively.
